# Homotopic local-global parcellation of the human cerebral cortex from resting-state functional connectivity

**DOI:** 10.1101/2022.10.25.513788

**Authors:** Xiaoxuan Yan, Ru Kong, Aihuiping Xue, Qing Yang, Csaba Orban, Lijun An, Avram J. Holmes, Xing Qian, Jianzhong Chen, Xi-Nian Zuo, Juan Helen Zhou, Marielle V Fortier, Ai Peng Tan, Peter Gluckman, Yap Seng Chong, Michael J Meaney, Danilo Bzdok, Simon B. Eickhoff, B.T. Thomas Yeo

## Abstract

Resting-state fMRI is commonly used to derive brain parcellations, which are widely used for dimensionality reduction and interpreting human neuroscience studies. We previously developed a model that integrates local and global approaches for estimating areal-level cortical parcellations. The resulting local-global parcellations are often referred to as the Schaefer parcellations. However, the lack of homotopic correspondence between left and right Schaefer parcels has limited their use for brain lateralization studies. Here, we extend our previous model to derive homotopic areal-level parcellations. Using resting-fMRI and task-fMRI across diverse scanners, acquisition protocols, preprocessing and demographics, we show that the resulting homotopic parcellations are as homogeneous as the Schaefer parcellations, while being more homogeneous than five publicly available parcellations. Furthermore, weaker correlations between homotopic parcels are associated with greater lateralization in resting network organization, as well as lateralization in language and motor task activation. Finally, the homotopic parcellations agree with the boundaries of a number of cortical areas estimated from histology and visuotopic fMRI, while capturing sub-areal (e.g., somatotopic and visuotopic) features. Overall, these results suggest that the homotopic local-global parcellations represent neurobiologically meaningful subdivisions of the human cerebral cortex and will be a useful resource for future studies. Multi-resolution parcellations estimated from 1479 participants are publicly available (GITHUB_LINK).

## 1 Introduction

Information processing in the human brain is facilitated by the transformation of neural signals across cortical areas (Ungerleider and Desimone, 1986; Felleman and Van Essen, 1991). Therefore, accurate delineation of cortical areas is an important goal in systems neuroscience (Amunts and Zilles, 2015). Cortical areas are defined based on the principle that an area should exhibit distinct architectonics, topography, connectivity, and function (Kaas, 1987; Felleman and Van Essen, 1991; Eickhoff et al., 2018a). Traditionally, these criteria were evaluated with a broad range of invasive techniques. However, recent progress in non-invasive brain imaging techniques offer the opportunities to delineate cortical areas in vivo (Sereno et al., 1995; Cohen et al., 2008; Van Essen and Glasser 2014).

One popular non-invasive brain imaging technique is resting-state functional connectivity (RSFC). RSFC measures the synchrony of fMRI signals between brain regions, while a participant is “resting” in the scanner in the absence of any explicit task (Biswal et al., 1995). RSFC has been widely used to estimate a small number of (typically less than 20) large-scale brain networks (Damoiseaux et al., 2006; Calhoun et al., 2008; Smith et al., 2009; Power et al., 2011). RSFC has also been used to parcellate (or subdivide) the brain into hundreds of finer parcels (Craddock et al., 2012; Shen et al., 2013; Honnorat et al., 2015; Eickhoff et al., 2015). We refer to these finer parcellations as areal-level parcellations, since their boundaries are known to align with certain cortical areas (Wig et al., 2014; Gordon et al., 2016; Schaefer et al., 2018). However, cortical areas are internally heterogeneous (Kaas 1987), so most RSFC parcellations also capture sub-areal features, such as visual eccentricity and somatomotor representations of different body parts (Yeo et al., 2011; Gordon et al., 2016; Schaefer et al., 2018). This sub-areal characteristic can be advantageous, for instance when modeling a behavioral task involving button presses, it might be useful for hand and tongue motor regions to be represented by different parcels.

We previously developed a gradient-weighted Markov Random Field (gwMRF) model for estimating areal-level RSFC parcellations (Schaefer et al., 2018). The gwMRF model integrates two popular approaches for estimating brain parcellations. The local gradient approach (Hirose et al., 2012; Wig et al., 2014; Laumann et al., 2015; Xu et al., 2016) detects abrupt changes in RSFC patterns between adjacent spatial locations in the cerebral cortex. The locations of these abrupt changes often correspond to known cortical areal boundaries (Wig et al., 2014; Gordon et al., 2016). On the other hand, the global similarity approach clusters the brain into parcels with similar RSFC patterns or resting-fMRI time courses (van den Heuvel et al., 2008; Power et al., 2011; Yeo et al., 2011; Ryali et al., 2013). By construction, the resulting parcels are more homogeneous (than those obtained with the local gradient approach), which is useful for dimensionality reduction of fMRI data. By combining both local gradient and global similarity approaches, the gwMRF parcellations agreed well with the boundaries of certain cortical areas defined using histology and visuotopic fMRI (Schaefer et al., 2018). The resulting gwMRF parcels were also more homogeneous than other parcellations evaluated using task-fMRI and resting-fMRI (Schaefer et al., 2018).

However, there is no homotopic correspondence between left and right gwMRF parcels, which limits their use for lateralization studies. Histological studies suggest the presence of homotopic pairs of cortical areas in roughly spatially homotopic locations across the hemispheres (Amunts et al., 2020). Homotopic areas do not necessarily have the same function, e.g., language processing is lateralized towards the left hemisphere in most participants (Szaflarski et al., 2006; Hartwigsen et al., 2021; Malik-Moraleda et al., 2022).

Despite functional lateralization, most higher-level cognitive functions are supported by distributed networks across the two hemispheres, enabled by inter-hemispheric commissural fibers (e.g., corpus callosum) and subcortical relays (Middleton and Strick, 2020; Schmahmann et al., 2008; Jones 2012). Direct and indirect connections between the two hemispheres in turn lead to strong homotopic RSFC observed in both electrophysiological (Duffy et al., 1996) and fMRI (Lowe et al., 1998; Salvador et al., 2005) studies. Spatial variation in homotopic RSFC has been observed along the functional hierarchy with sensory-motor regions exhibiting stronger homotopic RSFC than association regions (Stark et al., 2008). Within the sensory-motor cortex, tongue regions exhibit stronger homotopic RSFC than hand and foot regions (Yeo et al., 2011), consistent with nonhuman primate studies showing that the representations of midline structures in S1 and M1 (e.g., face) have denser callosal connections than those of distal limbs (e.g., hand and foot; Pandya and Vignolo 1971; Jones and Wise 1977; Killackey et al., 1983; Gould et al., 1986).

In this study, we extend the gwMRF model to derive homotopic areal-level parcellations, in which pairs of parcels exist in approximately spatially homotopic locations across the cerebral cortical hemispheres. Homotopic parcellations have been derived by manual delineation of parcellation boundaries (Glasser et al., 2016) or post hoc grouping of parcel pairs (Joliot et al., 2015). By contrast, our homotopic Markov Random Field (hMRF) approach is fully automated and seeks to preserve the desirable properties of the original gwMRF approach. We compared the hMRF parcellations with six publicly available parcellations (Craddock et al., 2012; Shen et al., 2013; Joliot et al., 2015; Glasser et al., 2016; Gordon et al., 2016; Schaefer et al., 2018) using multiple datasets. Based on task-fMRI and resting-fMRI, the hMRF parcels were as homogeneous as gwMRF parcels, while being more homogeneous than the other five parcellations. Compared with other fully automated approaches, the hMRF parcellations exhibited similar alignment with architectonic and visuotopic areal boundaries. Overall, while the hMRF parcellations do not correspond to cortical areas, they nevertheless provide a meaningful subdivision of the cerebral cortex that is useful for future analyses.

## 2 Material and Methods

### 2.1 Overview

A homotopic Markov Random Field (hMRF) parcellation procedure was developed and applied to resting-fMRI from the GSP dataset. Using data collected from multiple scanners, acquisition protocols and preprocessing procedures, the estimated homotopic parcellations were compared with six previously published resting-fMRI parcellations (Craddock et al., 2012; Shen et al., 2013; Joliot et al., 2015; Glasser et al., 2016; Gordon et al., 2016; Schaefer et al., 2018). Of these six parcellations, two of them were homotopic (Joliot et al., 2015; Glasser et al., 2016). A final set of hMRF parcellations at various resolutions were estimated from the full GSP dataset and further characterized.

### 2.2 fMRI Datasets

#### 2.2.1 GSP dataset

The GSP dataset consisted of resting-fMRI data from 1489 young adults aged between 18 to 35 years old (Holmes et al., 2015). All imaging data were collected on matched 3T Tim Trio scanners (Siemens Healthcare) at Harvard University and Massachusetts General Hospital using the vendor-supplied 12-channel phased-array head coil. For each participant, one or two resting-fMRI runs were acquired. Out of the total 1489 participants, 1083 participants had two runs and 406 participants had one run. Each run was acquired in 3 mm isotropic resolution with a TR of 3.0 s and lasted for 6 min and 12 s. The structural data consisted of one 1.2 mm isotropic scan for each participant.

Detailed information about the resting-fMRI preprocessing can be found elsewhere (Li et al., 2019). Here we provide the broad outlines of the processing steps, which utilized a mixture of FreeSurfer 5.3.0 (Fischl, 2012), FSL 5.0.8 (Jenkinson et al., 2012) and in-house Matlab functions. The processing steps were as follows. (1) First four frames were removed. (2) Slice time correction was applied. (3) Motion correction and censoring of outlier volumes were performed. Volumes with framewise displacement (FD) > 0.2 mm or root-mean-square of voxel-wise differentiated signal (DVARS) > 50, along with one volume before and two volumes after, were marked as outliers. Uncensored segments of data lasting fewer than five contiguous volumes were also labeled as censored frames. BOLD runs with more than half of the volumes labeled as censored frames were removed. (4) Alignment with structural images with boundary-based registration was performed (Greve and Fischl, 2009). (5) White matter and ventricular signals, whole brain signal, six motion parameters, and their temporal derivatives were regressed from the fMRI data. Outlier frames from step 3 were excluded when computing the regression coefficients. (6) Censored frames were interpolated with Lomb-Scargle periodogram (Power et al., 2014), (7) Bandpass filtering (0.009Hz < f < 0.08Hz) was performed. (8) Data was projected to the FreeSurfer fsaverage6 space and smoothed by a 6mm full-width half-maximum (FWHM) kernel. Only participants with at least one run remaining (N = 1479) were considered. The 1479 participants were further divided with no participant overlap between training (N = 740) and test (N = 739) sets. Age, sex, handedness and number of available runs were balanced across the training and test sets (Table 1). The GSP training set was used to estimate hMRF parcellations, while the GSP test set was used to evaluate the parcellations (Sections 2.4 and 2.5). The full dataset (N = 1479) was also used to estimate a final set of hMRF parcellations at different resolution (Section 2.6).

**Table 1.** Training and test sets were balanced across sex, handedness, age, and number of runs. There was no participant overlap across training and test sets.

| | % Female | % Right-handers | Age (mean ± std) | # runs (mean ± std) |
| --- | --- | --- | --- | --- |
| Training Set | 58.1 | 92.4 | 21.5 ± 2.9 | 1.7 ± 0.4 |
| Test Set | 58.3 | 92.3 | 21.5 ± 2.9 | 1.7 ± 0.4 |

#### 2.2.2 HCP dataset

For the purpose of evaluation (Section 2.4), we considered resting-fMRI data from the HCP S1200 release. Details about the acquisition protocol and minimal processing for the HCP data can be found elsewhere (Glasser et al., 2013; Van Essen et al., 2012b). Briefly, each participant has 4 resting-fMRI runs acquired from a custom-made Skyra scanner. Each run was acquired in 2 mm isotropic resolution with a TR of 0.72 s and lasted for 14 min and 33 s. The structural data consisted of one 0.7 mm isotropic scan for each participant.

We utilized ICA-FIX resting-fMRI data in MNI152 space, which have been denoised with ICA-FIX (Griffanti et al., 2014; Salimi-Khorshidi et al., 2014) and smoothed by a Gaussian kernel with 6 mm FWHM. We also considered MSMAII ICA-FIX resting-fMRI data, which were projected to fs_LR32k surface space (Van Essen et al., 2012b), smoothed by a Gaussian kernel with 2 mm FWHM and aligned with MSMAII (Robinson et al., 2014).

To further remove head motion-related artifacts (Burgess et al., 2016; Siegel et al., 2017), additional censoring was performed (volumes with FD > 0.2 mm or DVARS > 50 were marked as outliers) and runs with more than 50% censored frames were removed. Only participants with at least one resting-fMRI run remaining (N = 1030) were considered.

Furthermore, for the purpose of evaluation, we utilized task contrasts released by the HCP in fs_LR32k surface space. Details of the task contrasts can be found in the HCP documentation and publications (Barch et al., 2013). Briefly, there were seven tasks: social cognition, motor, gambling, working memory, language processing, emotional processing, and relational processing. For each task, there were multiple task contrasts released by the HCP in fsLR surface space. We used all independent task contrasts. The subset of HCP participants with available contrasts for all tasks was considered (N = 956).

#### 2.2.3 ABCD dataset

The GSP test set (Section 2.2.1) and HCP dataset (Section 2.2.2) allowed us to evaluate parcellation quality in young adults in MNI152, fsaverage and fs_LR32k spaces. We also considered 9- to 10-year-old children from the Adolescent Brain Cognitive Development (ABCD) dataset to evaluate parcellation quality in children. Details of the ABCD dataset can be found elsewhere (Casey et al., 2018). Briefly, each participant has 4 resting-fMRI runs acquired from Philips, GE or Siemens scanners. Consistent with our previous study (Chen et al., 2022), individuals from Philips scanners were excluded due to incorrect preprocessing.

Each resting-fMRI run was acquired in 2.4 mm isotropic resolution with a TR of 0.8 s and lasted for 5 min. The structural data consisted of one 1 mm isotropic scan for each participant.

We considered 2262 participants from our previous study (Chen et al., 2022). Briefly, we utilized minimally preprocessed fMRI data released by the ABCD study (Hagler et al., 2019). The minimally processed data were further preprocessed with the following steps. (1) Functional images were aligned to T1 images using boundary-based registration (Greve & Fischl, 2009). (2) Respiratory pseudo-motion was removed with a bandstop filter of 0.31- 0.43Hz (Fair et al., 2020). (3) Frames with FD > 0.3 mm or DVARS > 50, along with one volume before and two volumes after, were marked as outliers and subsequently censored.

Uncensored segments of data consisting of less than 5 frames were also censored. Runs with more than 50% censored frames were not considered. (4) Global, white matter and ventricular signals, 6 motion parameters, and their temporal derivatives were regressed from the fMRI data. Censored frames were not considered when computing the regression coefficients. (5) Censored frames were interpolated with Lomb-Scargle periodogram (Power et al., 2014). (6) Bandpass filtering (0.009Hz – 0.08Hz) was applied. (7) Finally, the data was projected onto FreeSurfer fsaverage6 surface space and smoothed using a 6 mm full-width half maximum kernel.

The ABCD study also provided task-fMRI data from the N-back, monetary incentive delay (MID), and stop signal tasks (SST). There were 2 fMRI runs for each task. Each run was acquired in 2.4 mm isotropic resolution with a TR of 0.8 s. Each MID run was 322.4- secs long. Each N-back run was 289.6-secs long. Each SST run was 349.6-secs long. Preprocessing of the task data was similar to the resting-fMRI data with some notable differences. Briefly, we utilized minimally preprocessed fMRI data released by the ABCD study (Hagler et al., 2019). The minimally processed data were further preprocessed with the following steps. (1) Functional images were aligned to T1 images using boundary-based registration (Greve & Fischl, 2009). (2) Respiratory pseudo-motion was removed with a bandstop filter of 0.31-0.43Hz (Fair et al., 2020). (3) Frames with FD > 0.3 mm or DVARS > 50, along with one volume before and two volumes after, were marked as outliers. Segments of data consisting of less than 5 non-outlier frames were also marked as outliers. Runs with more than 50% outlier frames were not considered. (4) 6 motion parameters, and their temporal derivatives were regressed from the fMRI data. Outlier frames were not used to compute regression coefficients. (5) Finally, the data was projected onto FreeSurfer fsaverage6 surface space and smoothed using a 6 mm full-width half maximum kernel. To estimate the task-related activation, we fitted general linear model to the preprocessed task-fMRI data using AFNI’s 3dDevonvolve (Cox, 1996). We utilized task regressors provided by ABCD (Casey et al., 2018), comprising 15 task conditions from MID, 9 conditions from N-back and 8 conditions from SST. The resulting activation z-scores were finally projected to the fsaverage6 surface mesh.

#### 2.2.4 GUSTO dataset

The previous evaluation datasets comprised mostly participants from North America. Given concerns about cross-ethnicity generalization failures in neuroimaging studies (Li et al., 2022), we also considered resting-fMRI data from 7.5-year-old children in the Growing Up in Singapore Towards Healthy Outcomes (GUSTO) dataset (Soh et al., 2014). Each participant has 1 resting-fMRI run collected on a Siemens Prisma scanner. Each run was acquired in 3 mm isotropic resolution with a TR of 2.62 s and lasted for 5.24 min. The structural data consisted of one 1 mm isotropic scan for each participant.

The processing steps are as follows. (1) First four frames were removed. (2) Slice time correction was applied. (3) Motion correction was performed, and frames with FD > 0.6 mm or DVARS > 80 were marked as outliers. (4) Alignment with structural images with boundary-based registration was performed (Greve and Fischl, 2009). (5) White matter and ventricular signals, whole brain signal, six motion parameters, and their temporal derivatives were regressed from the fMRI data. Outlier frames from step 3 were excluded when computing the regression coefficients (6) AFNI despiking was applied to the data. (7) The resulting time courses were bandpass filtered (0.009Hz < f < 0.08Hz). (8) The fMRI data was projected to the FreeSurfer fsaverage6 space. (9) Surface data were smoothed by a 6mm full-width half-maximum (FWHM) kernel. Participants with poor T1-fMRI alignment (based on visual QC) was excluded. Participants with less than 4 minutes of non-outlier frames were also excluded, yielding a final set of 393 participants.

### 2.3 hMRF parcellation procedure

A typical MRF model is defined by an objective function comprising several competing terms encoding the ideal properties of a segmentation or parcellation. The homotopic Markov Random Field (hMRF) model is the same as the gwMRF model (Schaefer et al., 2018) except for two additional terms to encourage homotopic parcels.

The gwMRF objective function comprises three terms, which encode the global connectivity similarity objective, the local gradient objective, and a spatial contiguity constraint. The global connectivity similarity term encourages brain locations with similar preprocessed fMRI time courses to be assigned to the same parcel. The local gradient term penalizes adjacent brain locations with different parcellation labels. This penalty is relaxed in the presence of strong local RSFC gradients (hence the name “gradient weighted”). With only the first two terms, the resulting parcellation will contain spatially distributed parcels due to long-range RSFC. Therefore, the spatial contiguity term encourages brain locations within a parcel to be near to the parcel center. However, the spatial contiguity term has the side effect of encouraging rounder parcels. Therefore, care was taken to ensure the spatial contiguity term was just sufficiently strong to obtain spatially contiguous parcels. More details about the gwMRF model can be found elsewhere (Schaefer et al., 2018). For completeness, Supplementary Methods S1 provides details about the three gwMRF terms.

Our proposed hMRF objective function has a fourth homotopic constraint term that encourages homotopic pairs of vertices to be assigned to homotopic pairs of parcels. Homotopic vertex pairs were defined as follows. The left and right fsaverage6 hemispheres were first registered using the Freesurfer’s “mris_left_right_register” function. After registration, the closest right hemisphere vertex was found for each left hemisphere vertex.

Similarly, the closest left hemisphere vertex was found for each right hemisphere vertex. Here, the closest vertex was defined based on geodesic distance on the spherical surface mesh. A pair of left and right hemisphere vertices was considered homotopic if they were each other’s closest neighbor. Without loss of generality, the *p*-th parcels on the left and right hemispheres are assumed to be homotopic pair of parcels. For example, in the case of the 400-region hMRF parcellation, we assume that parcels 1 to 200 are in the left hemisphere and parcels 201 to 400 are in the right hemisphere. Furthermore, we assume that parcels 1 and 201 are homotopic, parcels 2 and 202 are homotopic, etc. The homotopic constraint imposes a penalty if a pair of homotopic vertices is assigned to non-homotopic pairs of parcels.

Because of the strong homotopic correlations between the two hemispheres, a fifth “bookkeeping” term is necessary to prevent any parcel from spanning across both hemispheres. Using the example of the 400-region hMRF parcellation, this bookkeeping term imposes a very large penalty if any right hemisphere vertex is assigned to parcels 1 to 200 or if any left hemisphere vertex is assigned to parcels 201 to 400.

Mathematical details of the hMRF model are found in Supplementary Methods S1 and S2. Tradeoffs among different terms of the hMRF model are governed by a set of hyperparameters. For a fixed set of hyperparameters, we used the alpha expansion algorithm to estimate the parcellation labels within a maximum-a-posteriori estimation framework (Delong et al., 2010), following our previous study (Schaefer et al., 2018). Details of the estimation procedure is found in Supplementary Methods S3 and S4. Supplementary Methods S5 provides details on how the hyperparameters are set in the following analyses.

### 2.4 Quantitative Evaluation Measures

Evaluating the quality of a human cerebral cortical parcellation is difficult due to a lack of ground truth. Motivated by the definition of a cortical area (Felleman and Van Essen, 1991), we considered four evaluation metrics utilized in previous parcellation studies (Glasser et al., 2016; Gordon et al., 2016; Gordon et al., 2017; Schaefer et al., 2018): architectonic alignment, visuotopic alignment, task functional inhomogeneity and resting-state connectional homogeneity. These evaluation metrics were utilized to compare hMRF parcellations with other public parcellations after controlling for the number of parcels (Section 2.5). A good parcellation should align well with known cytoarchitectonic and visuotopic boundaries, as well as have homogeneous function and connectivity.

#### 2.4.1 Architectonic boundaries

Ten human architectonic areas were considered: 1, 2, 3 (areas 3a and 3b combined), 4 (areas 4a and 4p combined), 6, 17, 18 hOc5, 44, and 45 (Geyer et al., 1996, 1999, 2000, 2001; Amunts et al., 1999, 2000, 2004; Geyer 2004; Malikovic et al., 2007). These histological areas were projected to the fsLR surface template by Van Essen et al (2012a) based on Fischl et al (2008). To evaluate the alignment between these histological areas and the boundaries of a given parcellation, averaged geodesic distances were computed. In short, for each boundary vertex of an architectonic area, the geodesic distance to the nearest parcellation boundary was computed, then averaged across all boundary vertices of the area. Lower geodesic distance indicated better alignment between histological and parcellation boundaries. When comparing two parcellations, paired-sample t-test of the geodesic distances from both hemispheres was used (degrees of freedom or dof = 19).

#### 2.4.2 Visuotopic boundaries

Eighteen visuotopic areas were considered (Abdollahi et al., 2014). The areas were obtained by averaging individual fMRI visuotopic mapping in fsLR surface space after multimodal alignment (Abdollahi et al., 2014). Each visuotopic area generally occupied half of its true extent since it was difficult to stimulate the peripheral visual field (Hinds et al., 2009). Therefore, we only considered visuotopic boundaries between adjacent areas, yielding 46 pairs of adjacent areas across both hemispheres. The agreement between visuotopic areal boundaries and parcellation boundaries was quantified by computing the geodesic distance. For each visuotopic boundary vertex, the geodesic distance to the nearest parcellation boundary was computed. The geodesic distances were averaged across all boundary vertices of each pair of adjacent areas. Lower geodesic distance indicated better alignment between visuotopic and parcellation boundaries. When comparing two parcellations, paired-sample t-test of the geodesic distances was used (dof = 45).

#### 2.4.3 Task functional inhomogeneity

Given a task activation contrast, the task functional inhomogeneity of a parcel was defined as the standard deviation (SD) of activation z-values across vertices within the parcel. The task functional inhomogeneities were combined across all parcels, yielding an overall task functional inhomogeneity metric (Gordon et al., 2017; Schaefer et al., 2018):

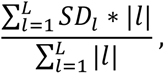

where *SD_l_* was the task functional inhomogeneity of the *l*-th parcel, |*l*| indicated the number of vertices in parcel *l* and *L* was the total number of parcels. Therefore, larger parcels were weighted more in the overall task functional inhomogeneity metric. A lower task inhomogeneity metric indicated that activation within each parcel was more uniform, suggesting higher parcellation quality.

In this study, we utilized task contrasts from the HCP and ABCD datasets (Sections 2.2.2 and 2.2.3), which were in fsLR and fsaverage space respectively. For a given parcellation, task inhomogeneity was computed for each task contrast and each participant. Task inhomogeneities were then averaged across all contrasts within each task. When comparing two parcellations for a given dataset, task inhomogeneities were further averaged across all tasks before applying the paired-sample t-test (dof = 955 for the HCP dataset and dof = 2261 for the ABCD dataset).

#### 2.4.4 Resting-state connectional homogeneity

RSFC homogeneity was defined as the averaged Pearson’s moment-product correlations between resting-fMRI time courses of all pairs of vertices (or voxels) within a given parcel. The correlations were then averaged across all parcels, while accounting for parcel size (Schaefer et al 2018; Kong et al 2021a):

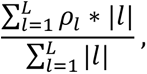

where *ρ*_*l*_ was the averaged correlation within parcel *l*, |*l*| indicated the number of vertices in parcel *l* and *L* was the total number of parcels. A higher resting-state homogeneity meant that vertices within a parcel shared more similar time courses. Therefore, a higher resting-state homogeneity indicated better parcellation quality.

The above normalization by parcel size is important to avoid situations in which a high homogeneity measure can be obtained for a meaningless parcellation. For example, imagine a 400-region parcellation comprising 399 parcels (containing 1 voxel each) and 1 huge parcel (containing the remaining voxels). Then the 399 small parcels have a resting- homogeneity of 1, while the gigantic parcel has a resting-homogeneity of about zero. If we simply average the homogeneity across the 400 parcels without accounting for parcel size, the mean homogeneity would be almost 1 (almost perfect homogeneity), while in reality, this 400-region parcellation is meaningless. Therefore, the normalization by parcel size was performed for both task functional inhomogeneity and resting-state homogeneity.

In the current study, resting homogeneity was computed using (1) GSP test set in fsaverage space (N = 739), (2) HCP dataset in MNI152 volumetric space (N = 1030), (3) HCP dataset in fsLR space (N = 1030), (4) ABCD dataset in fsaverage space (N = 2262) and (5) GUSTO dataset in fsaverage space (N = 393). Therefore, the datasets spanned across multiple volume and surface coordinate systems. The GSP dataset consisted of participants aged 18 to 35, the HCP datasets consisted of participants aged 22 to 35, while the ABCD and GUSTO datasets comprised children aged 7 to 10 years old. Furthermore, while the GSP, HCP and ABCD datasets comprised largely white participants, the GUSTO comprised East Asian participants. Finally, the HCP dataset was processed using ICA-FIX, while the remaining datasets were processed with whole brain signal regression. The wide variety of datasets allowed us to evaluate the generalizability of the hMRF parcellations to new datasets in terms of resting-homogeneity.

To compare two parcellations for a given dataset, resting-state homogeneity was computed for each participant before a paired-sample t-test was performed (dof = 738 for GSP, dof = 1029 for HCP, dof = 2261 for ABCD and dof = 392 for GUSTO).

A cortical parcellation with more parcels will have smaller parcels, so will generally perform better on the resting homogeneity and task inhomogeneity metrics. A cortical parcellation with more parcels will also generally have lower architectonic and visuotopic geodesic distances. The reason is that more parcels will lead to more parcellation boundary vertices. Therefore, it is more likely that histological (or visuotopic) boundaries will lie closer to some parcellation boundary vertices. Overall, when assessing the above evaluation metrics, it is critical to control for the number of parcels (see next section).

#### 2.4.5 Homotopic resting-state functional connectivity

RSFC was computed between homotopic pairs of parcels. More specifically, for each parcel pair, every vertex from one parcel was correlated with every vertex from the homotopic parcel. These correlations were averaged within parcel pairs, then averaged across all parcel pairs, weighted by the number of vertices within each parcel pair. The resulting resting-state homotopic connectivity was compared between the hMRF parcellations and the two homotopic parcellations (AICHA and Glasser) using the same datasets as in section 2.4.4. The assumption was that a better homotopic parcellation should exhibit stronger homotopic RSFC

### 2.5 Comparison with other parcellations

To evaluate the hMRF parcellations, we utilized the GSP training set (N = 740) to estimate a set of hMRF parcellations with different resolutions in fsaverage6 surface space. Details regarding the estimation procedure can be found in the Supplementary Methods. The resulting hMRF parcellations were compared with six public parcellations (Craddock et al., 2012; Shen et al., 2013; Joliot et al., 2015; Glasser et al., 2016; Gordon et al., 2016; Schaefer et al., 2018) using four evaluation metrics (Section 2.4). Five of these parcellations were generated with automatic algorithms from resting-fMRI data (Craddock et al., 2012; Shen et al., 2013; Joliot et al., 2015; Gordon et al., 2016; Schaefer et al., 2018). The remaining parcellation was generated from multimodal MRI data using a semi-automatic algorithm with inputs from an anatomist (Glasser et al., 2016).

Both the Glasser (Glasser et al., 2016) and AICHA (Joliot et al., 2015) parcellations were homotopic parcellations, while the remaining parcellations were non-homotopic. The Shen, Craddock and Schaefer parcellations are available in multiple resolutions. Given that there are estimated to be about 300 to 400 areas in the human cerebral cortex (Van Essen et al., 2012a), we chose the 400-region Craddock parcellation as a baseline. The most commonly used resolutions for the Shen and Schaefer parcellations were 268 and 400 parcels respectively, so they were chosen for comparison. Finally, the Gordon, Glasser and AICHA parcellations had 333, 360 and 384 parcels respectively.

For fair comparisons, there were several important issues to consider. First, the number of parcels had a strong influence on the evaluation metrics, so the hMRF parcellation procedure was run with different number of parcels to match the publicly available parcellations. Second, the Gordon parcellation included many unlabeled vertices near parcel boundaries, which artificially inflated the evaluation metrics. Therefore, when comparing with the Gordon parcellation, boundary vertices with the worst resting homogeneity in the GSP training set were removed from the hMRF parcellation to match the number of unlabeled vertices in the Gordon parcellation.

Another issue is that different parcellations were developed in different coordinate systems. The Shen, Craddock, and AICHA parcellations were in MNI152 volumetric space. The Glasser and Gordon parcellations were in fsLR surface space. The Schaefer and hMRF parcellations were in fsaverage6 surface space. On the other hand, the data used for computing the evaluation metrics were also in different coordinate systems. Data used for computing architectonic and visuotopy metrics were in fsLR space. The GSP resting-fMRI data, GUSTO resting-fMRI data, ABCD resting-fMRI data and ABCD task-fMRI data were in fsaverage6 space. The HCP resting-fMRI data and task-fMRI data were in fsLR space. HCP resting-fMRI data was also available in MNI152 space.

To compute the evaluation metrics, a parcellation was transformed into the coordinate system where the evaluation data resided. For example, to compute task activation inhomogeneity using the ABCD task-fMRI data, the Shen, Craddock, AICHA, Glasser, and Gordon parcellations were transformed into fsaverage6 surface space. On the other hand, to evaluate architectonic or visuotopic geodesic distances, the Shen, Craddock, AICHA, Schaefer and hMRF parcellations were projected to fsLR surface space. The non-linear transformations between fsLR, fsaverage and MNI spaces are detailed elsewhere (Van Essen et al., 2012a; https://wiki.humanconnectome.org/display/PublicData/HCP+Users+FAQ; Wu et al., 2018).

The two issues (number of parcels and coordinate systems) were not independent, since transforming a parcellation from one space to another might change the number of parcels. For example, after projecting the Craddock parcellation from MNI152 to fsaverage or fsLR space, the number of parcels was reduced from 400 to 355. Furthermore, the coverage of the parcellations varied across different coordinate systems, so an intersection procedure was applied to pairs of parcellations to ensure that each pair covered the same portion of the cerebral cortex. The intersection procedure can further reduce the number of parcels. For example, after the application of the intersection procedure, the resolution of the Craddock parcellation in MNI152 space dropped to 346 parcels. In this case, we ran the hMRF procedure twice, with 346 and 355 parcels, to match the Craddock parcellations in volumetric and surface spaces respectively. In the case of the Shen parcellation, there were 268 and 197 parcels in volumetric and surface spaces respectively. Therefore, the hMRF procedure was run twice, with 268 and 197 parcels, to match the Shen parcellations in volumetric and surface spaces respectively.

For the Shen, Craddock and Gordon parcellations, the number of parcels were different between the two hemispheres, which was problematic since hMRF parcellations have equal number of parcels on both hemispheres. In the case of the Shen parcellation in fsaverage6 space, there were 100 parcels in the left hemisphere and 97 parcels in the right hemisphere. Therefore, we ran the hMRF parcellation procedure with 200 parcels. The right hemisphere of the hMRF parcellation has more parcels than the right hemisphere of the Shen parcellation. Therefore, a merging procedure was applied to merge parcels in the right hemisphere of the hMRF parcellation (in a way that maximizes resting-homogeneity in the GSP training set), resulting in 97 parcels in the right hemisphere. The same procedure was repeated for the Craddock and Gordon parcellations.

Finally, when mapping a parcellation from MNI152 to fsaverage space, the parcellation boundary tended to become rough, leading to lower resting-homogeneity. To reduce such bias, we smoothed the boundaries of the Shen, Craddock and AICHA parcellations after projecting them from MNI152 to fsaverage space. The resulting parcellations were visually appealing, but it is of course not possible to fully eliminate biases that arose from the mismatch between a parcellation’s native coordinate system and the space in which the evaluation metrics was computed. For example, the AICHA parcellation (whose native space was MNI152) had inherent advantage over the hMRF parcellation (whose native space was fsaverage6) when computing resting-state homogeneity with the HCP resting-fMRI data in MNI152 space. As another example, the hMRF parcellation had inherent advantage over the AICHA parcellation when computing resting-state homogeneity with the ABCD resting-fMRI data in fsaverage6 space.

### 2.6 Cerebral cortical parcellations of 1479 participants and further characterization

The previous analyses served to compare the hMRF parcellations with other existing parcellations. To provide a final set of parcellations for the community, the hMRF parcellation procedure was finally applied to the full GSP dataset (N = 1479). Details regarding the hyperparameter settings can be found in Supplementary Methods S5. Since a single parcellation resolution was unlikely to be optimal across all applications, we generated parcellations (in the fsaverage6 surface space) from 100 to 1000 parcels in intervals of 100 parcels. The parcellations were also projected to fsLR and MNI152 spaces. To visualize the resulting parcellations, we assigned each parcel to one of 7 or 17 networks based on maximal spatial overlap with the Yeo networks (Yeo et al., 2011).

Since the Yeo networks were asymmetric, homotopic parcels in the hMRF parcellations could be assigned to different networks. We further characterized this phenomenon for the 400-region hMRF parcellation by computing RSFC between homotopic pairs of parcels. We hypothesized that homotopic parcels with different network assignments would exhibit weaker homotopic RSFC. Spatial overlap (Dice coefficient) for each pair of homotopic parcels was also computed based on our established cross-hemispheric vertex-level correspondence. More specifically, for each pair of homotopic parcels, we divided the total number of spatially homotopic vertices shared between both parcels by the total number of vertices enclosed in both parcels.

We also investigated the relationship between homotopic RSFC and task activation lateralization in the HCP dataset. More specifically, we considered the “story – math” contrast from the language, as well as task contrasts (tongue – average, left fingers – average, right fingers – average, right foot – average, left foot – average) from the motor task. For each activation contrast, we averaged the activation z values within each parcel of the 400- region hMRF parcellation. For each contrast and each homotopic parcel pair, an activation laterality index was defined as the absolute difference between left parcel activation and right parcel activation. Activation laterality was only computed for parcels strongly activated by the task contrast. We defined strongly activated parcels (of a particular task contrast) as those parcels whose average activation was at least 70% of the parcel with the largest average activation. In the case of the motor task, we averaged the activation laterality indices across the five contrasts, yielding a single activation laterality map for the motor task.

Finally, we further characterized the 400-region hMRF parcellation by overlaying the boundaries of histological areas and visuotopical cortical areas. Visual inspection was then performed to explore the agreement between the hMRF parcellation and cortical areal boundaries.

### 2.7 Data and code availability

Code for the hMRF model can be found here (GITHUB_LINK). Co-authors (RK, AX and LA) reviewed the code before merging into the GitHub repository to reduce the chance of coding errors. Multi-resolution hMRF parcellations generated from 1489 participants are publicly available (GITHUB_LINK).

The HCP preprocessing code can be found here (https://github.com/Washington-University/HCPpipelines). The remaining datasets utilized the CBIG preprocessing pipeline (https://github.com/ThomasYeoLab/CBIG/tree/master/stable_projects/preprocessing/CBIG_f <u>MRI_Preproc2016</u>). The datasets were processed by different researchers, so the pipeline was run with slightly different parameters (as reported in Section 2.2).

The GSP (http://neuroinformatics.harvard.edu/gsp/), HCP (https://www.humanconnectome.org/) and ABCD (https://nda.nih.gov/abcd/) datasets are publicly available. The list of ABCD participants used in this study have been uploaded to the NIMH Data Archive (NDA). Researchers with access to the ABCD data will be able to access the participant list: https://nda.nih.gov/study.html?id=XXX. The GUSTO dataset can be obtained via a data transfer agreement (https://www.gusto.sg/).

## 3 Results

### 3.1 Homotopic differences between hMRF and gwMRF parcellations within area 17

The left primary visual area V1 receives stimuli from the right visual hemifield, while the right primary visual area V1 receives stimuli from the left visual hemifield (Wandell et al., 2007). Given that the cross-hemisphere mapping is symmetric, we do not expect an asymmetric parcellation of V1 at the resolution of our parcellations. Given strong correspondence between histologically-defined architectonic area 17 and V1, here we use architectonic area 17 as a proxy of V1.

Figure 1 overlays the boundaries of area 17 (Amunts et al., 1999) on the 400-region parcellations generated by gwMRF and hMRF. Both parcellations agree well with area 17 boundary. However, the gwMRF parcellation subdivided left area 17 into three parcels and right area 17 into two parcels (Figure 1B). On the other hand, the hMRF parcellation subdivided left and right areas 17 into equal number of parcels with similar spatial topography across the two hemispheres, consistent with the fact that area 17 is not known to be asymmetric.

**Figure 1.**
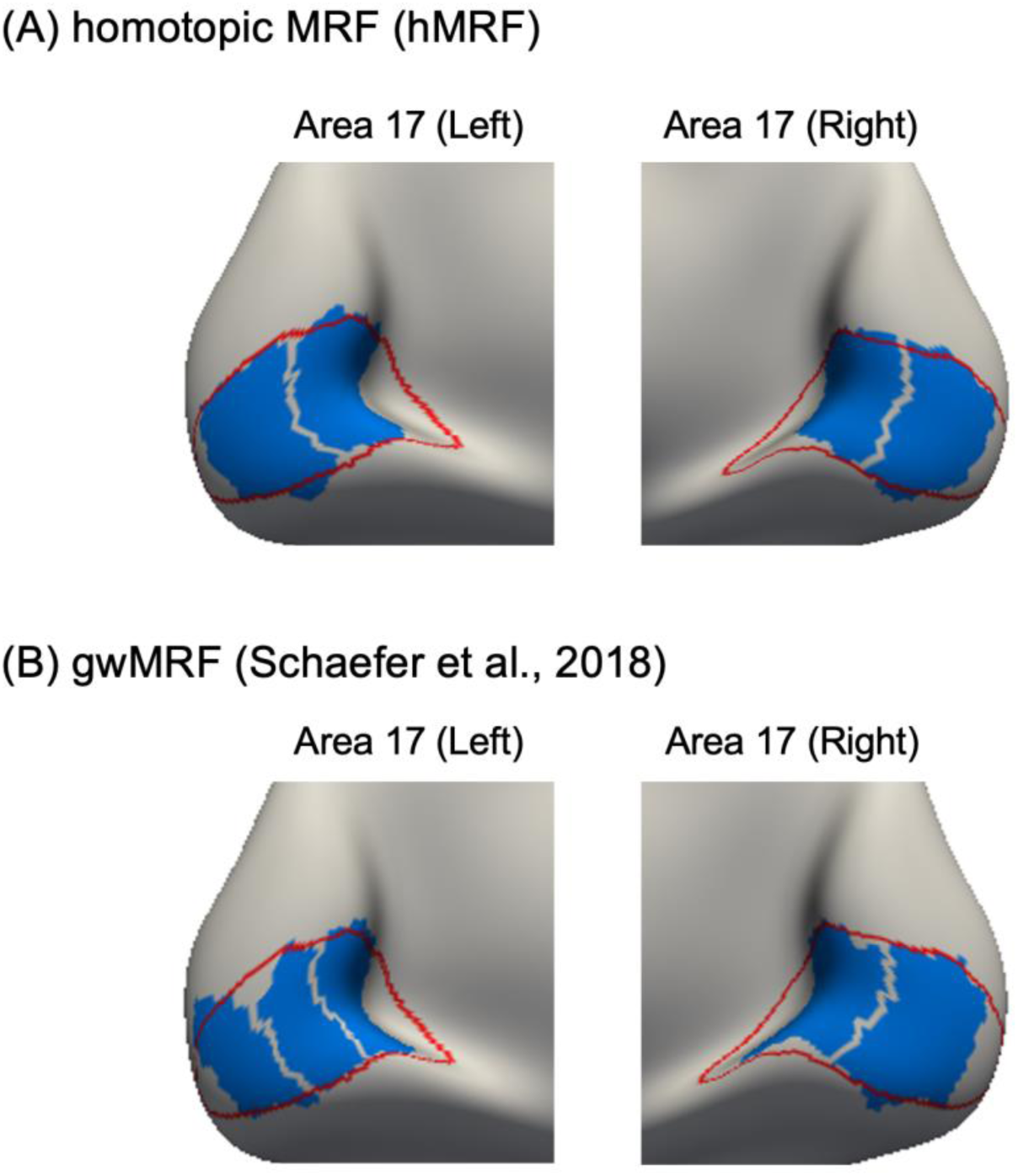
Difference in homotopic correspondence between hMRF and gwMRF parcellations within histologically-defined area 17. (A) Parcels (blue) of the 400-region hMRF parcellation within histological (red) boundaries of area 17. The hMRF parcellation subdivided left and right areas 17 into equal number of parcels with similar spatial topography across the two hemispheres. (B) Parcels (blue) of the 400-region gwMRF parcellation within histological (red) boundaries of area 17. The gwMRF parcellation subdivided left area 17 into three parcels and right area 17 into two parcels.

Comparison of hMRF parcellations with other parcellations are shown in Figure S1. The homotopic parcellations (AICHA, Glasser and hMRF) and the Gordon parcellation yielded homotopic parcels that were well-aligned with area 17. The non-homotopic Craddock parcellation subdivided left area 17 into two parcels and right area 17 into three parcels. The non-homotopic Shen parcellation yielded roughly homotopic parcels that overlapped with area 17, but the parcel boundaries did not align with the boundary of area 17.

### 3.2 Architectonic alignment

Figure S2 shows the geodesic distances between the boundaries of 10 histologically-defined architectonic areas and six publicly available parcellations. Geodesic distances between the architectonic areas and hMRF parcellations are also shown. Number of parcels was matched between the publicly available parcellations and corresponding hMRF parcellations. Lower distance indicates better alignment.

Across the 10 architectonic areas, the hMRF approach generated parcellations with similar architectonic distance as Shen (p = 0.441, average 3.8%), better distance than Gordon (p ≈ 0, average 24.7%), similar distance to AICHA (p = 0.210, average 8.2%), better distance than Craddock (p = 0.031, average 11.2%), worse distance than Glasser (p = 0.032, average 17.9%), and similar distance to Schaefer (p = 0.835, average 1.0%).

After correcting for multiple comparisons with a false discovery rate (FDR) of q < 0.05, only the comparison with the Gordon parcellation remained significant. Overall, the hMRF parcellations exhibited architectonic alignment comparable with (or better than) other parcellations.

### 3.3 Visuotopic alignment

Figure S3 shows the geodesic distances between the boundaries of 18 visuotopic areas (Abdollahi et al., 2014) and six publicly available parcellations. Geodesic distances between the visuotopic areas and hMRF parcellations are also shown. Number of parcels was matched between the publicly available parcellations and corresponding hMRF parcellations. Lower distance indicates better alignment.

Across the 18 visuotopic areas, the hMRF parcellations achieved similar visuotopic distance to Shen (p = 0.279, average 10.5%), better distance to Gordon (p ≈ 0, average 39.7%), similar distance to ACIHA (p = 0.731, average 3.3%), similar distance to Craddock (p = 0.989, average 0.1%), worse distance to Glasser (p ≈ 0, average 46.3%) and similar distance to Schaefer (p = 0.083, average 16.8%).

After correcting for multiple comparisons with an FDR of q < 0.05, the comparisons with the Gordon and Glasser parcellations remained significant. We note that the Glasser parcellation was derived with a semi-automated algorithm that required an anatomist to manually select multi-modal information to match prior knowledge of areal boundaries. Overall, the hMRF parcellations achieved visuotopic alignment comparable with (or better than) other fully automatic approaches.

### 3.4 Task-fMRI activation inhomogeneity

Figure 2 compares the task inhomogeneity of the hMRF parcellations with six publicly available parcellations using the HCP task-fMRI data. A lower task inhomogeneity indicated higher parcellation quality. Task inhomogeneity was not comparable between parcellations of different resolutions, so the number of parcels was matched between the publicly available parcellations and corresponding hMRF parcellations.

**Figure 2.**
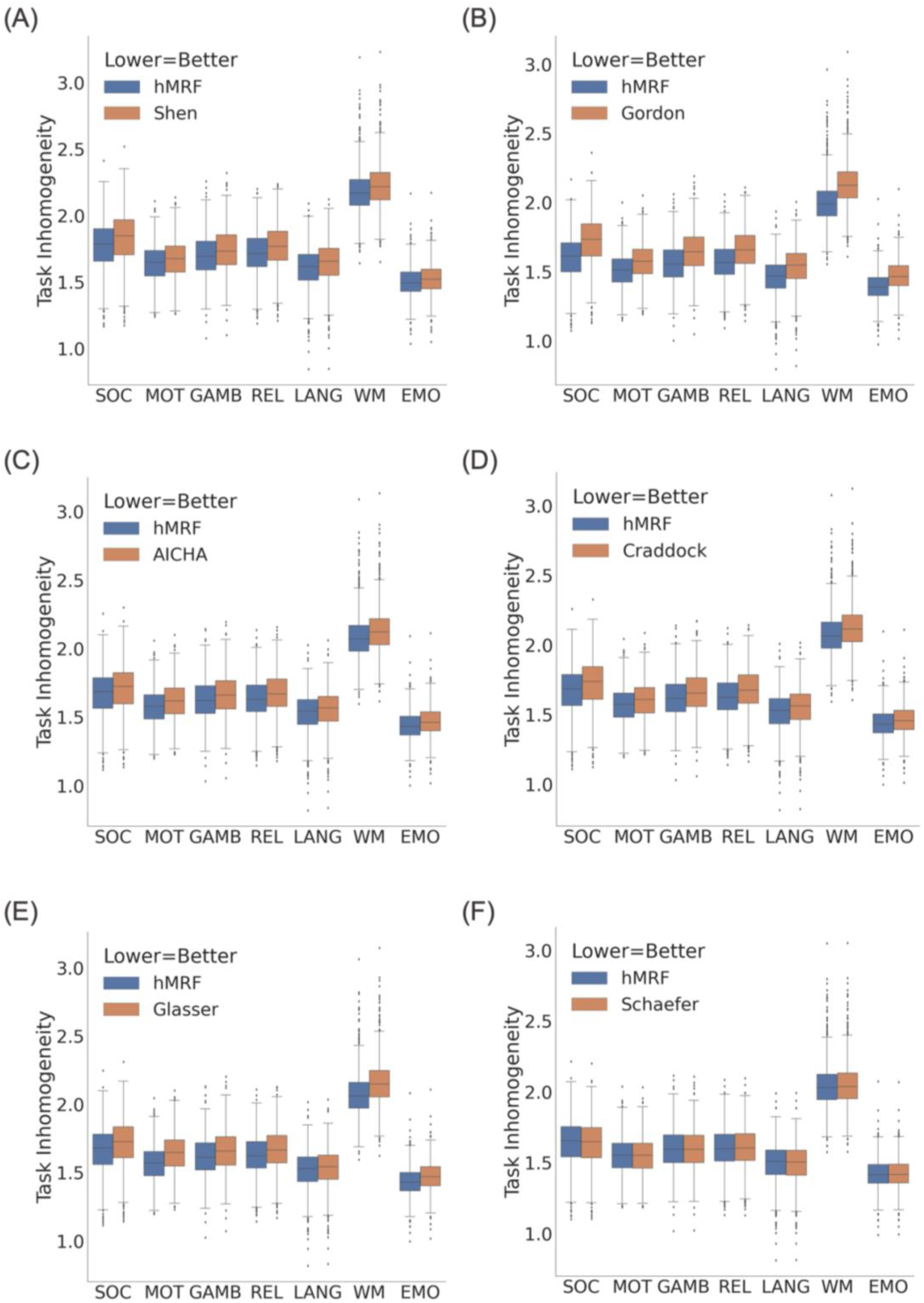
Task inhomogeneity computed from task-fMRI data in the HCP dataset (N = 1030). Lower task inhomogeneity indicates better parcellation quality. We note that task inhomogeneity cannot be compared across the panels because of the different number of parcels across panels. The hMRF parcellations exhibited comparable task inhomogeneity with the Schaefer parcellation and better task inhomogeneity than all other parcellations.

In the HCP dataset (Figure 2), the hMRF parcellations exhibited lower (better) task inhomogeneity than the Shen (average 2.4%; p ≈ 0), Gordon (average 5.6%; p ≈ 0), AICHA (average 2.1%; p ≈ 0), Craddock (average 2.4%; p ≈ 0), Glasser (average 2.9%; p ≈ 0) and Schaefer (average 0.06%; p = 3.0e-17) parcellations. It is worth mentioning that the Glasser parcellation was partially derived using task-fMRI data from the HCP dataset, so should have an inherent advantage in this metric.

Figure 3 compares the task inhomogeneity of the hMRF parcellations with six publicly available parcellations using the ABCD task-fMRI data. The hMRF parcellations exhibited lower (better) task inhomogeneity than the Shen (average 1.5%, p ≈ 0), Gordon (average 4.9%; p ≈ 0), AICHA (average 2.0%; p ≈ 0), Craddock (average 1.8%; p ≈ 0) and Glasser (average 4.0%; p ≈ 0) parcellations. On the other hand, the hMRF parcellations exhibited higher (worse) task inhomogeneity than the Schaefer parcellation (average 0.02%; p = 4.9e-08).

**Figure 3.**
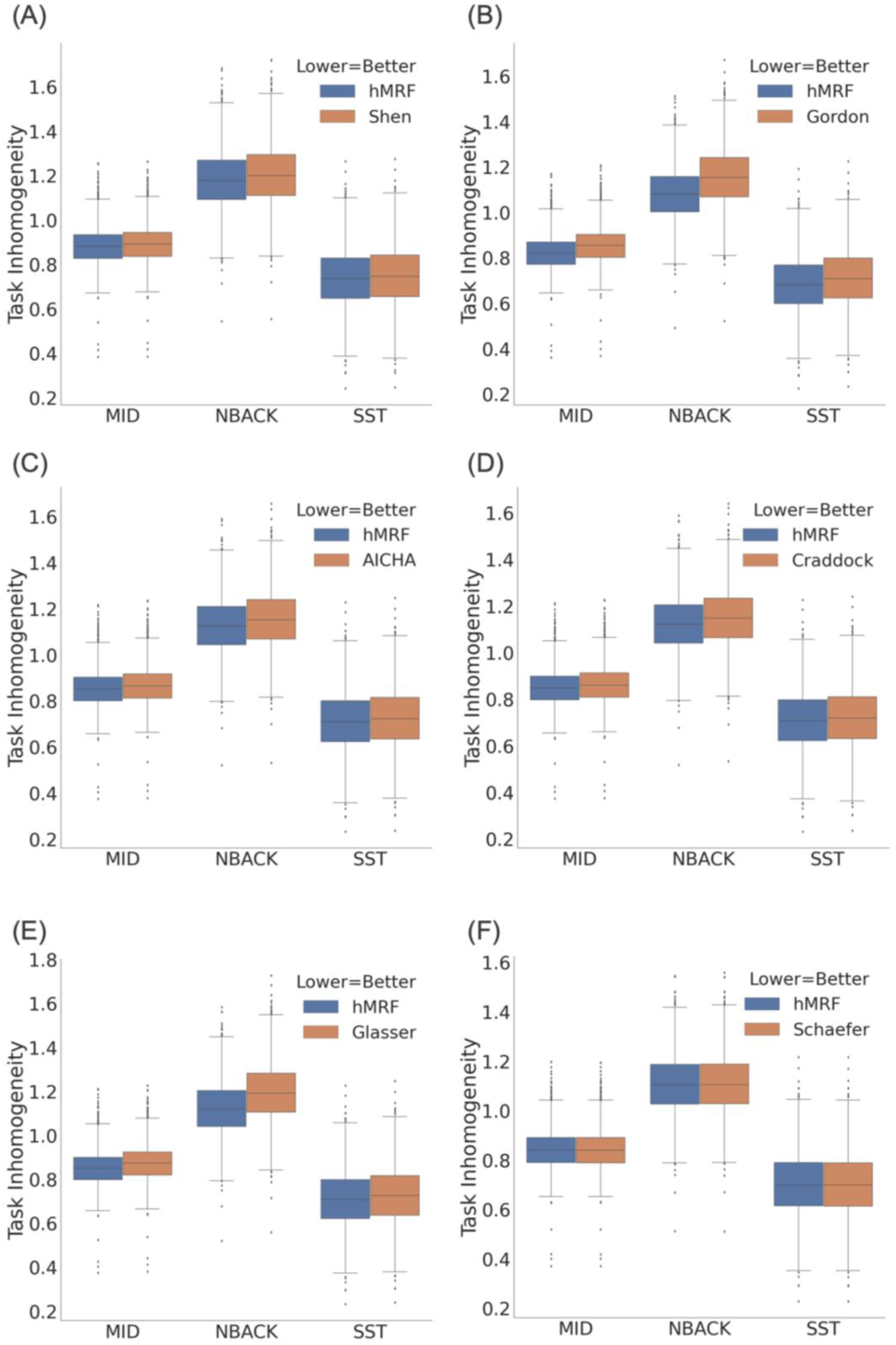
Task inhomogeneity computed from task-fMRI data in the ABCD dataset (N = 2262). Lower task inhomogeneity indicates better parcellation quality. We note that task inhomogeneity cannot be compared across the panels because of the different number of parcels across panels. The hMRF parcellations exhibited comparable task inhomogeneity with the Schaefer parcellation and better task inhomogeneity than all other parcellations.

Overall, the hMRF parcellations exhibited comparable task inhomogeneity with the Schaefer parcellation and better task inhomogeneity than the other five parcellations. In particular, differences between hMRF and Schaefer parcellations were less than 0.1%, which was an order of magnitude smaller than the differences between hMRF and non-Schaefer parcellations. Differences between hMRF and non-Schaefer parcellations were larger, but still modest with differences in the order of 1% to 5% on average.

### 3.5 Resting-fMRI homogeneity

Figures 4 and S3 compare the hMRF parcellations with 6 publicly available parcellations in terms of resting-fMRI homogeneity across a variety of datasets. Higher resting-fMRI homogeneity indicated higher parcellation quality. Resting-fMRI homogeneity was not comparable between parcellations of different resolutions, so the number of parcels was matched between the publicly available parcellations and corresponding hMRF parcellations.

**Figure 4.**
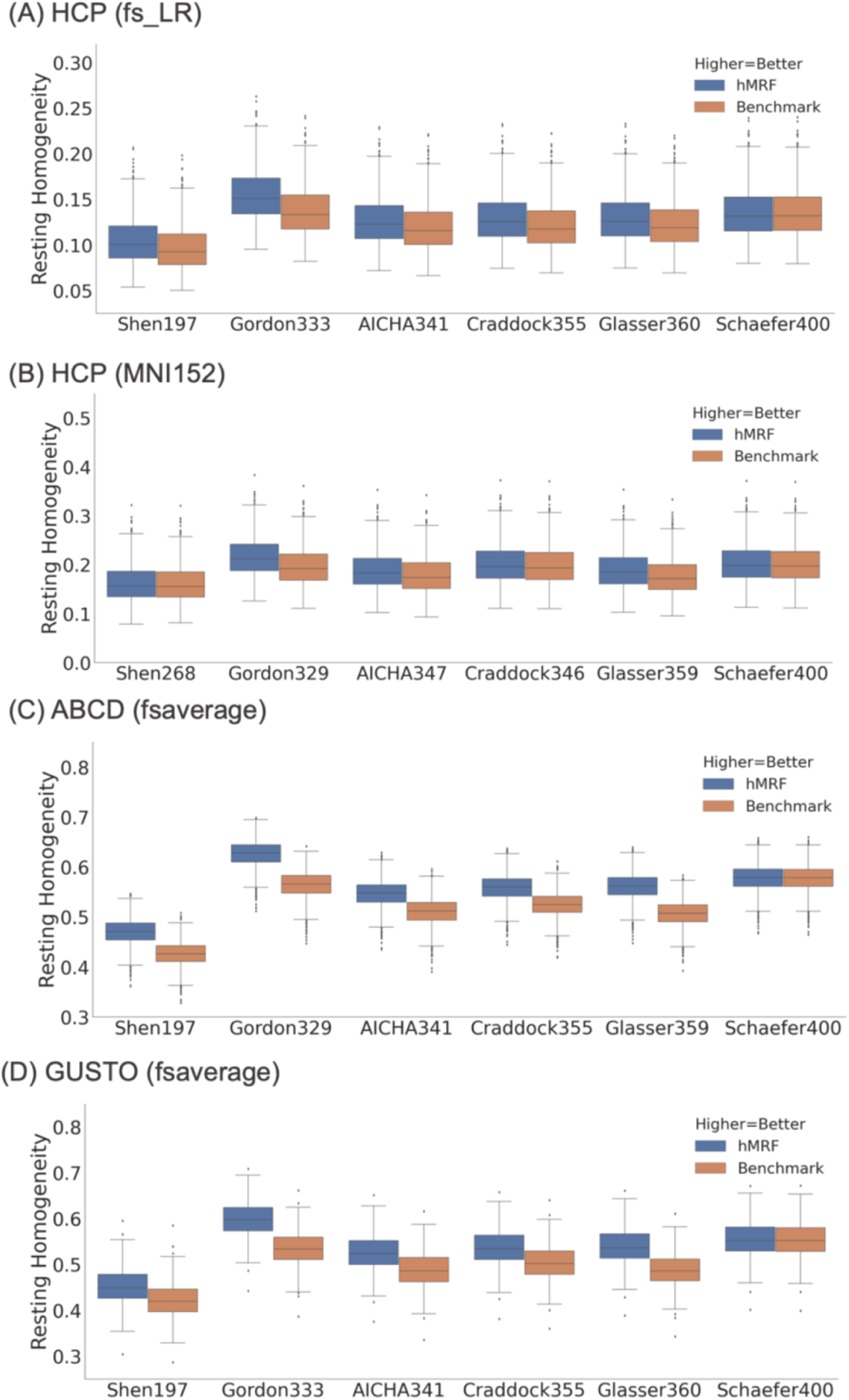
Resting-fMRI homogeneity computed with the (A) HCP dataset (N = 1030) in fsLR space, (B) HCP dataset (N = 1030) in MNI152 space, (C) ABCD dataset (N = 2262) in fsaverage space, and (D) GUSTO dataset (N = 393) in fsaverage space. We note that resting- fMRI homogeneity was not comparable across different publicly available parcellations because of differences in the number of parcels. However, the number of parcels was matched between the publicly available parcellations and corresponding hMRF parcellations. The hMRF parcellations achieved comparable resting-fMRI homogeneity with the Schaefer parcellation. On the other hand, the hMRF parcellations were more homogeneous than the 5 non-Schaefer parcellations across all data sets. Resting-fMRI homogeneity in the GSP test set is shown in Figure S4.

The hMRF parcellations were significantly more homogeneous across all four datasets than the Shen (average 6.6%, p < 7.7e-37), Gordon (average 11.5%, p ≈ 0), AICHA (average 6.6%, p ≈ 0), Craddock (average 5.5%, p < 9.8e-324), and Glasser (average 9.0%, p < 9.5e-284) parcellations. As noted above, the Glasser parcellation was partially derived from the HCP resting-fMRI data, so should have an inherent advantage in the HCP dataset. On the other hand, the hMRF and Schaefer parcellations were comparable with mean difference of 0.11% across all datasets.

A parcellation with higher number of parcels should result in higher resting-fMRI homogeneity. However, we note that the 333-region Gordon parcellation exhibited higher resting-fMRI homogeneity than other parcellations of higher resolutions (e.g., Glasser parcellation). The reason is that the Gordon parcellation unlabeled vertices at the boundaries between parcels, which artificially boosted its homogeneity. Similarly, when generating hMRF parcellation for comparison with the Gordon parcellation, we unlabeled the same number of vertices as the Gordon parcellation, which boosted our resting-fMRI homogeneity.

We also note that the resting-fMRI homogeneity was much lower in the HCP dataset (in fsLR space) compared with the GSP and ABCD datasets. A major reason is that the HCP dataset was smoothed with a 2mm FWHM (in fsLR space), while the GSP and ABCD datasets were smoothed with a 6mm FWHM. In general, greater spatial smoothing should yield higher resting-fMRI homogeneity (on average).

In the current results (Figure 4), the HCP data in MNI152 space was smoothed by 6 mm. As a control analysis, we repeated the analysis with no smoothing. Unsurprisingly, compared with 6mm smoothing (Table S1B), resting-fMRI homogeneity of all parcellations were significantly lower when there was no spatial smoothing (Table S1A). Nevertheless, the hMRF parcellations were significantly more homogeneous than the non-Schaefer parcellations with improvements ranging from 3% to 12%. Interestingly, the percentage improvements over non-Schaefer parcellations were actually higher when there was no spatial smoothing. Similar to the previous results, the hMRF and Schaefer parcellations were comparable with mean difference of only 0.16%.

Finally, we also compared the hMRF parcellation with the popular Automated Anatomical Labelling (AAL) atlas (Rolls et al., 2020). Given that the hMRF parcellation was derived from resting-state fMRI, we expected the hMRF parcellation to have an inherent advantage over the AAL atlas. Indeed, the hMRF parcellation exhibited significantly better resting-state homogeneity than the AAL atlas across all datasets (Table S2).

### 3.6 Homotopic resting-state functional connectivity

Figures 5 and S5 compare the hMRF parcellations with two homotopic parcellations in terms of homotopic resting-state functional connectivity. The hMRF parcellations exhibited stronger homotopic resting-state functional connectivity than the two homotopic parcellations: Glasser (mean 7.8%; p ≈ 0) and AICHA (mean 4.9%; p ≈ 0). Similar conclusions were obtained using HCP data in MNI52 space with no smoothing (Table S3).

**Figure 5.**
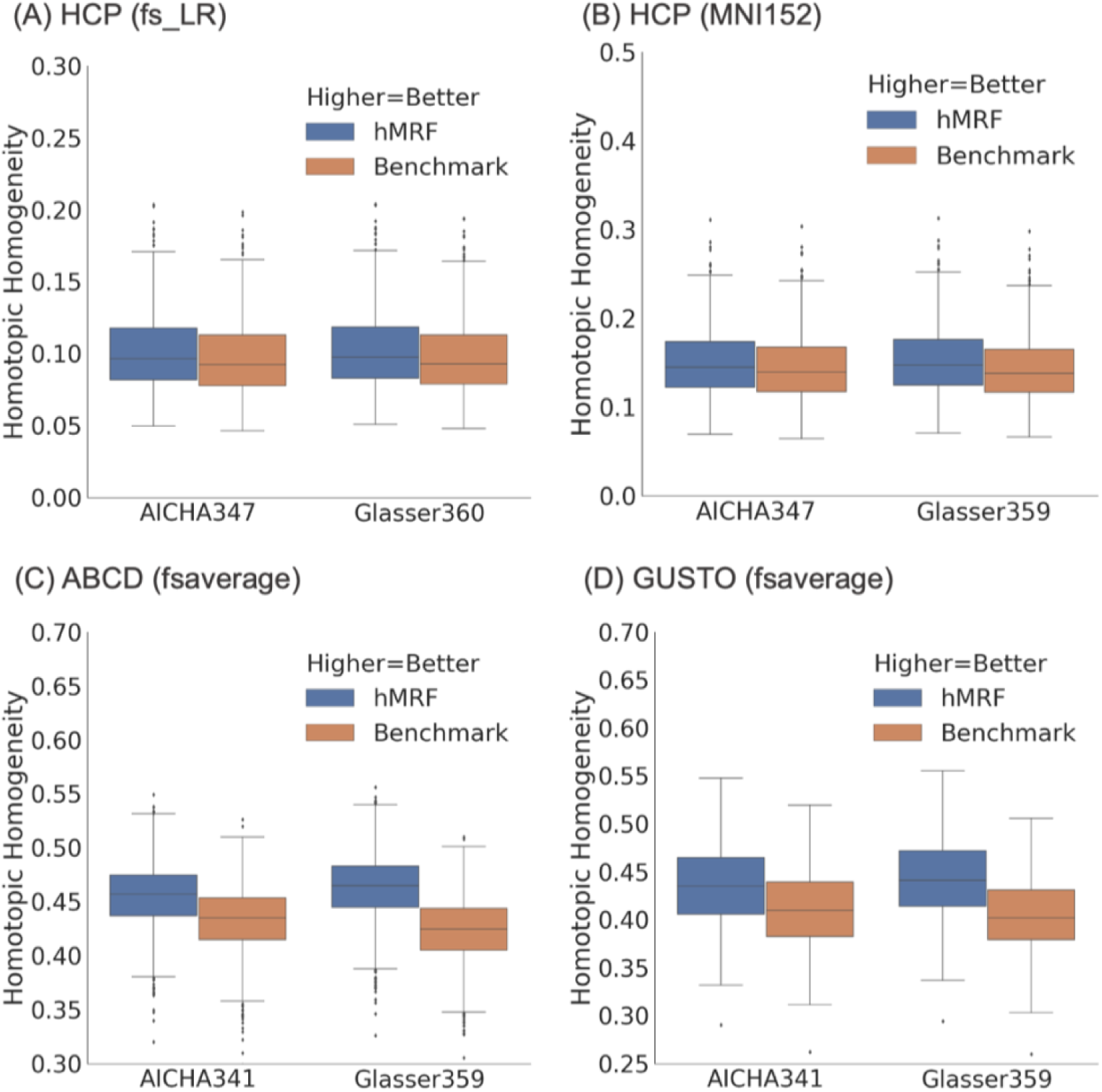
Homotopic resting-state functional connectivity in the (A) HCP dataset (N = 1030) in fsLR space, (B) HCP dataset (N = 1030) in MNI152 space, (C) ABCD dataset (N = 2262) in fsaverage space, and (D) GUSTO dataset (N = 393) in fsaverage space. We note that homotopic functional connectivity was not comparable between the AICHA and Glasser parcellations because of differences in the number of parcels. The hMRF parcellations achieved higher (better) homotopic functional connectivity than the Glasser and AICHA homotopic parcellations. Results in the GSP test set is shown in Figure S5.

### 3.7 RSFC lateralization of hMRF parcellations estimated from 1479 Participants

Cerebral parcellations at multiple resolutions, from 100 to 1000 parcel at every 100- parcel interval, were generated from the full GSP dataset (N = 1479) and visualized in Figure S6. The first row of Figure 6 shows the Yeo 7-network and 17-network parcellations (Yeo et al., 2011). The second row of Figure 6 shows the 400-region hMRF parcellation, where the color of each parcel was assigned based on maximal spatial overlap with the Yeo networks. Network assignment for parcellations of other resolutions are shown in Figure S7.

**Figure 6.**
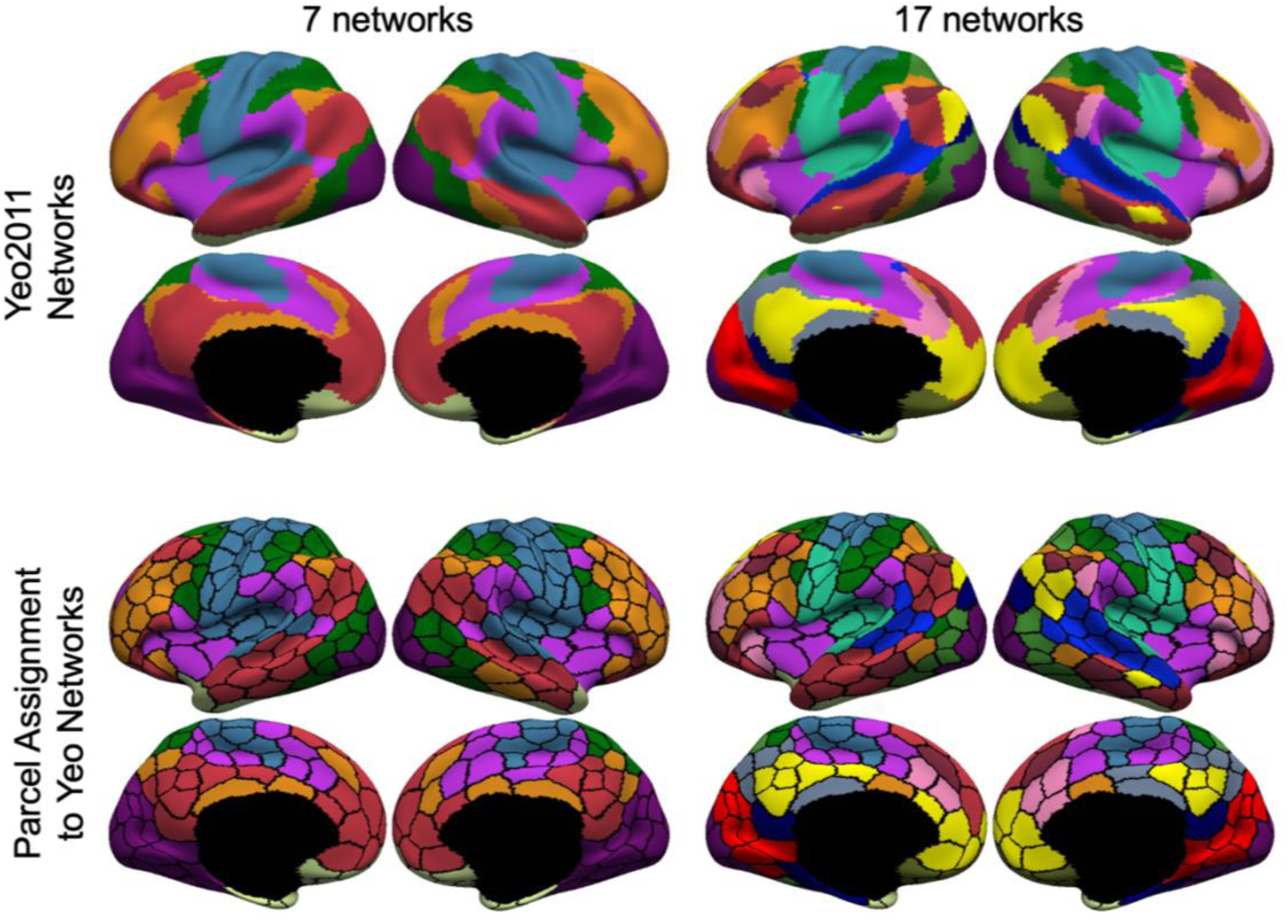
Assignment of parcels to 7 or 17 networks for hMRF parcellations with 400 regions. First row shows the Yeo 7 and 17 networks (Yeo et al., 2011). Second row shows the 400-region hMRF parcellation with each parcel assigned a network color based on its spatial overlap with the 7-network or 17-network parcellation.

Some of the Yeo networks were clearly asymmetric across the hemispheres. For example, the frontoparietal control network (orange network in the first column of Figure 6) was larger in the right lateral prefrontal cortex than in the left lateral prefrontal cortex. This lateralization difference translated to the network assignment of the hMRF parcellations.

Figure 7A shows the 7-network assignment of the 400-region hMRF parcellation with black arrows indicating a pair of homotopic parcels assigned to different networks. The left parcel was assigned to the default network, while the right parcel was assigned to the frontoparietal control network.

**Figure 7.**
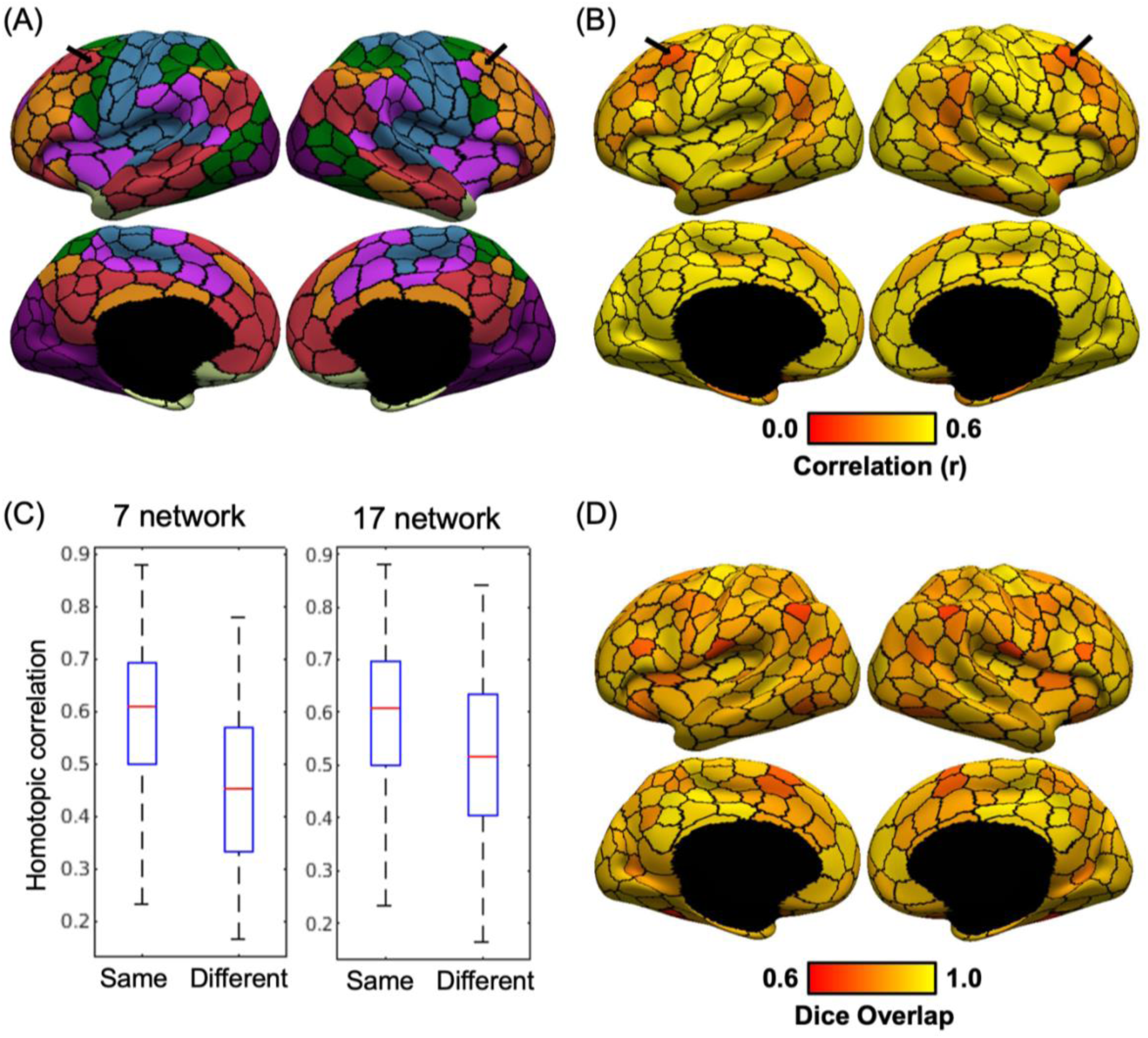
RSFC lateralization of the 400-region hMRF parcellation. (A) Lateralization in network assignment of the 400-region hMRF parcellation. Black arrows indicate a homotopic pair of regions assigned to two different networks from the Yeo 7-network parcellation. (B) RSFC between homotopic pairs of parcels in the GSP dataset. Black arrows indicate that the homotopic pair of parcels highlighted in panel A exhibited weaker RSFC with each other (i.e., homotopic correlation) compared with other homotopic parcel pairs. (C) Boxplot of RSFC between homotopic parcel pairs assigned to the same or different networks for both 7- network and 17-network assignments. Homotopic parcel pairs assigned to the same network exhibited stronger RSFC than homotopic parcel pairs assigned to different networks. (D) Spatial overlap (as measured by Dice coefficient) between homotopic pairs of parcels.

Figure 7B shows the RSFC between homotopic pairs of parcels in the 400-region hMRF parcellation in the GSP dataset. As indicated by the black arrows, the homotopic parcels with different network assignment shown in Figure 7A, also exhibited weaker RSFC compared to the rest of the cerebral cortex. This phenomenon is further quantified in Figure 7C, which shows that homotopic parcel pairs assigned to the same networks exhibited stronger RSFC than homotopic parcel pairs assigned to different networks (p = 3.72e-4 for 7-network assignment and p = 0.0028 for 17-network assignment). Therefore, lateralization differences in the spatial topography of the Yeo networks were reflected in the strength of RSFC between homotopic parcels.

Another clear observation was that sensory-motor parcels showed stronger homotopic RSFC (Figure 7B), while association parcels showed weaker RSFC, consistent with previous studies (Stark et al., 2008; Zuo et al., 2010). Finally, Figure 7D shows the degree of spatial overlap (as measured by the Dice coefficient on the fsaverage6 surface) between homotopic pair of parcels. The degree of spatial overlap varied across the cortex with no clear pattern.

There was almost no correlation between the spatial overlap (Figure 7D) and RSFC (Figure 7B) of homotopic parcel pairs (r = 0.20). In other words, homotopic parcel pairs with greater spatial overlap did not necessarily have stronger or weaker homotopic RSFC.

### 3.8 Task activation lateralization of the 400-region hMRF parcellation

Figures S8 to S12 show the HCP task contrasts overlaid on the 400-region hMRF parcellation boundaries. Consistent with previous parcellations (e.g., Schaefer et al., 2018), task activations were highly distributed across the cortex, cutting across multiple parcellation boundaries. This is expected given that a single task contrast involves multiple cognitive processes supported by multiple cortical areas (Poldrack 2006; Barrett and Satpute 2013; Yeo et al., 2015).

Figure 8A shows the unthresholded “story – math” language task contrast, in which activation z values were averaged within each of the 400-region hMRF parcels. Figure 8B shows the task activation laterality index defined as the absolute difference between right hemisphere and left hemisphere activation values. This laterality index was only computed for parcels whose average activation-z values were at least 70% of the most activated parcel. Figure 8C shows the RSFC between homotopic pairs of parcels in the HCP dataset. Because the HCP dataset exhibited a strong posterior-to-anterior SNR gradient (Figure S13), the SNR map was regressed from the raw homotopic correlations (Figure 8D). We note that this SNR gradient was absent in the GSP dataset (see Figure 3 of Yeo et al., 2011), so a similar regression was not performed in the GSP dataset (Figure 7B).

**Figure 8.**
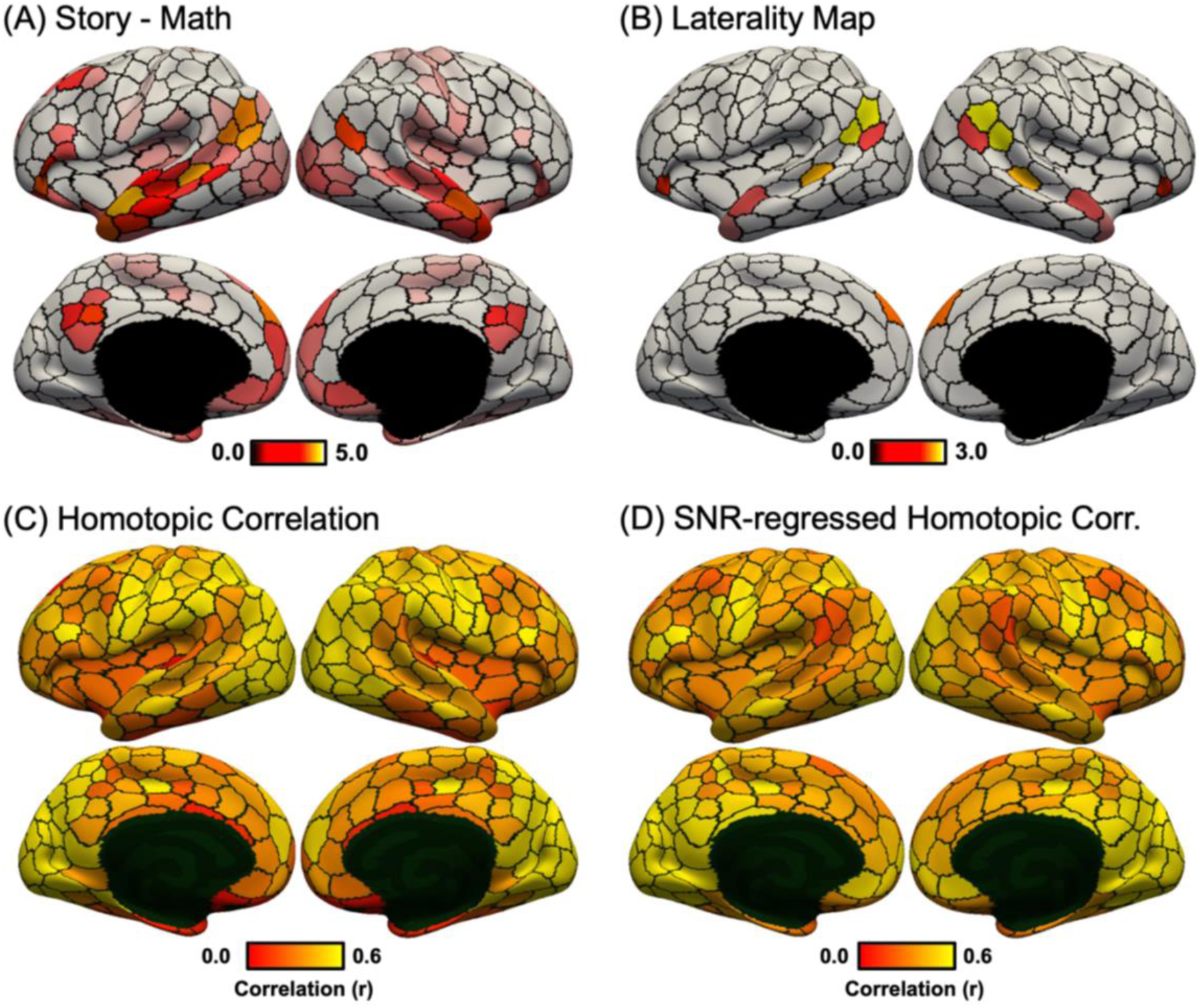
Language task activation lateralization in the HCP dataset. (A) Group-average “story – math” language task contrast in the HCP dataset. Activation z values were averaged within each of the 400-region hMRF cortical parcellation (shown as black boundaries). (B) Group-average task laterality map defined as the absolute difference between left and right hemisphere activation values. Laterality was only computed for parcels whose average activations were at least 70% of the most activated parcel (number of suprathreshold parcels = 16). Laterality maps for alternative thresholds can be found in Figure S14. (C) RSFC between homotopic pairs of parcels in the HCP dataset. (D) RSFC between homotopic pairs of parcels in the HCP dataset after regressing HCP SNR map (Figure S13). There was a negative correlation between task activation laterality (Figure 8B) and SNR-regressed homotopic correlations (Figure 8D).

We hypothesized that homotopic parcels with greater task activation laterality also exhibited weaker homotopic correlations. This hypothesis was confirmed by the data: correlation between language task activation laterality (Figure 8B) and SNR-regressed homotopic correlation (Figure 8D) was negative (r = -0.76; spin test p = 0.012). Similar results were obtained with the motor task (Figure 9). As expected, finger and foot activations were lateralized while the tongue activation was bilateral (Figure 9F). Correlation between motor task activation laterality (Figure 9F) and SNR-regressed homotopic correlation (Figure 8D) was negative (r = -0.83; spin test p = 0.028). Similar results were also obtained with different activation thresholds (Figure S14).

**Figure 9.**
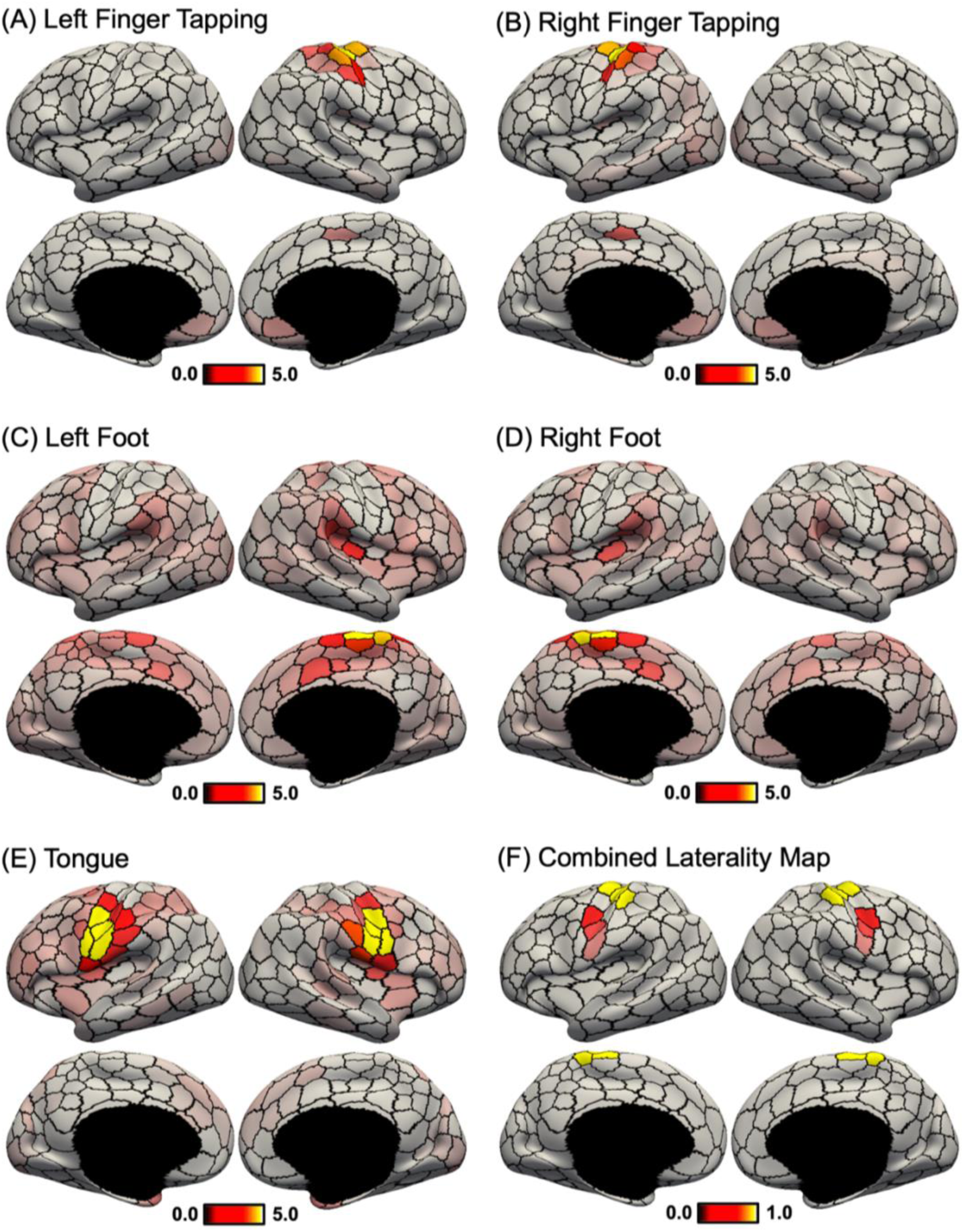
Motor task activation lateralization in the HCP dataset. (A) Group-average “left finger – average” motor task contrast. Activation z values were averaged within each of the 400-region hMRF cortical parcellation (shown as black boundaries). (B) Group-average “right finger – average” motor task contrast. (C) Group-average “left foot – average” motor task contrast. (D) Group-average “right foot – average” motor task contrast. (E) Group-average “tongue – average” motor task contrast. (F) Task laterality map averaged across the five contrasts. Similar to Figure 8, task laterality was defined as the absolute difference between left and right hemisphere activation values. Laterality was only computed for parcels whose average activations were at least 70% of the most activated parcel (number of suprathreshold parcels = 16). Laterality maps for alternative thresholds can be found in Figure S14. There was a negative correlation between task activation laterality (Figure 9F) and SNR-regressed homotopic correlations (Figure 8D).

### 3.9 Comparison of 400-region hMRF parcellation with architectonic and visuotopic areas

Figure 10 shows the 400-region hMRF parcellation (estimated from the full GSP dataset) overlaid with boundaries of architectonic areas 3, 4, 2, hOc5, and 17 on the right cortical hemisphere (Fischl et al., 2008; Van Essen et al., 2012a). Figures for other architectonic areas as well as those on the left hemisphere are shown in Figure S15.

**Figure 10.**
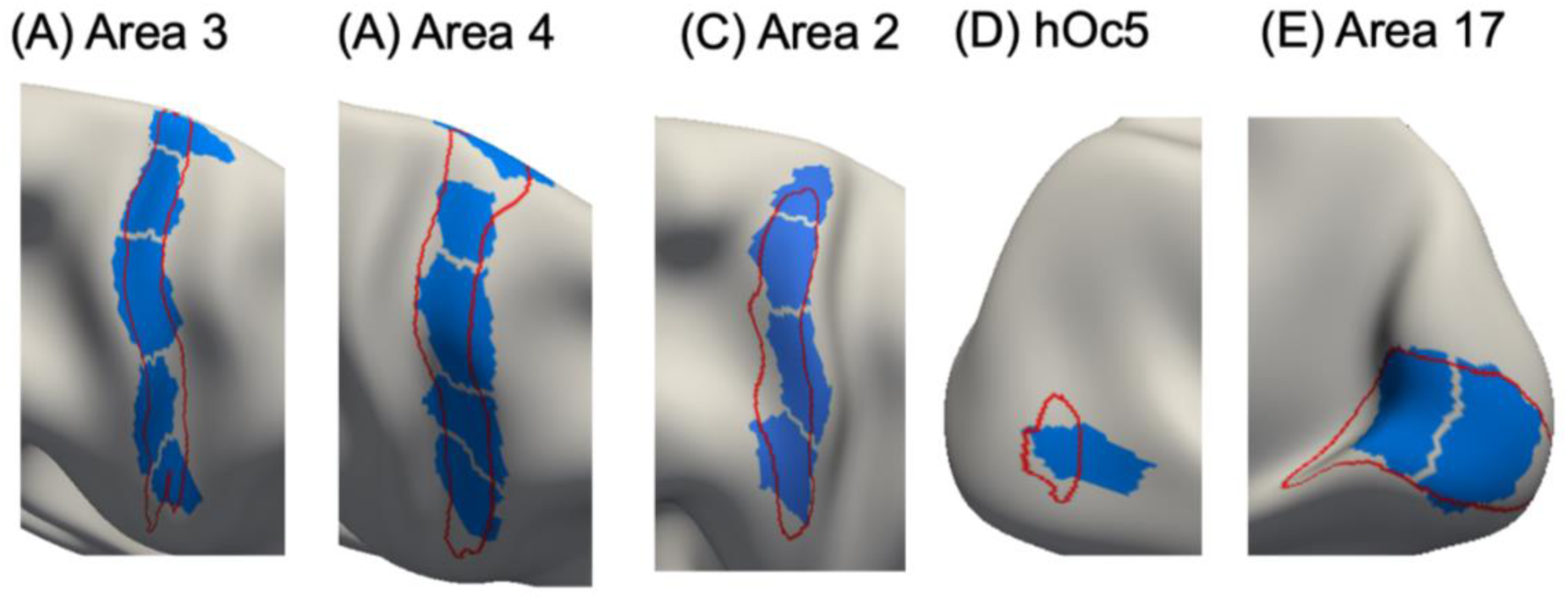
Parcels (blue) of the 400-region hMRF parcellation overlaid on histological (red) boundaries of right hemisphere (A) area 3, (B) area 4, (C) area 2, (D) hOc5, and (E) area 17. Other histological areas are shown in Figure S15.

The parcels corresponded well with the histological boundaries (Figure 10). However, consistent with previous studies, the parcellation also seemed to fractionate each area into multiple subunits. Within areas 3, 4, and 2, the fractionated parcels might correspond to somatotopic representations of different body parts. Parcels within primary area 17 appeared to fractionate the eccentricity axis. Area 44 was captured well by the parcellation on both hemispheres (Figure S15).

For other areas, the alignment between the parcels and areal boundaries was less precise. The parcels overlapping with area 18 aligned well with the boundary between areas 18 and 17, but tended to extend beyond the boundary between areas 18 and 19. The parcellation was not able to capture area 1 boundary, which was long and thin. Area 45 maximally overlapped with two hMRF parcels on each hemisphere. For area 6, the alignment between the parcels and areal boundary on the left hemisphere was more precise than on the right hemisphere.

Figure S16 overlays the 400-region parcellation boundary on 18 visuotopically mapped areas (Adollahi et al., 2014). The parcels overlapping with visuotopically-defined V1 (which is known to correspond to architectonic area 17) were in good agreement with V1 boundaries. For other visuotopic areas, correspondence was more muted. The parcels might fractionate higher order visuotopic areas based on eccentricity, similar to the fractionation within V1.

## 4 Discussion

In this study, we extended the gwMRF model to derive a homotopic parcellation of the cerebral cortex. Compared with the original gwMRF parcellation (Schaefer et al., 2018), the resulting hMRF parcellation exhibited similar alignment to architectonic and visuotopic boundaries, as well as similar task-state inhomogeneity and resting-state homogeneity.

Furthermore, the hMRF parcellations exhibited better task inhomogeneity and resting homogeneity than five publicly available parcellations (Craddock et al., 2012; Shen et al., 2013; Joliot et al., 2015; Glasser et al., 2016; Gordon et al., 2016). Consistent with previous studies, we found that weaker correlations between homotopic parcels were associated with greater lateralization in resting network organization, as well as greater lateralization in language and motor task activation. Similar to the Schaefer parcellation, the 400-region homotopic parcellation agreed with the boundaries of a number of cortical areas estimated from histology and visuotopic fMRI, while capturing sub-areal (e.g., somatotopic and visuotopic) features. Therefore, while the parcels do not correspond to cortical areas, we believe the hMRF parcellations represent meaningful functional units of the human cerebral cortex, and the multi-resolution parcellations will be a useful resource for future studies.

### 4.1 Interpretation and potential applications of the hMRF parcellations

Histological studies have generally found pairs of cortical areas in roughly spatially homotopic locations across the left and right hemispheres (Amunts et al., 2020). In fact, many histological studies simply report cortical areas on one hemisphere with the strong implicit assumption that homotopic cortical areas exist on the other hemisphere (e.g., Petrides and Pandya, 1999). However, it is important to note that homotopic areas do not necessarily have the same function as demonstrated by well-known lateralization of brain activation during language tasks (Malik-Moraleda et al., 2022)

Similar functional lateralization was also observed for the 400-region hMRF parcellation during both resting and task states. For example, a number of homotopic parcels overlapped with different large-scale resting-state networks (Yeo et al., 2011). Homotopic parcel pairs assigned to the same network exhibited stronger RSFC than homotopic parcel pairs assigned to different networks (Figure 7C). Sensory-motor parcels also exhibited stronger homotopic correlations than association parcels in both GSP (Figure 7B) and HCP (Figures 7D) datasets, consistent with previous studies of homotopic correlations (Stark et al., 2008; Zuo et al., 2010).

A potential mechanism for the observed sensory-association gradient in homotopic correlations is that sensory-motor regions are connected by thickly myelinated fast conducting callosal fibers, while association regions are connected by thinly myelinated slow conducting callosal fibers (LaMantia and Rakic, 1990; Aboitiz et al., 1992). However, we note that these fibers cannot fully explain the strong inter-hemispheric coupling in early visual regions given that callosal fibers terminate preferentially in regions representing the vertical midline of the visual field (Van Essen and Zeki, 1978; Van Essen et al., 1982). Instead, homotopic correlations (and RSFC in general) are mediated by indirect synaptic coupling (Vincent et al., 2007; Stark et al., 2008; Lu et al., 2011; Xue et al., 2021).

Previous studies (e.g., Yeo et al., 2011) have demonstrated that somatomotor regions representing distal limbs (e.g., hand and foot) exhibited stronger homotopic correlations than somatomotor regions representing midline regions (e.g., tongue), which is consistent with non-human primate studies showing denser callosal fibers connecting regions representing midline structures (Pandya and Vignolo 1971; Jones and Wise 1977; Killackey et al., 1983; Gould et al., 1986). It is also well-known that finger or foot movements activate contralateral somatomotor regions while tongue movements activate bilateral tongue regions (Ehrsson et al., 2003). Therefore, it is perhaps not surprising that homotopic parcels exhibiting stronger laterality during the motor task also exhibited weaker homotopic correlations (Figure 9). Similar results were obtained during the language task: homotopic parcels exhibiting stronger laterality during the language task also exhibited weaker homotopic correlations (Figure 8).

Overall, the hMRF parcels replicated well-known homotopic properties of the cerebral cortex during resting and task states. The hMRF parcellations also exhibited stronger homotopic RSFC than other homotopic parcellations (Joliot et al., 2015; Glasser et al., 2016). Therefore, we expect the hMRF parcellations to be a useful dimensionality reduction tool for future brain lateralization studies in health and disease. Finally, hMRF and Schaefer parcellations exhibit highly matched properties in terms of architectonic alignment, visuotopic alignment, task inhomogeneity and resting homogeneity. Therefore, the hMRF parcellations can also be utilized in studies of brain organization unrelated to lateralization.

### 4.2 hMRF parcels align with cortical areal boundaries

The local gradient approach for parcellating the cerebral cortex detects abrupt changes in RSFC patterns (Cohen et al., 2008; Laumann et al., 2015; Gordon et al., 2016). On the other hand, the global similarity approach clusters voxels (or vertices) with similar functional connectivity patterns (van den Heuvel et al., 2008; Power et al., 2011; Yeo et al., 2011; Ryali et al., 2013; Gordon et al., 2017). Studies have suggested that the local gradient approach can detect certain areal boundaries not revealed by the global similarity approach (Wig et al., 2014; Buckner and Yeo, 2014).

We previously proposed the gwMRF model to integrate local and global parcellation approaches (Schaefer et al., 2018). The resulting 400-region Schaefer parcellation aligned well with certain areal boundaries, while exhibiting excellent resting homogeneity and task inhomogeneity. In the current study, the proposed hMRF model extended the gwMRF model by encouraging homotopic parcels. One concern was that the additional homotopic constraint might lead to parcellations with worse areal alignment. However, across both histological and visuotopic areas, we found that the hMRF parcellations exhibited comparable areal alignment with the Schaefer, Craddock, Shen and AICHA parcellations (Craddock et al., 2012; Shen et al., 2013; Joliot et al., 2015; Schaefer et al., 2018) and better areal alignment that the Gordon parcellation (Gordon et al., 2016). Compared with the semi-manual multimodal Glasser parcellation (Glasser et al., 2016), the hMRF parcellation exhibited comparable histological areal alignment, but worse visuotopic areal alignment (see further discussion in Sections 4.6 and 4.7).

Similar to the Schaefer parcellation, the 400-region hMRF parcellation aligned well with certain cortical areas, e.g., the boundary of histological area 17 (and visuotopic V1), as well as the boundary between histological areas 3 and 4 along the central sulcus. However, the hMRF parcellation did not align well with all cortical areas. It is worth noting that for some areas, the architectonic and retinotopic boundaries are themselves not perfectly aligned. For example, histological area 18 is not well aligned to visuotopic areas V2 or V3. Therefore, it is not possible for the hMRF parcellation to align well with both histological and visuotopic boundaries.

Similar to the Schaefer parcellation, the 400-region hMRF parcellation also captured subareal features. For example, the hMRF parcellation appeared to fractionate somatosensory and motor areas based on the different body representations, as well as visual regions based on their eccentricity representation. This was particularly clear for area 17 or V1 (Figure 1), but likely extended to other visuotopic areas. As mentioned in the introduction, we believe that this subareal characteristic can be potentially useful in many applications. For example, differentiation of the hand and tongue regions might be useful when modeling a task involving button presses. Similarly, many tasks require participants to fixate on certain visual stimuli, so we expect differential response between central (low eccentricity) and peripheral (high eccentricity) visual regions.

### 4.3 hMRF parcels are homogeneous during resting and task states

Brain parcellations are often used for dimensionality reduction in MRI analyses (Eickhoff et al., 2018b). For example, resting-state or task-state fMRI time courses within each brain parcel are often averaged to compute function connectivity matrices, which are in turn used for studying mental disorders (Fair et al., 2013; Xia et al., 2018; Kebets et al., 2019), predicting behavioral traits (Finn et al., 2015; Greene et al., 2018; Chen et al., 2022), graph theoretic analyses (Achard et al., 2006; Wang et al., 2010; Fornito et al., 2013) and neural mass modeling (Cabral et al., 2011; Hansen et al., 2015; Kong et al., 2021b). Such a dimensionality reduction strategy only makes sense if the resting-fMRI or task-fMRI time courses are similar (i.e., homogeneous) within each parcel.

In our previous study (Schaefer et al., 2018), we demonstrated that the Schaefer parcellations exhibited better task inhomogeneity and resting homogeneity than four other public parcellations (Craddock et al., 2012; Shen et al., 2013; Glasser et al., 2016; Gordon et al., 2016). One concern is that the additional homotopic constraint in the hMRF model might weaken the homogeneity properties of the local-global parcellations. However, we found that the hMRF parcellations exhibited comparable homogeneity with the Schaefer parcellation and better homogeneity than five other parcellations (Craddock et al., 2012; Shen et al., 2013; Joliot et al., 2015; Glasser et al., 2016; Gordon et al., 2016).

Here, we evaluated task inhomogeneity using task contrasts from the HCP and ABCD datasets. We note that a strict task contrast might isolate a purer cognitive construct, but it was unlikely that any HCP or ABCD task contrast would isolate a single cognitive process.

We also did not expect that any task contrast would only activate a single cortical area. Therefore, similar to our previous study (Schaefer et al., 2018), task inhomogeneity was defined as the standard deviation of task activation z-values within each parcel. Furthermore, rather than choosing a subset of “pure” task contrasts, we utilized all unique contrasts in the HCP and ABCD datasets. Low (good) task inhomogeneity could be achieved if activation strength was uniform within each parcel. This approach did not require the task contrast to activate a single cognitive process or single parcel.

To further elaborate this logic, suppose every location within parcel X supports 30% cognitive process A, 50% cognitive process B and 20% cognitive process C. In this scenario, the parcel should still exhibit low task inhomogeneity with respect to a task contrast T that recruits 60% cognitive process A and 40% cognitive process B. Although task contrast T does not recruit the same distribution of cognitive processes as parcel X, this will not affect the task inhomogeneity of the parcel, since all locations within the parcel are affected equally. Therefore, this hypothetical example shows that the task inhomogeneity metric does not rely on the task contrasts recruiting pure cognitive processes.

As another example, suppose every location within parcel Y supports 30% cognitive process A, 50% cognitive process B and 20% cognitive process D. In this case, parcel Y should still exhibit low task inhomogeneity with respect to task contrast T from the previous paragraph. However, task contrast T will not be able to differentiate between parcels X and Y. Therefore, an implication is that although we used all available task contrasts within HCP and ABCD, these contrasts might not sufficiently differentiate among parcels. For example, none of the task contrasts likely differentiate among visual areas MT, MST and FST (Amano et al. 2009; Huk et al. 2002; Kolster et al. 2010).

A final point is that cortical areas are often functional heterogeneous, e.g., ocular dominance bands, orientation bands, and cytochrome oxidase dense “puffs” in primary visual cortex (Kaas et al., 1987). Therefore, a task contrast might only activate a portion of a cortical area. In this scenario, separating this portion of a cortical area out as a separate parcel might improve (decrease) the task inhomogeneity metric (although two other parcels will need to be fused to maintain the same number of parcels). Since our primary goal is to provide functional units for future fMRI analyses, the separation of a cortical area into more functionally uniform regions (as measured by task fMRI) can be desirable.

In the case of resting homogeneity, our evaluation utilized data across diverse acquisition protocols, preprocessing and demographics. The hMRF parcellations were estimated from young adults in the GSP dataset acquired from Siemens Tim Trio scanners (Holmes et al., 2015) whose fMRI data was preprocessed using a pipeline involving whole-brain signal regression in FreeSurfer fsaverage space (Li et al., 2019). The improvements in resting homogeneity were demonstrated in additional data from different Siemens and GE scanners with different acquisition protocols (single-band versus multiple-band), preprocessed with different pipelines (ICA-FIX versus whole-brain signal regression), in different coordinate systems (fsaverage, fsLR and MNI152) and in participants with different demographics (young adults, children and Asian populations).

Overall, our results suggest that hMRF parcellations are homogeneous during resting and task states, suggesting their potential utility for dimensionality reduction in future fMRI studies.

### 4.4 Resolution of cortical parcellation

There is no consensus on the “optimal” resolution of brain parcellations (Eickhoff et al., 2015). From the neuroscience perspective, the brain is hierarchically organized (Churchland and Sejnowski 1988; Mesulam 1998) from molecules to synapses to neurons to areas to systems. Large-scale networks comprise multiple cortical areas, while cortical areas can be again subdivided given their heterogeneity (Kaas 1987; Amunts and Zilles 2015).

Consequently, finding the “right” number of parcels is a biologically ill-posed problem. Indeed, it is unlikely that there is an optimal resolution for cortical parcellations. In fact, parcellations of different resolutions are likely useful for different applications. For example, behavioral prediction generally improves with higher dimensional parcellations before plateauing or becoming worse, although the exact results can be quite variable across studies (Dadi et al., 2019; Pervaiz et al., 2020; Kong et al., 2022). On the other hand, a recent study has suggested that brain–behavior relationships are scale-dependent (Betzel et al., 2019).

Finally, in many applications (e.g., neural mass modelling or edge-centric network analyses), higher resolution brain parcellations are often not computationally feasible, so studies often employ lower resolution parcellations (Deco et al., 2013; Hansen et al., 2015; Faskowitz et al., 2020; Kong et al., 2021b). Consequently, we generated hMRF parcellations at multiple resolutions, ranging from 100 to 1000 parcels at 100-parcel interval, which will hopefully be useful for a wide range of applications.

### 4.5 Considerations when comparing brain parcellations

An important consideration when comparing parcellations is the number of parcels. In general, greater number of parcels will lead to better evaluation metrics. Here, we controlled for the number of parcels by estimating hMRF parcellations with the same number of parcels as the benchmarked parcellations.

Another important consideration is that different parcellations are typically estimated from different datasets in different coordinate systems. Therefore, we employed multiple datasets across different coordinate systems to increase confidence in the generalizability of our parcellations to new datasets and coordinate systems. However, we note that some biases could not be fully eliminated. Projecting a parcellation from the original coordinate system (in which the parcellation was derived) will lead to some degradation in the parcellation quality. This is especially true when projecting parcellations between a surface coordinate system (fsLR or fsaverage) and a volumetric coordinate system (MNI152).

Therefore, a parcellation estimated in coordinate system X will enjoy inherent advantage over other parcellations projected to coordinate system X. For example, when using the HCP dataset in fsLR space, hMRF parcellations exhibited better resting homogeneity than the Shen, Craddock and AICHA parcellations with relatively large improvements of 8.1%, 6.4% and 5.9% respectively. On the other hand, when using the same HCP dataset in MNI152 space (with no spatial smoothing), hMRF parcellations were still statistically more homogeneous than the Shen and Craddock parcellations, but the margins were reduced to 5.5%, 3.0% and 5.0% improvements respectively.

### 4.6 Parcel size and shape

Consistent with the Schaefer parcellation, the hMRF parcellation utilized a spatial contiguity constraint encouraging brain locations within a parcel to be near to the parcel center. This is necessary to achieve spatially connected (as opposed to distributed) parcels. However, this has the side effect of encouraging rounder and more uniformly sized parcels. As discussed in our previous study (Schaefer et al., 2018), there is not a trivial way to impose a spatial contiguity constraint in “global similarity” approaches. Therefore, global similarity approaches (e.g., Craddock and Shen parcellations) generally have rounder parcels.

In terms of roundness, the hMRF parcellations were less round than the Gordon and Craddock parcellation, similar in roundness as the Shen parcellation and rounder than the AICHA, Glasser and Schaefer parcellations (Table S4). On the other hand, in terms of parcel size distribution as measured by the volumetric ratio between the 90th percentile parcel to the 10th percentile parcel, hMRF parcellations were more uniform than the AICHA, Glasser and Gordon parcellations, less uniform than the Craddock parcellation, and comparable to the Shen and Schaefer parcellations (Table S5). Therefore, compared with the other six parcellations, the hMRF parcellations were intermediate in terms of roundness and parcel size distribution.

There might be concerns that parcel roundness and size distribution might bias the homogeneity metrics in the presence of significant spatial smoothing. However, in the control analysis using HCP data in MNI152 space with no spatial smoothing, the hMRF parcellations exhibited better (greater) resting-fMRI homogeneity than the non-Schaefer parcellations. Indeed, the magnitude of improvement was greater when there was no spatial smoothing than when there was 6mm spatial smoothing. This suggests that the better resting-fMRI homogeneity exhibited by hMRF parcellations was not simply a result of rounder and more uniformly sized parcels.

From the neuroscience perspective, cortical areas can exhibit irregular size and shape distributions. In terms of size, the ratio of the largest and smallest cortical areas was roughly 200 (Van Essen et al., 2012a). In terms of shape, cortical areas can range from relatively round areas (e.g., area 44) to narrow long areas (e.g., area 3). The 400-region hMRF parcellation split V1 into visual eccentricity bands and areas 3 into somatotopic subregions. While these splits were biologically meaningful, they also increased parcel uniformity. Since V1 is one of the largest cortical areas, splitting V1 resulted in parcels with more uniform sizes. Similarly, splitting the long and thin cortical area 3 into somatotopic subregions resulted in parcels that were rounder.

The main goal of our parcellations is to provide functional units that are useful for future fMRI analysis. Therefore, the task inhomogeneity and resting-state homogeneity metrics are in some sense more important than the architectonic and visuotopic alignment metrics. Nevertheless, the architectonic and visuotopic alignment metrics do provide neurobiological insights into the cortical parcellations. The hMRF parcellation exhibited better architectonic and visuotopic alignment than the Gordon parcellation, suggesting that the hMRF parcellation might align with traditional cortical areal boundaries as well as gradient-based parcellations derived from resting-state fMRI.

Compared with the multimodal Glasser parcellation, the hMRF parcellation exhibited similar architectonic alignment and worse visuotopic alignment. This suggests that at least in the visual cortex, the Glasser parcellation aligned better with traditional cortical areal boundaries, although this likely came at the expense of worse task inhomogeneity and resting homogeneity (see discussion in Section 4.7).

Overall, given the superior task inhomogeneity and resting homogeneity, hMRF parcellations might be more useful than gradient-based parcellations when utilized as a dimensionality reduction tool for new fMRI data.

### 4.7 Limitations and future work

There are several limitations in the current study. First, the hMRF parcellations were derived by combining resting-fMRI data across many participants. However, recent studies have documented that the topography of brain networks and parcellations vary substantially across individuals, so a group-average parcellation will miss out on fine-scale individual-specific features (Laumann et al., 2015; Glasser et al., 2016; Gordon et al., 2017; Bijsterbosch et al., 2019; Kong et al., 2019; Seitzman et al., 2019). Future work will investigate the estimation of homotopic areal-level parcellations in individual participants. Similarly, the lateralization analyses in the current study were performed at the group-level. We leave individual-level analysis for future work.

Second, the hMRF parcellations were estimated only from resting-fMRI data. Traditionally, cortical areas are defined based on multi-modal criteria, including architectonics, topography, function and connectivity (Kaas, 1987; Felleman and Van Essen, 1991). In the current study, we estimated parcellations from resting-fMRI, which provides an indirect measure of brain connectivity reflecting multi-synaptic coupling (Vincent et al., 2007; Lu et al., 2011). We then validated the parcellations using histological boundaries, visuotopic boundaries, task inhomogeneity and resting homogeneity, which acted as proxies for architectonics, topography, function and connectivity respectively.

Given that the multi-modal Glasser parcellation exhibited better alignment with visuotopic boundaries, incorporating further multimodal information might further improve alignment with cortical areal boundaries. Indeed, several studies have suggested that brain networks reconfigure during tasks (Cole et al., 2014; Krienen et al., 2014) and estimates of brain parcellations are sensitive to task states (Salehi et al., 2020). Some have interpreted these results to suggest the need for separate parcellations during resting and task states (Salehi et al., 2020). However, our interpretation is that the boundaries of cortical areas (e.g., V1) should be invariant to transient task states over the span of a few days, although cortical areal boundaries can certainly be shaped during development and long-term experiences (Arcaro et al., 2017; Gomez et al., 2019). Instead, these results (Salehi et al., 2020) suggest the value of estimating areal-level parcellations from multi-modal data.

However, we note that incorporating multi-modal features can be challenging because of the need to select among potentially conflicting information (Eickhoff et al., 2018a). In the case of the semi-manual Glasser parcellation (Glasser et al., 2016), a trained anatomist ignored strong gradients within somatomotor and visual cortices based on prior neuroanatomical knowledge. More specifically, cortical visual maps are arranged in clusters with parallel eccentricity representations (Wandell et al., 2007). For example, the eccentricity representations within visual areas V1, V2, V3, hV4, LO-1 and LO-2 are spatially contiguous (Wandell et al., 2017). Consequently, regions with the same eccentricity representations across different visual areas might share more similar resting-fMRI and task-fMRI properties than regions with different eccentricity representations within a given visual area. By selecting for visual areas, the Glasser parcellation achieved better visuotopic alignment at the expense of worse task-inhomogeneity and resting-homogeneity across all datasets, including the HCP dataset from which the Glasser parcellation was partially derived from.

As another example, the Glasser parcellation “wasted” precious parcels distinguishing between cortical areas (e.g., areas 3a and 3b), which have extremely similar resting-fMRI and task-fMRI properties. On the other hand, the Glasser parcellation did not differentiate among body representations (e.g., hand and tongue) with highly distinctive resting-fMRI and task-fMRI properties. Overall, it remains an open problem how to automatically select among competing and conflicting information across multimodal features. We leave this for future work.

Finally, the cerebral cortex forms spatially organized circuits with subcortical structures (Jones 1985; Haber 2003; Strick et al. 2009). Here, we limited our parcellations to the cerebral cortex, but there are also homotopic correspondences across subcortical structures (Saltoun et al., 2022). Our approach is in principle applicable to subcortical structures, but accurate whole-brain parcellation in a single step is nontrivial because of significant signal-to-noise difference between the cerebral cortex and subcortical structures. As such, subcortical structures might be more effectively parcellated separate from the cerebral cortex, as is done in many studies (Choi et al. 2012; Dobromyslin et al. 2012; Plachti et al., 2019; Tian et al., 2020; Xue et al., 2021). We leave the parcellation of subcortical structures for future work.

## 5 Conclusion

We develop a homotopic variant of the gwMRF local-global parcellation model, where each parcel has a spatial counterpart on the other hemisphere. The resulting parcels are homogeneous in both resting and task states across datasets from diverse scanners, acquisition protocols, preprocessing and demographics. The parcellations also agree well with a number of known visuotopic and histological boundaries, while capturing meaningful sub-areal features. Finally, the homotopic local-global parcellations replicate known homotopic and lateralization properties of the cerebral cortex. Multi-resolution homotopic local-global parcellations are publicly available as a resource for future studies (GITHUB_LINK).

## Acknowledgement

This work was supported by the Singapore National Research Foundation (NRF) Fellowship (Class of 2017), the National University of Singapore Yong Loo Lin School of Medicine (NUHSRO/2020/124/TMR/LOA), the Singapore National Medical Research Council (NMRC) LCG (OFLCG19May-0035), NMRC STaR (STaR20nov-0003), Singapore Ministry of Health (MOH) Centre Grant (CG21APR1009), and the United States National Institutes of Health (R01MH120080). This study is also supported by the National Research Foundation (NRF) under the Open Fund-Large Collaborative Grant (OF-LCG; MOH- 000504) administered by the Singapore Ministry of Health’s National Medical Research Council (NMRC) and the Agency for Science, Technology and Research (A*STAR). In RIE2025, GUSTO is supported by funding from the NRF’s Human Health and Potential (HHP) Domain, under the Human Potential Programme. Any opinions, findings and conclusions or recommendations expressed in this material are those of the authors and do not reflect the views of the Singapore NRF, MOH, A*STAR or NMRC. Xi-Nian Zuo was supported by the Startup Funds for Leading Talents at Beijing Normal University, the National Basic Science Data Center ‘Chinese Data-sharing Warehouse for In-vivo Imaging Brain’ (NBSDC-DB-15). Computational work for this article was partially performed on resources of the National Supercomputing Centre, Singapore (https://www.nscc.sg). Data were in part provided by the Human Connectome Project, WU-Minn Consortium (Principal Investigators: David Van Essen and Kamil Ugurbil; 1U54MH091657) funded by the 16 NIH Institutes and Centers that support the NIH Blueprint for Neuroscience Research; and by the McDonnell Center for Systems Neuroscience at Washington University.

Data used in the preparation of this article were obtained from the Adolescent Brain Cognitive Development^SM^ (ABCD) Study (https://abcdstudy.org), held in the NIMH Data Archive (NDA). This is a multisite, longitudinal study designed to recruit more than 10,000 children age 9-10 and follow them over 10 years into early adulthood. The ABCD Study® is supported by the National Institutes of Health and additional federal partners under award numbers U01DA041048, U01DA050989, U01DA051016, U01DA041022, U01DA051018, U01DA051037, U01DA050987, U01DA041174, U01DA041106, U01DA041117, U01DA041028, U01DA041134, U01DA050988, U01DA051039, U01DA041156, U01DA041025, U01DA041120, U01DA051038, U01DA041148, U01DA041093, U01DA041089, U24DA041123, U24DA041147. A full list of supporters is available at https://abcdstudy.org/federal-partners.html. A listing of participating sites and a complete listing of the study investigators can be found at https://abcdstudy.org/consortium_members/. ABCD consortium investigators designed and implemented the study and/or provided data but did not necessarily participate in the analysis or writing of this report. This manuscript reflects the views of the authors and may not reflect the opinions or views of the NIH or ABCD consortium investigators.

