## Supplementary file for "Homotopic local-global parcellation of the human cerebral cortex from resting-state functional connectivity"

### Supplemental Material

This supplemental material consists of Supplemental Methods and Supplemental Results to complement the Methods and Results sections in the main text.

### Supplementary Methods

This section provides the mathematical and implementation details for the hMRF parcellation procedure. Section S1 discusses the choice of surface mesh used in hMRF parcellation procedure. Section S2 specifies the mathematical details regarding the hMRF model. Section S3 summarizes how to optimize the hMRF objective function. Section S4 provides details regarding the derivations of results from Supplementary Section S2. Section S5 discusses how the hyperparameters were set for the generation of hMRF parcellations.

#### *S1. Choice of Surface Mesh*

Our parcellation procedure is performed on the Freesurfer fsaverage6 template, where each spherical hemispheric mesh consists of 40,962 vertices. Each vertex is connected to 5 or 6 neighbor's vertices via edges. Note that there is no vertex-to-vertex correspondence between the left and right hemispheres in fsaverage6 space, therefore vertex-wise correspondence was established with the FreeSurfer command *mris_left_right_register*.

#### *S2. Mathematical Model*

In this section, we discuss the intuition behind the hMRF model. The fMRI time courses of each participant at each vertex were normalized to be of zero mean and standard deviation of one. The GSP dataset consists of participants with one or two runs. For participants with two runs, each run was normalized separately. For each vertex, the time courses of all runs of all participants were concatenated into a long column vector and normalized into unit norm. The concatenated time course at vertex $n$ is denoted as $y_n$. This normalization and concatenation approach is motivated by the fact that the inner product of two concatenated time courses is equivalent to computing the Pearson product-moment correlation for each participant and then averaging across all participants (for more details, see Supplementary Methods S3 of Schaefer et al., 2018).

Let $N$ denote the total number of vertices. Our input data thus consists of the normalized and concatenated time courses $\{y_1, \ldots, y_N\}$, denoted briefly as $y_{1:N}$ (each of length $D$), and the 3-dimensional spherical coordinates $s_{1:N}$. Our goal is to estimate the cortical labels for each vertex $l_{1:N} = \{l_1, \cdots, l_N\}$, where $l_n \in \{1, \ldots, L\}$ and $L$ indicates the desired number of parcels. The following model describes the joint distribution of labels $l_{1:N}$, spatial coordinates $s_{1:N}$, and the fMRI time courses $y_{1:N}$:

$$p(l_{1:N}, y_{1:N}, s_{1:N}) \\ = \frac{1}{Z}\exp\{ -U_{global}(l_{1:N}, y_{1:N}) - V_{grad}(l_{1:N}) - U_{xyz}(l_{1:N}, s_{1:N}) - V_{homo}(l_{1:N}) - U_{sep}(l_{1:N})\}, \quad (1)$$

where $Z$ is the normalization term to ensure that the resultant term is a valid probability distribution. $U_{global}(l_{1:N}, y_{1:N})$ is the unary potential encoding the global similarity of fMRI time courses. $V_{grad}(l_{1:N})$ defines the pairwise potential incorporating the local gradient approach. $U_{xyz}(l_{1:N}, s_{1:N})$ is the unary potential encoding the spatial contiguity of individual parcels. $V_{homo}(l_{1:N})$ is the pairwise potential encoding the homotopic constraint. $U_{sep}(l_{1:N})$ is a bookkeeping unary potential to keep the left hemisphere parcels on the left hemisphere and the right hemisphere parcels on the right hemisphere.

We note that $U_{global}(l_{1:N}, y_{1:N})$, $V_{grad}(l_{1:N})$, and $U_{xyz}(l_{1:N}, s_{1:N})$ were the same the original local-global parcellation approach (Schaefer et al., 2018). More specifically, the global similarity potential is written as

$$U_{global}(l_{1:N}, y_{1:N}) \\ = -\sum_{n=1}^{N} \log\, p(y_n | l_n;\, \kappa_{l_n}, \mu_{l_n}) \;= -\sum_{n=1}^{N} \kappa_{l_n} y_n^T \mu_{l_n} - \sum_{n=1}^{N} \log z(\kappa_{l_n}), \quad (2)$$

where $p(y_n | l_n;\, \kappa_{l_n}, \mu_{l_n})$ is a von Mises-Fisher distribution. $\kappa_{l_n}$ and $\mu_{l_n}$ are the concentration parameter and mean direction respectively for the von Mises-Fisher distribution associated with parcel $l_n$. In practice, as can be seen in Supplementary Methods S3, $\mu_{l_n}$ is iteratively estimated and corresponds to the mean time course of parcel $l_n$ normalized to unit norm. Therefore, if the time course of vertex $n$ is similar to the mean time course of parcel $l_n$ (i.e.,

$y_n^T \mu_{l_n}$ is large), then the cost of assigning vertex $n$ to parcellation label $l_n$ will be smaller. Overall, the global similarity potential encourages vertices with similar time courses to be assigned to the same parcel.

The local gradient potential is written as

$$V_{grad}(l_{1:N}) = \sum_{n=1}^{N} \left( \sum_{m \in N_{q_n}} \Delta(l_n, l_m) * c * (e^{-kGrad(q_n, q_m)} - e^{-k}) \right), \quad (3)$$

where $c$ and $k$ are constants, $N_{q_n}$ denotes the neighboring vertices of a vertex $q_n$ on the surface mesh and the delta function

$$\Delta(l_n, l_m) = \begin{cases} 1 & l_n \neq l_m \\ 0 & l_n = l_m \end{cases} \quad (4)$$

Therefore, the delta function penalizes neighboring vertices with different parcellation labels. The delta function is multiplied by the gradient-weighted exponential term $e^{-kGrad(q_n, q_m)} - e^{-k}$. Here, $Grad(q_n, q_m)$ is the magnitude of RSFC gradient between vertices $q_n$ and $q_m$ and its value ranges from 0 to 1. Consistent with our previous study (Schaefer et al., 2018), we utilized the gradient map from a previous study (Gordon et al., 2016). If $l_n = l_m$ (i.e., vertices $n$ and $m$ have the same parcellatin label), then the penalty is always zero. If $l_n \neq l_m$ (i.e., vertices $n$ and $m$ have different parcellation labels), the penalty is weighted by the exponential term $e^{-kGrad(q_n, q_m)} - e^{-k}$. As $Grad(q_n, q_m)$ increases from zero to one, the exponential term decays from $(1 - e^{k})$ to zero. Therefore, the penalty goes to zero if there is a very strong local RSFC gradient. $k$ controls the decay rate of the exponential function and $c$ controls for weight of the terms relative to other terms in the model. Both $k$ and $c$ are hyperparameters that are manually set.

Similarly, we utilize the von Mises-Fisher distribution for the spatial contiguity term:

$$U_{xyz}(l_{1:N}, s_{1:N}) = -w_{xyz} * \sum_{n=1}^{N} \log\, p(s_n | l_n;\, v_{l_n}, \tau_{l_n})$$

$$= -w_{xyz} * (\sum_{n=1}^{N} \tau_{l_n} s_n^T v_{l_n} + \sum_{n=1}^{N} \log z(\tau_{l_n})), \qquad (5)$$

where $s_n$ is the 3D coordinates of the $n$-th vertex on the fsaverage6 sphere. $\tau_{l_n}$ and $v_{l_n}$ are the concentration parameter and mean direction respectively for the von Mises-Fisher distribution associated with parcel $l_n$. In practice, as can be seen in Supplementary Methods S3, $v_{l_n}$ is iteratively estimated and corresponds to the mean spatial coordinates of parcel $l_n$ normalized to unit norm (i.e., sphere). Therefore, if the spatial coordinates of vertex $n$ is similar to the mean spatial coordinates of parcel $l_n$ (i.e., $s_n^T v_{l_n}$ is large), then the cost of assigning vertex $n$ to parcellation label $l_n$ will be smaller. Here $w_{xyz}$ is a tunable hyperparameter that controls the overall strength of the spatial contiguity term relative to the other terms in the cost function. The concentration parameters $\tau_{1:L}$ are also tunable. Parcels tend to get rounder as $\tau_{1:L}$ increases. Overly round parcels are not biologically realistic, so the original local-global parcellation procedure (Schaefer et al., 2018) involves an iterative procedure to adjust $\tau_{1:L}$ to ensure the spatial contiguity of individual parcels without making the parcels too round. In the current study, the procedure was simplified by initializing the model with a moderate value for $\tau_{1:L}$, and only increased the $\tau$ for spatially disconnected parcels (when such parcels emerged during the optimization procedure).

The remaining two terms $V_{homo}(l_{1:N})$ and $U_{sep}(l_{1:N})$ are necessary to achieve homotopic parcels. The homotopic pairwise potential is defined as:

$$V_{homo}(l_{1:N}) = \sum_{n=1}^{N} \left( \sum_{m \in H_{q_n}} \Delta'(l_n, l_m) \right), \qquad (6)$$

where $H_{q_n}$ is the homotopic vertex on the opposite hemisphere and

$$\Delta'(l_n, l_m) = \begin{cases} d & l_n \text{ and } l_m \text{ are homotopic parcels} \\ 0 & l_n \text{ and } l_m \text{ are not homotopic parcels} \end{cases} \qquad (7)$$

For example, in the case of the 400-region hMRF parcellation, without loss of generality, we assume that parcels 1 to 200 are in the left hemisphere and parcels 201 to 400 are in the right hemisphere. Furthermore, we assume that parcels 1 and 201 are homotopic, parcels 2 and 202 are homotopic, etc. A larger value of $d$ will result in greater penalty for homotopic vertices not being assigned to homotopic parcels and places a larger weight on this homotopic penalty term, relative to the other terms in the cost function.

Finally, because of the strong homotopic correlations between the hemispheres, we have an additional unary "bookkeeping" term that prevents parcellation labels from spanning the hemispheres.

$$U_{sep}(l_{1:N}) = -w_{sep}\left(\sum_{n=1}^{N_{hemi}} \log p_{lh}(l_n) + \sum_{n=N_{hemi}+1}^{N} \log p_{rh}(l_n)\right), \quad (8)$$

where $N_{hemi} = N/2$ and

$$p_{lh}(l_n) = \begin{cases} \frac{1}{N_{hemi}} & if\ l_n = 1,\ 2,\ \cdots, N_{hemi} \\ 0 & \text{otherwise} \end{cases} \quad (9)$$

$$p_{rh}(l_n) = \begin{cases} \frac{1}{N_{hemi}} & if\ l_n = N_{hemi}+1,\ N_{hemi}+2,\ ,\cdots, N \\ 0 & \text{otherwise} \end{cases} \quad (10)$$

To explain the above equations, let's consider the case of the 400-region hMRF parcellation, so $N_{hemi}$ = 200. Without loss of generality, we assume that parcels 1 to 200 are in the left hemisphere and parcels 201 to 400 are in the right hemisphere. Eq. (9) assigns zero probability (or infinite penalty) if a left hemisphere vertex is assigned to a right hemisphere parcel (i.e., parcels 201 to 400), while Eq. (10) assigns zero probability (or infinite penalty) if a right hemisphere vertex is assigned to a left hemisphere parcel (i.e., parcels 1 to 200). $w_{sep}$ is a hyperparameter that is simply set to a very big number to prevent any parcel from spanning across the hemispheres.

*S3. Model estimation*

Given observed spatial coordinates $s_{1:N}$, the fMRI time courses $y_{1:N}$, and fixed hyperparameters $c, d, k$, $w_{sep}$, $w_{xyz}$ and $\tau_{1:L}$, we can estimate labels $l_{1:N}$, concentration parameter $\kappa_{l_n}$, and mean directions $\upsilon_{l_n}$ and $\mu_{l_n}$using the maximum-a-posteriori (MAP) principle:

$$\underset{l_{1:N},\mu_{1:L},\kappa_{1:L},\upsilon_{1:L}}{\operatorname{argmax}} \log p(l_{1:N},\kappa_{1:L},\upsilon_{1:L},\mu_{1:L} \mid s_{1:N},\ y_{1:N},c,d,k,w_{xyz},w_{sep},\tau_{1:L}) \qquad (11)$$

We can achieve the optimization by block coordinate descent. At each iteration, given $l_{1:N}$, we can estimate $\{\kappa_{1:L},\upsilon_{1:L},\mu_{1:L}\}$, where:

$$\mu_l = \frac{\sum_{n=1}^{N} y_n \delta(l_n, l)}{||\sum_{n=1}^{N} y_n \delta(l_n, l)\,||} \qquad (12)$$

$$\kappa_l = \frac{(D-2)\Gamma_l}{2-\Gamma_l^2} + \frac{(D-1)\Gamma_l}{2(D-2)}, \text{where } \Gamma_l = \frac{1}{N}\sum_{n=1}^{N} \delta(l_n, l) y_n^T \mu_l \qquad (13)$$

$$\upsilon_l = \frac{\sum_{n=1}^{N} s_n \delta(l_n, l)}{||\sum_{n=1}^{N} s_n \delta(l_n, l)\,||}\ , \qquad (14)$$

where $\delta(l_n, l) = 1\ if\ l_n = l$ and zero otherwise. $D$ indicates the length of the time course $y_n$. $||\cdot||$ stands for $L_2$ norm. Therefore, $\mu_l$ is obtained by summing over the time courses of all vertices assigned to parcel $l$, and then normalized to unit norm; similarly, $\upsilon_l$ is obtained by summing over the spatial coordinates of vertices assigned to parcel $l$, and then normalized to unit norm.

On the other hand, given $\{\kappa_{1:L},\upsilon_{1:L},\mu_{1:L}\}$, we can use graph cuts (Delong et al., 2012) to estimate the labels $l_{1:N}$. Here we use the alpha expansion graph cut algorithm, consistent with our previous study (Schaefer et al., 2018). Given an initialization of the labels $l_{1:N}$ (and fixed values of $c, d, k,\ w_{sep},\ w_{xyz}$ and $\tau_{1:L}$), we can iterate between Eqs. (12) to (14) and the alpha expansion algorithm until convergence to estimate $l_{1:N},\kappa_{1:L},\upsilon_{1:L},\mu_{1:L}$. We refer to this iterative optimization procedure as the **MAP1** algorithm. Supplementary Section S4 provides more details about the derivation of the MAP1 algorithm.

Recall that $d$ is the hyperparameter for the homotopic constraint in Eq. (7). If $d$ is too small, then there will not be homotopic parcels. If $d$ is too big, then the homotopic parcels on both hemispheres will be identical. Therefore, we want $d$ to be just big enough so each parcel has a homotopic parcel on the other hemisphere, but not so big that the homotopic parcels are identical on both hemispheres. On the other hand, $\tau_{1:L}$ are the hyperparameters for the spatial contiguity constraint in Eq. (5). We want $\tau_{1:L}$ to be large enough so that parcels are spatially connected, but not so large that they become too round. To achieve these tradeoffs, $d$ and $\tau_{1:L}$ were set to some initial values. The MAP1 algorithm was then run until convergence. The initial value of $d$ was set to be moderately large, so that after MAP1 converged, parcels remained homotopic. We then reduced $d$ by half and re-ran the MAP1 algorithm (initializing $\{\kappa_{1:L}, \upsilon_{1:L}, \mu_{1:L}\}$ with their previous estimates). We continued reducing $d$ (and re-running the MAP1 algorithm) until parcels no longer become homotopic, in which case, we increased $d$ again. Similarly, $\tau_{1:L}$ were initialized to a moderate value. After the MAP1 algorithm converged, we increased the $\tau$ of spatially disconnected parcels and re-ran the MAP1 algorithm. This process was repeated until all parcels were spatially connected. We refer to the repeated running of MAP1 algorithm and the automatic tuning of $d$ and $\tau_{1:L}$ as the **MAP2** algorithm.

Overall, the entire algorithm proceeded as follows. For fixed values of $c, k, w_{sep}$ and $w_{xyz}$ and initial values of $d$ and $\tau_{1:L}$, we randomly initialized parcels on the left hemisphere and then reflected the parcels on the right hemisphere based on the vertex-wise homotopic correspondence obtained from mris_left_right_register. We then ran MAP2 algorithm until convergence. We repeated with different random initializations of the initial parcels. The initialization that led to the best cost function value in terms of the global similarity and local gradient terms was selected as the final estimate. We referred to this as the **MAP3** algorithm

*S4. MAP Estimation*

This section derives the MAP1 algorithm (from Section S3) in detail. Given observed spatial coordinates $s_{1:N}$, the fMRI time courses $y_{1:N}$, and fixed hyperparameters $c, d, k, w_{sep}, w_{xyz}$ and $\tau_{1:L}$, we can estimate parcellation labels $l_{1:N}$, concentration parameter $\kappa_{1:L}$, mean directions $\upsilon_{1:L}$ and mean directions $\mu_{1:L}$ using the maximum-a-posteriori (MAP) principle:

$$\underset{l_{1:N},\mu_{1:L},\kappa_{1:L},\upsilon_{1:L}}{\operatorname{argmax}} \log p(l_{1:N}, \kappa_{1:L}, \upsilon_{1:L}, \mu_{1:L} \mid s_{1:N},\ y_{1:N};\ c, d, k, w_{xyz}, w_{sep}, \tau_{1:L}) \quad (15)$$

Note that Eq. (15) is the same as Eq. (11) in Section S3. Assuming a uniform prior on $\kappa_{1:L}, v_{1:L}, \mu_{1:L}$, we can write Eq. (15) as:

$$\underset{l_{1:N},\mu_{1:L},\kappa_{1:L},v_{1:L}}{\operatorname{argmax}} \log p(l_{1:N}, s_{1:N},\ y_{1:N} | \kappa_{1:L}, v_{1:L}, \mu_{1:L}, c, d, k, w_{xyz}, w_{sep}, \tau_{1:L}) \quad (16)$$

We can then plug in Eq. (1), so the above equation becomes:

$$\underset{l_{1:N},\mu_{1:L},\kappa_{1:L},v_{1:L}}{\operatorname{argmax}} \frac{1}{Z}\exp\{ -U_{global}(l_{1:N}, y_{1:N}) - V_{grad}(l_{1:N}) - U_{xyz}(l_{1:N}, s_{1:N}) - V_{homo}(l_{1:N}) - U_{sep}(l_{1:N})\} \quad (17)$$

$Z$ is just a normalization term (that is not dependent on $l_{1:N}, \kappa_{1:L}, v_{1:L}, \mu_{1:L}$), so we can just drop $Z$ from the above maximization problem. We will also expand each term in Eq. (17) following Eq. (2) to Eq. (10), apply the log (to cancel the exp), and finally flip the sign of the maximization problem into a minimization problem.

$$\begin{aligned}\underset{l_{1:N},\mu_{1:L},\kappa_{1:L},v_{1:L}}{\operatorname{argmin}} \quad & -\left(\sum_{n=1}^{N} \kappa_{l_n} y_n^T \mu_{l_n} + \sum_{n=1}^{N} \log z(\kappa_{l_n})\right) \\ & + \sum_{n=1}^{N}\left(\sum_{m\in N_{q_n}} \Delta(l_n, l_m) * c * \left(e^{-kGrad(q_n,q_m)} - e^{-k}\right)\right) \\ & - w_{xyz} * \left(\sum_{n=1}^{N} \tau_{l_n} s_n^T v_{l_n} + \sum_{n=1}^{N} \log z(\tau_{l_n})\right) + \sum_{n=1}^{N}\left(\sum_{m\in H_{q_n}} \Delta'(l_n, l_m)\right) \\ & - w_{sep} * \left(\sum_{n=1}^{N_{lh}} \log p_{lh}(l_n) + \sum_{n=N_{lh}+1}^{N} \log p_{rh}(l_n)\right) \qquad (18)\end{aligned}$$

To optimize Eq. (18), we will alternate between estimating parcellation labels $l_{1:N}$ (while fixing $\kappa_{1:L}, v_{1:L}, \mu_{1:L}$) and estimating von-Mises distribution parameters $\kappa_{1:L}, v_{1:L}, \mu_{1:L}$ (while fixing $l_{1:N}$).

To estimate $l_{1:N}$ (while fixing $\kappa_{1:L}, v_{1:L}, \mu_{1:L}$), we note that the global minimum is NP-hard to estimate. The alpha-expansion graph cut algorithm allows the estimation of a locally optimal solution whose cost is guaranteed to be close to the global optimum (Boykov et al. 2001).

Furthermore, alpha expansion cannot be used given the negative unary terms in Eq. (18), so a large positive constant $M$ is added to offset the negative terms. The potentials were also divided by the total number of vertices $N$ to avoid numerical overflow, thus yielding the final cost function for estimating $l_{1:N}$ (while fixing $\kappa_{1:L}, v_{1:L}, \mu_{1:L}$):

$$\begin{aligned}\underset{l_{1:N}}{\operatorname{argmin}} \quad & -\frac{1}{N}\left(\sum_{n=1}^{N} \kappa_{l_n} y_n^T \mu_{l_n} + \sum_{n=1}^{N} \log z\left(\kappa_{l_n}\right)\right) \\ & + \frac{1}{N}\sum_{n=1}^{N}\left(\sum_{m \in N_{q_n}} \Delta(l_n, l_m) * c * \left(e^{-kGrad(q_n, q_m)} - e^{-k}\right)\right) \\ & - w_{xyz} * \frac{1}{N}\left(\sum_{n=1}^{N} \tau_{l_n} s_n^T v_{l_n} + \sum_{n=1}^{N} \log z\left(\tau_{l_n}\right)\right) \\ & + \frac{1}{N}\sum_{n=1}^{N}\left(\sum_{m \in H_{q_n}} \Delta'(l_n, l_m)\right) \\ & - w_{sep} * \frac{1}{N}\left(\sum_{n=1}^{N_{lh}} \log p_{lh}(l_n) + \sum_{n=N_{lh}+1}^{N} \log p_{rh}(l_n)\right) + M \end{aligned} \quad (19)$$

Alpha expansion was utilized to optimize Eq. (19) with respect to parcellation labels $l_{1:N}$.

To estimate $\kappa_{1:L}, v_{1:L}, \mu_{1:L}$ (while fixing $l_{1:N}$) is equivalent to maximum likelihood estimation of von Mises-Fisher distributions. Under the constraints that $\mu_l^T \mu_l = 1$, $v_l^T v_l = 1$ and $\kappa > 0$, we differentiate Eq. (18) with respect to $\{\kappa_{1:L}, v_{1:L}, \mu_{1:L}\}$ and set the derivatives to zero, we get (Lashkari et al. 2010):

$$\mu_l = \frac{\sum_{n=1}^{N} y_n \delta(l_n, l)}{\|\sum_{n=1}^{N} y_n \delta(l_n, l)\|} \quad (20)$$

$$\kappa_l = \frac{(D-2)\Gamma_l}{2-\Gamma_l^2} + \frac{(D-1)\Gamma_l}{2(D-2)}, \text{where } \Gamma_l = \frac{1}{N}\sum_{n=1}^{N} \delta(l_n, l) y_n^T \mu_l \quad (21)$$

$$v_l = \frac{\sum_{n=1}^{N} s_n \delta(l_n, l)}{\|\sum_{n=1}^{N} s_n \delta(l_n, l)\|}, \quad (22)$$

which are the same as Eq. (12) to Eq. (14). Overall, the MAP1 algorithm iterates between applying alpha expansion on Eq. (19) and computing Eq. (20) to Eq. (22).

*S5. Hyperparameters of the hMRF model*

This section discusses the settings of various hyperparameters in the hMRF model. For both benchmarking and the generation of a final set of parcellations, the MAP3 algorithm was ran with 1000 random initializations. As previously mentioned in Section S2, $w_{sep}$ was simply set to be a very large number. $\tau_{1:L}$ was initialized with a function to ensure that the normalization term in von Mises-Fisher distribution does not trigger numerical overflow in the Bessel function, yielding $\tau$ to be equal to 25.742.

That left us with $c$, $k$, $w_{xyz}$ and the initial value of $d$ as the true "free" hyperparameters. The settings of these hyperparameters differed when the dimensionality of the input data $y_{1:N}$, or the number of parcels $L$ changed. For the purpose of comparisons with other publicly available parcellations, the hyperparameters $c$, $k$, $w_{xyz}$ and initial value of $d$ were fixed as 80k, 15, 100k and 16,000 respectively, based on visual inspection of the parcellations estimated from the GSP training set.

For the generation of the final set of parcellations for public release, different sets of hyperparameters were used for different resolutions. Hyperparameters were selected based on visual inspection of the parcellations in the full GSP dataset. Architectonic geodesic distances, visuotopic geodesic distances and resting-homogeneity in the GSP dataset were also computed to ensure the final parcellations were both visually and quantitatively compelling. For the 100-region parcellation, $c$, $k$, $w_{xyz}$ and initial value of $d$ were set to be 10k, 15, 100k and 20k respectively. For the 200-region parcellation, $c$, $k$, $w_{xyz}$ and initial value of $d$ were set to be 110k, 15, 120k and 25k respectively. For the 300-region parcellation, $c$, $k$, $w_{xyz}$ and initial value of $d$ were set to be 130k, 15, 130k and 30k respectively. For the 400-region parcellation, $c$, $k$, $w_{xyz}$ and initial value of $d$ were set to be 140k, 15, 150k and 50k respectively. For the 500-region parcellation, $c$, $k$, $w_{xyz}$ and initial value of $d$ were set to be 180k, 15, 250k and 50k respectively. For the 600-region to 1000-region parcellations, $c$, $k$, $w_{xyz}$ and initial value of $d$ were set to be 200k, 15, 300k and 50k respectively.

**Supplementary Results**

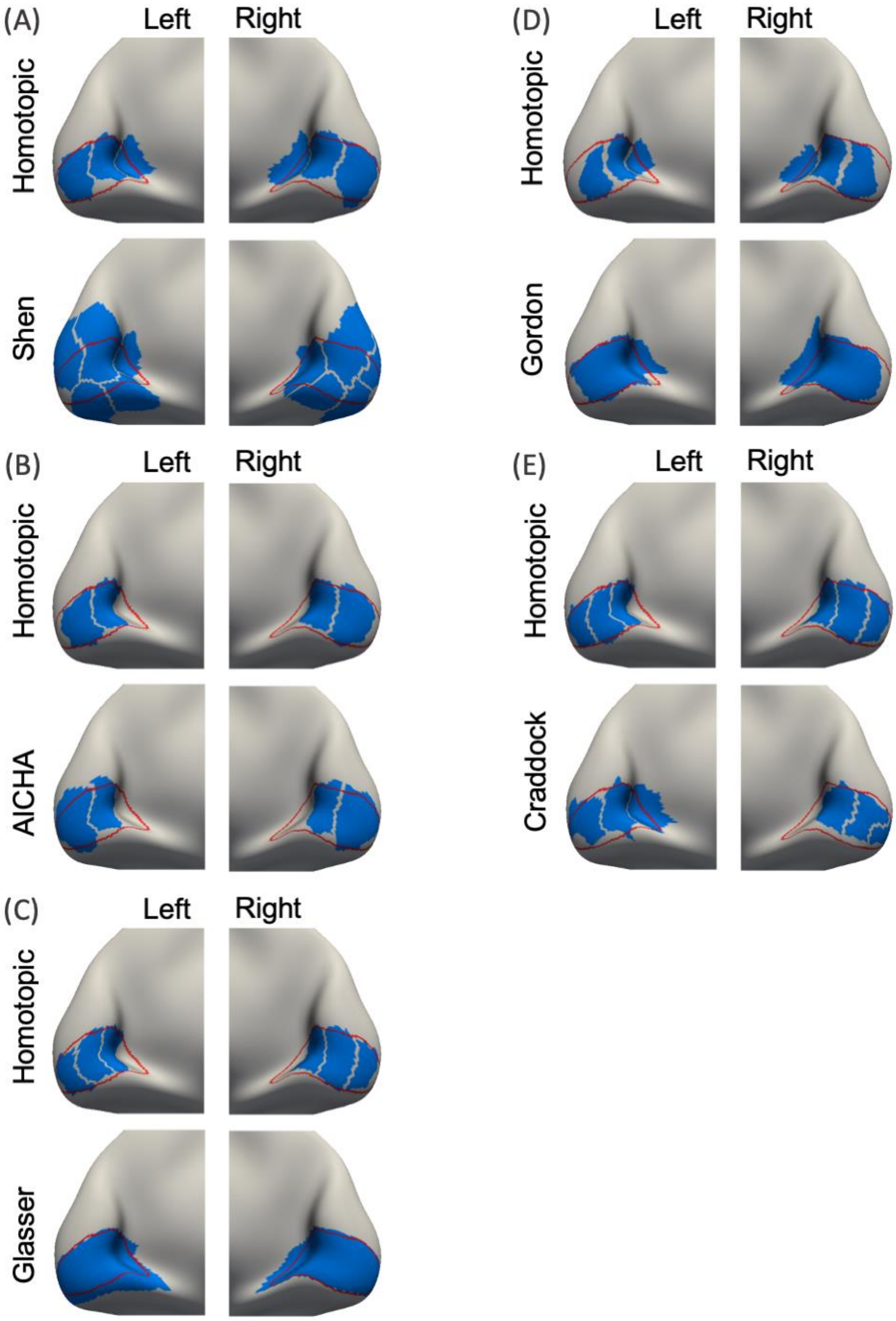


Figure S1. Comparison of homotopic correspondence between hMRF and non-Schaefer parcellations within histologically-defined area 17. Histological boundaries of area 17 are shown in red. Parcels are shown in blue.

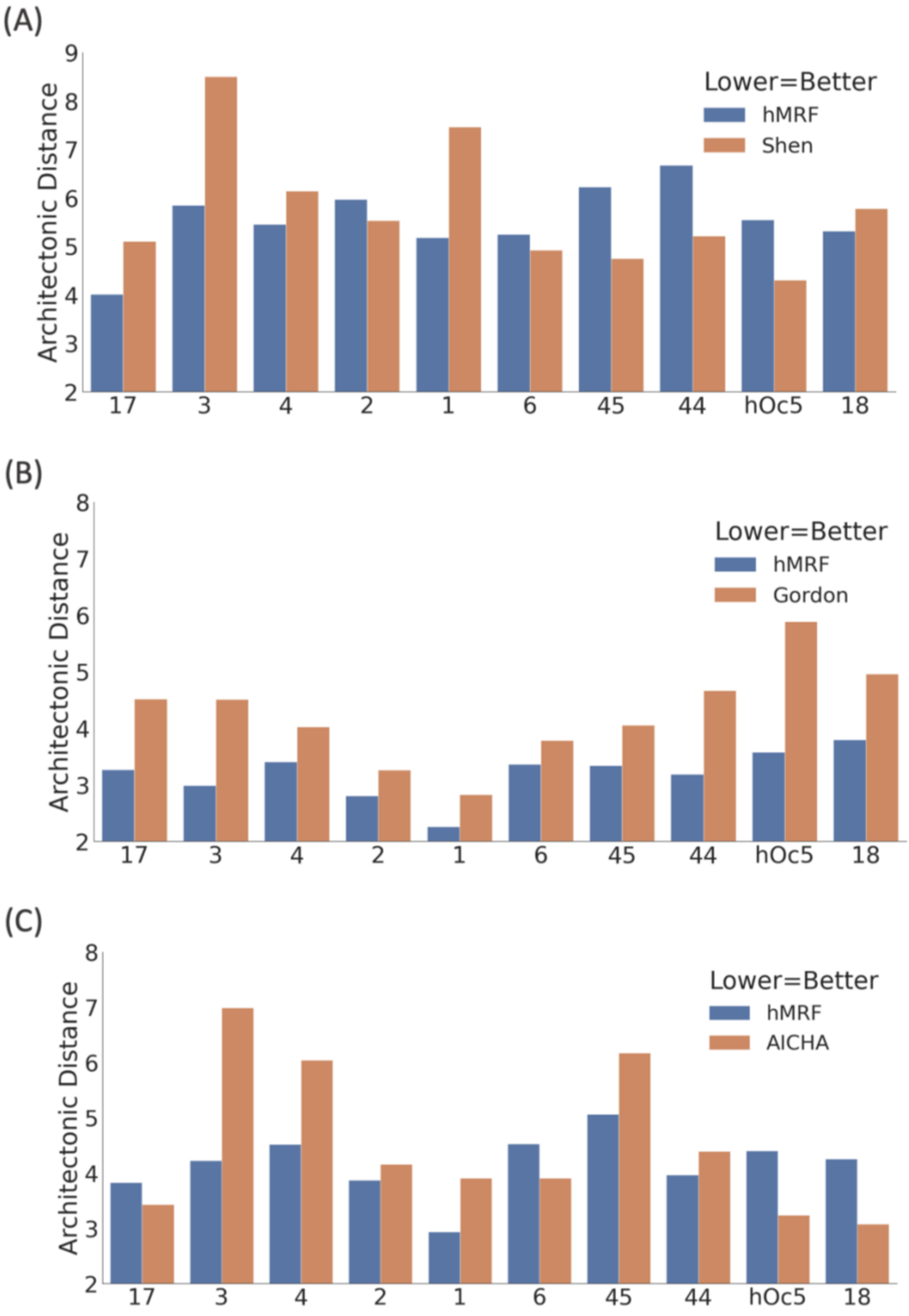


Figure S2. Distance between parcellation and architectonic boundaries as measured by average geodesic distance (mm). Lower distance indicates better agreement. Number of parcels was matched between the publicly available parcellations and corresponding hMRF parcellations. Figure continues next page.

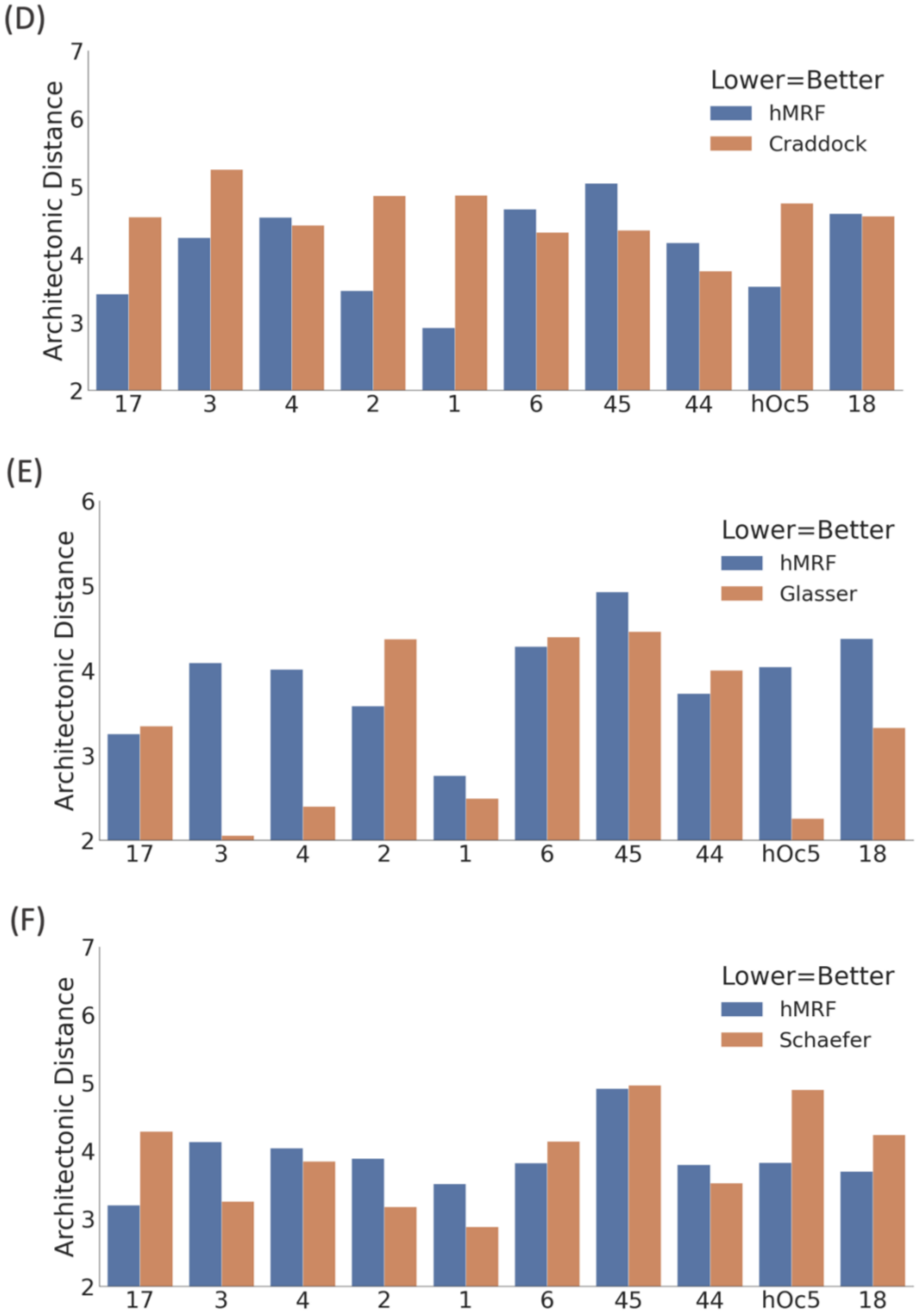


Figure S2 (cont). Distance between parcellation and architectonic boundaries as measured by average geodesic distance (mm). Lower distance indicates better agreement. Number of parcels was matched between the publicly available parcellations and corresponding hMRF parcellations. The hMRF approach generated parcellations that achieved (A) similar architectonic distance to Shen ($p = 0.441$ uncorrected), (B) better distance than Gordon ($p \approx 0$ uncorrected), (C) similar distance to AICHA ($p = 0.210$ uncorrected), (D) better distance than Craddock ($p = 0.031$ uncorrected), (E) worse distance than Glasser ($p = 0.032$ uncorrected), and (F) similar distance to Schaefer ($p = 0.835$). After correcting for multiple comparisons with a false discovery rate (FDR) of $q < 0.05$, only the comparison with the Gordon parcellation remained significant. Overall, the hMRF parcellations exhibited architectonic alignment comparable with (or better than) other parcellations. We note that the parcellations comprised 197, 333, 341, 355, 360 and 400 parcels in subplots A, B, C, D, E and F respectively. Therefore, comparisons between subplots are not meaningful because more parcels lead to more boundary vertices, and therefore lower geodesic distances (on average).

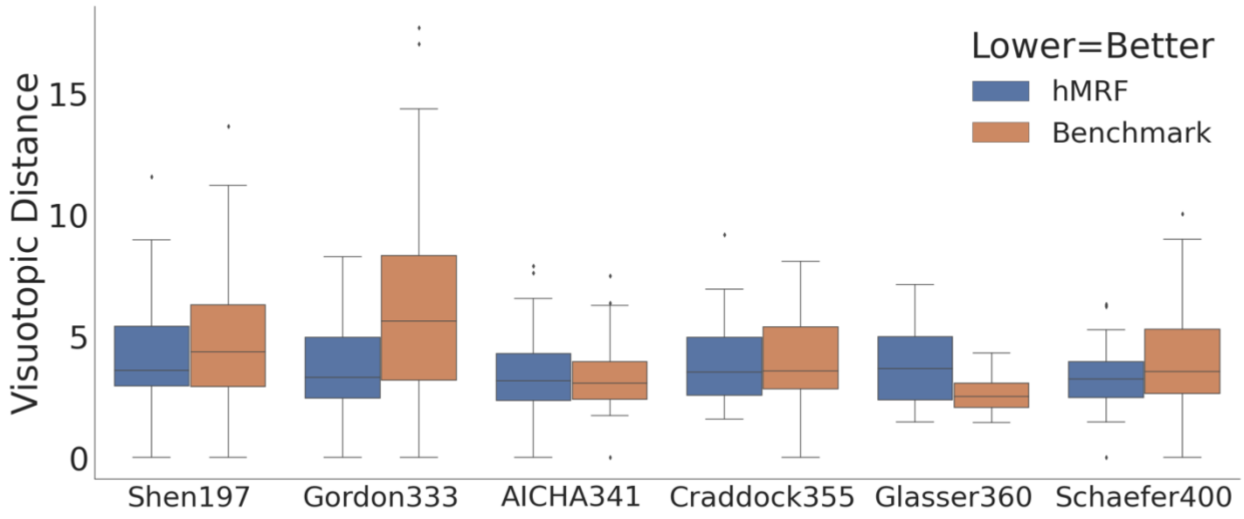


Figure S3. Distance between parcellation and visuotopic boundaries (Abdollahi et al., 2014) as measured by average geodesic distance (mm). Lower distance indicates better agreement. Number of parcels was matched between the publicly available parcellations and corresponding hMRF parcellations. The hMRF parcellations achieve (A) similar visuotopic distance to Shen ($p = 0.279$), (B) better distance to Gordon ($p \approx 0$), (C) similar distance to ACIHA ($p = 0.731$), (D) similar distance to Craddock ($p = 0.986$), (E) worse distance to Glasser ($p \approx 0$) and (F) similar distance to Schaefer ($p = 0.083$). After correcting for multiple comparisons with a FDR of $q < 0.05$, the comparisons with the Gordon and Glasser parcellations remained significant. We note that the Glasser parcellation was derived with a semi-automated algorithm that required an anatomist to manually select multi-modal information to match prior knowledge of areal boundaries. Overall, the hMRF parcellations achieved visuotopic alignment comparable with (or better than) other fully automatic approaches. Like before, comparisons between parcellations with different number of parcels (e.g., Shen197 versus Gordon333) are not meaningful.

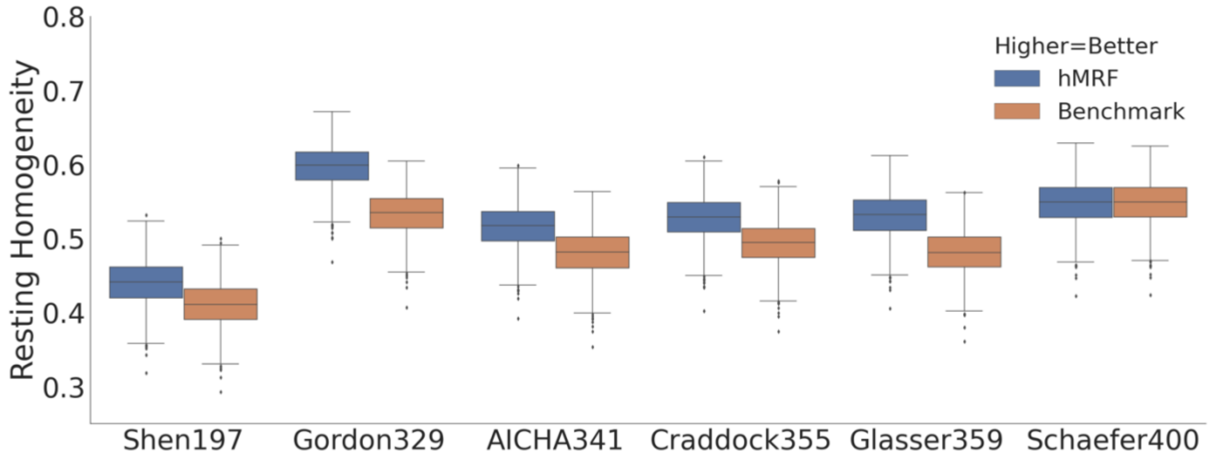


Figure S4. Resting-fMRI homogeneity computed with the GSP test set (N = 739) in fsaverage space. The hMRF parcellations achieved comparable resting-fMRI homogeneity with the Schaefer parcellation. On the other hand, the hMRF parcellations were more homogeneous than 5 parcellations.

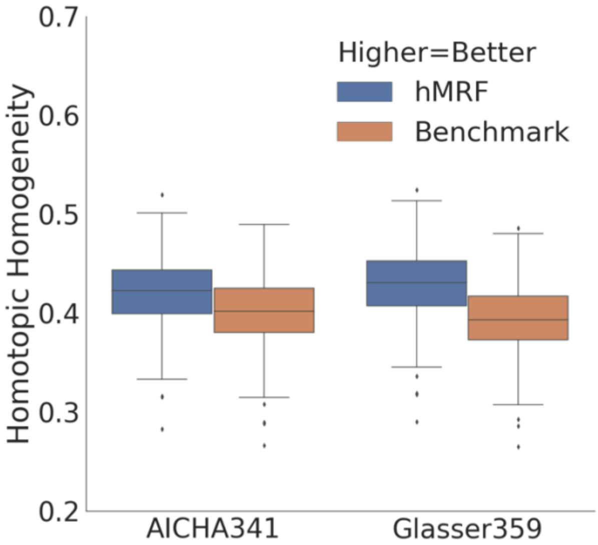

Figure S5. Homotopic resting-state functional connectivity in the GSP test set (N = 739) in fsaverage space. The hMRF parcellations achieved higher (better) homotopic resting-state functional connectivity than the Glasser and AICHA parcellations.

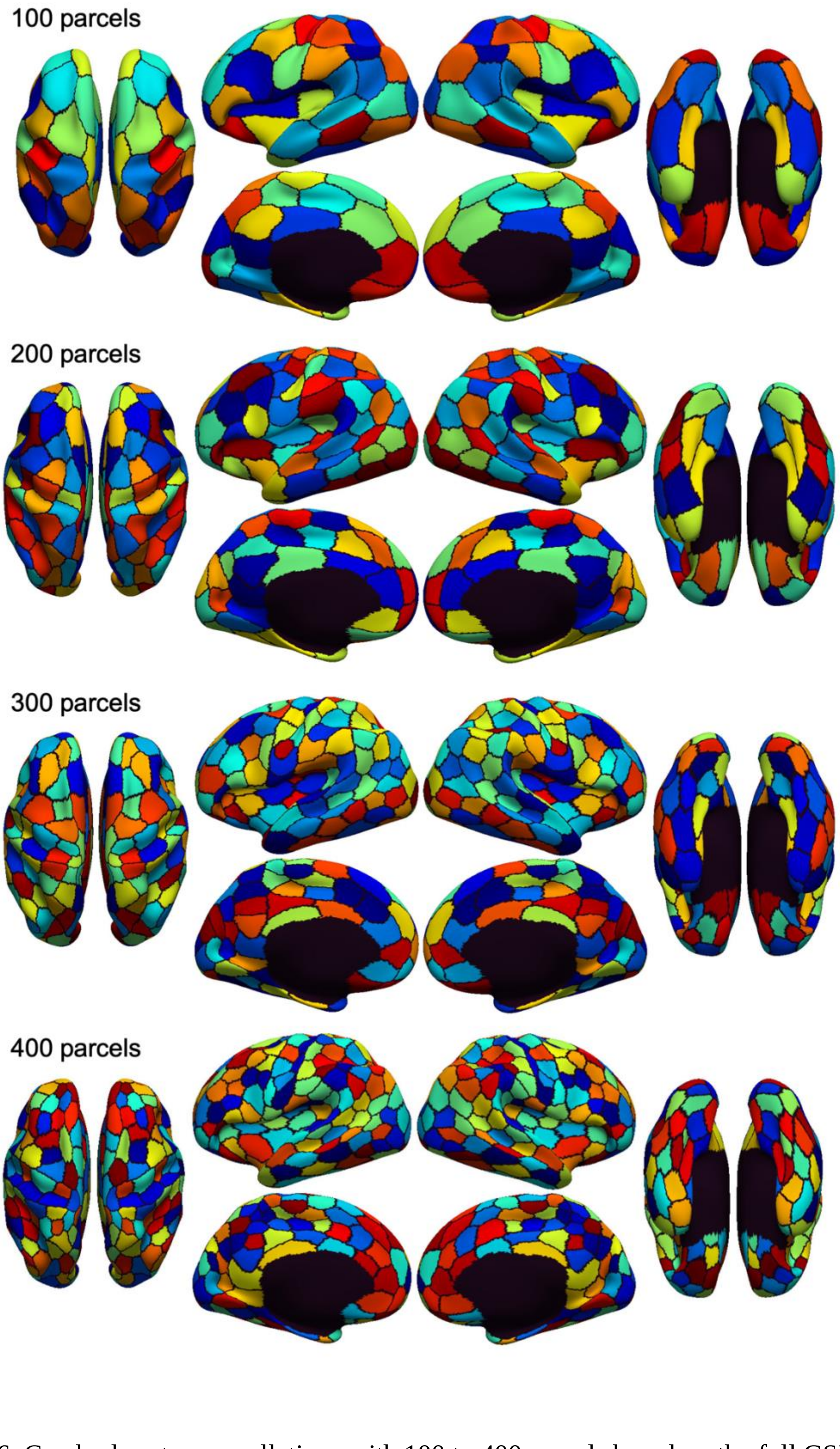


Figure S6. Cerebral cortex parcellations with 100 to 400 parcels based on the full GSP dataset of 1479 subjects. Homotopic parcels have identical colors. Figure continues next page.

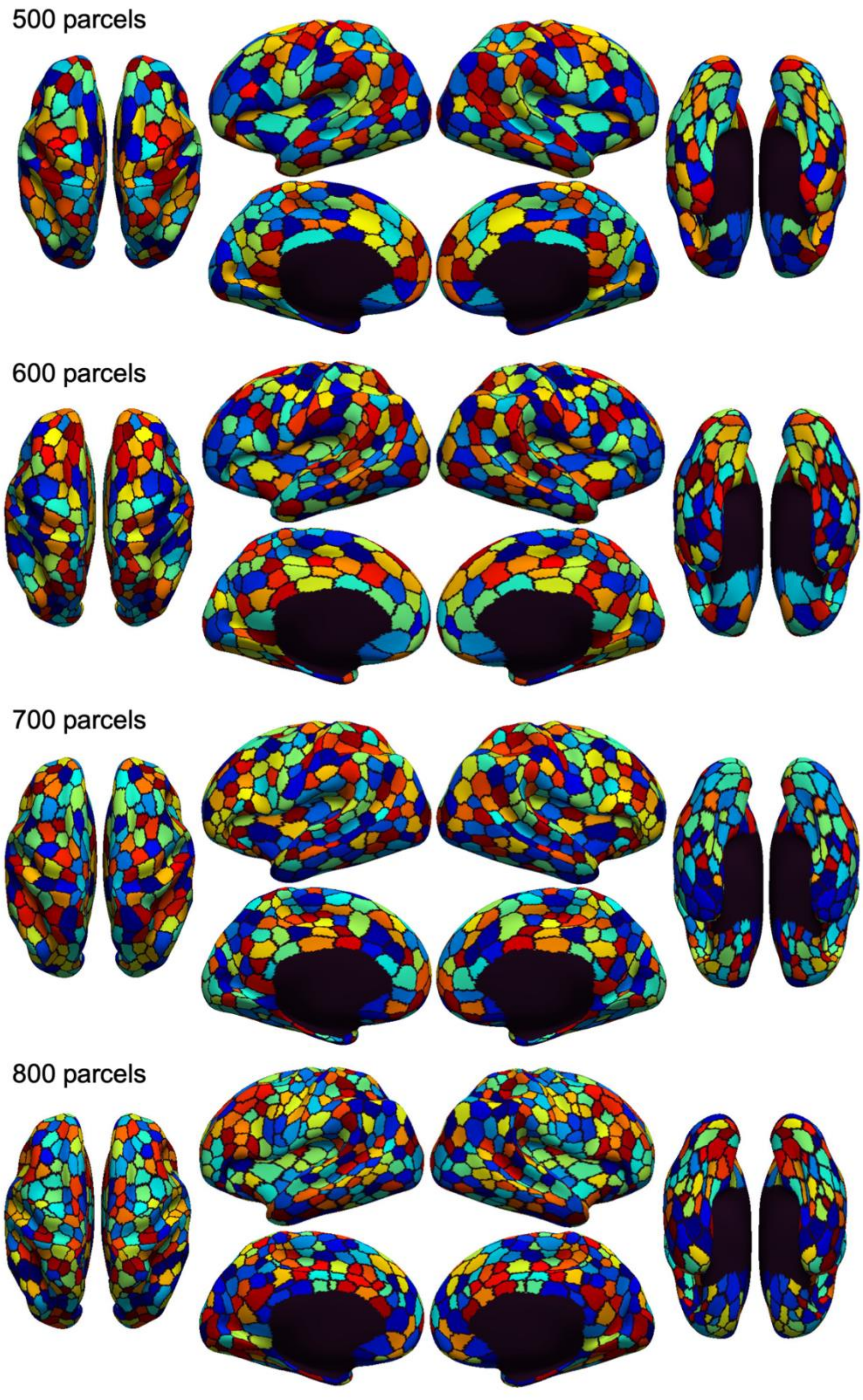


Figure S6 (cont). Cerebral cortex parcellations with 500 to 800 parcels based on the GSP dataset of 1479 subjects. Homotopic parcels have identical colors. Figure continues next page.

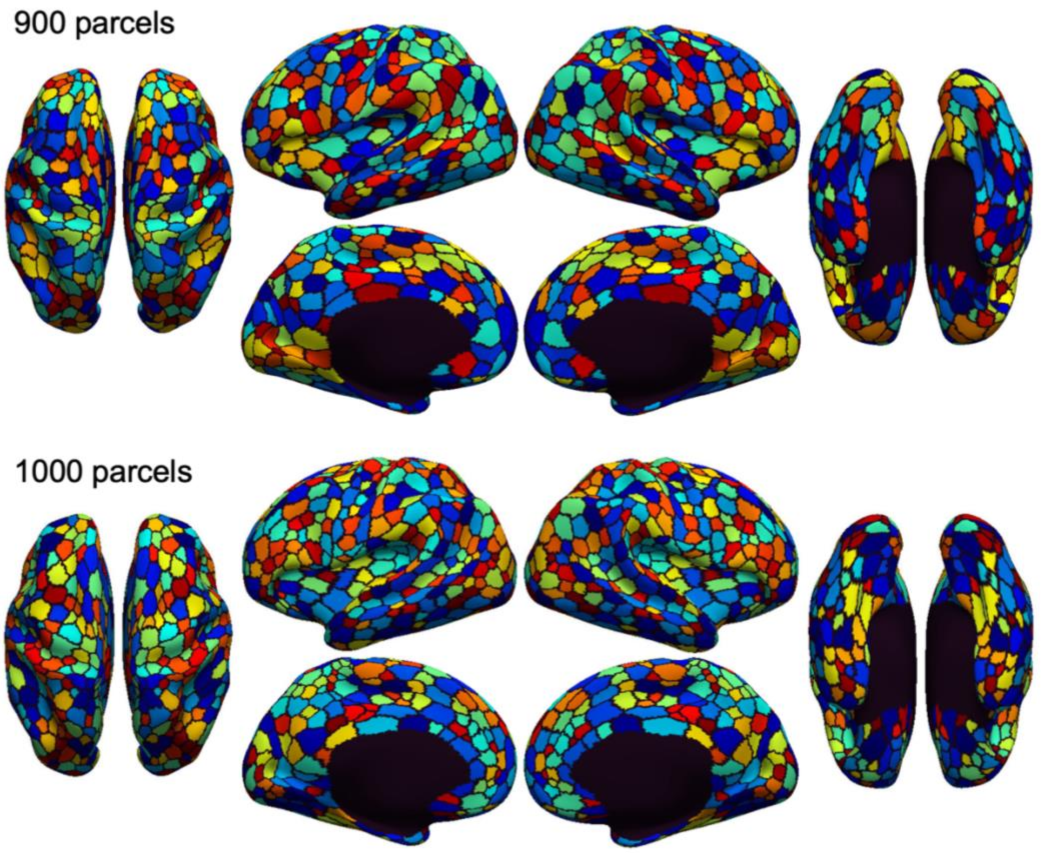


Figure S6 (cont). Cerebral cortex parcellations with 900 to 1000 parcels based on the full GSP dataset of 1479 subjects. Homotopic parcels have identical colors.

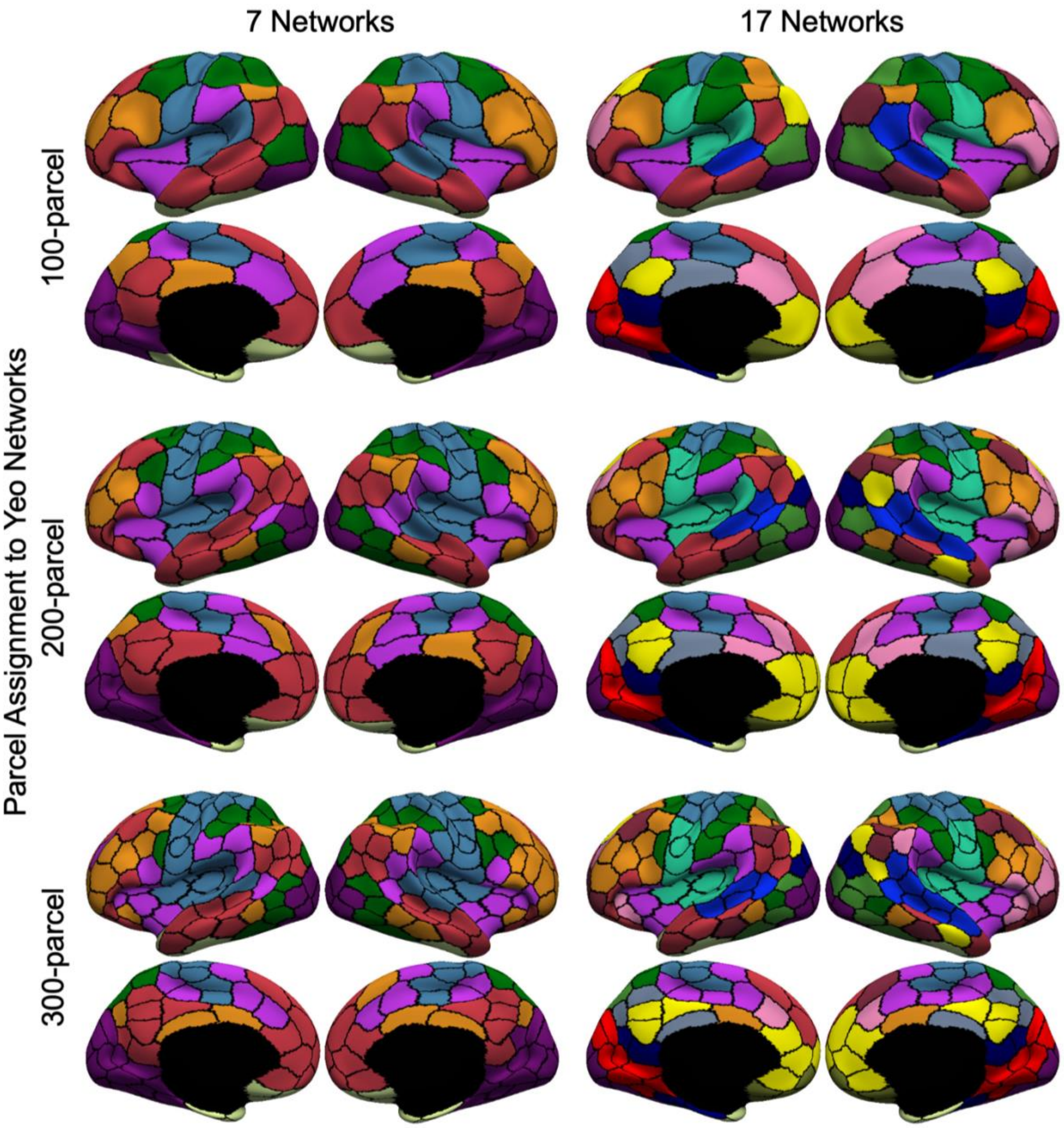


Figure S7. Assignment of parcels to 7 or 17 networks (Yeo et al., 2011) for hMRF parcellations with 100 to 300 regions. Note that the 400-region parcellation is shown in the main text (Figure 5). Figure continues next page.

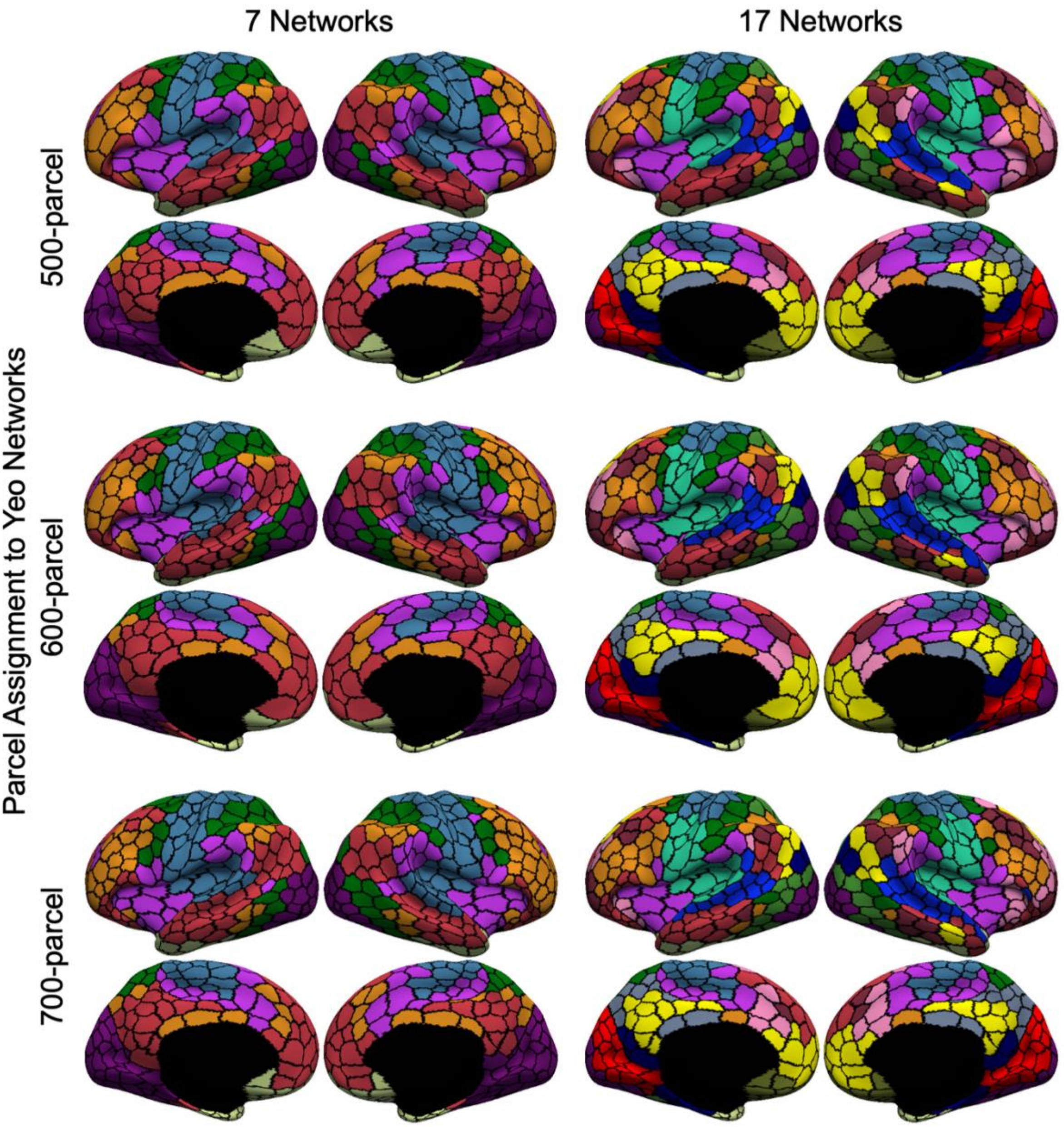


Figure S7 (cont). Assignment of parcels to 7 or 17 networks (Yeo et al., 2011) for hMRF parcellations with 500 to 700 regions. Figure continues next page.

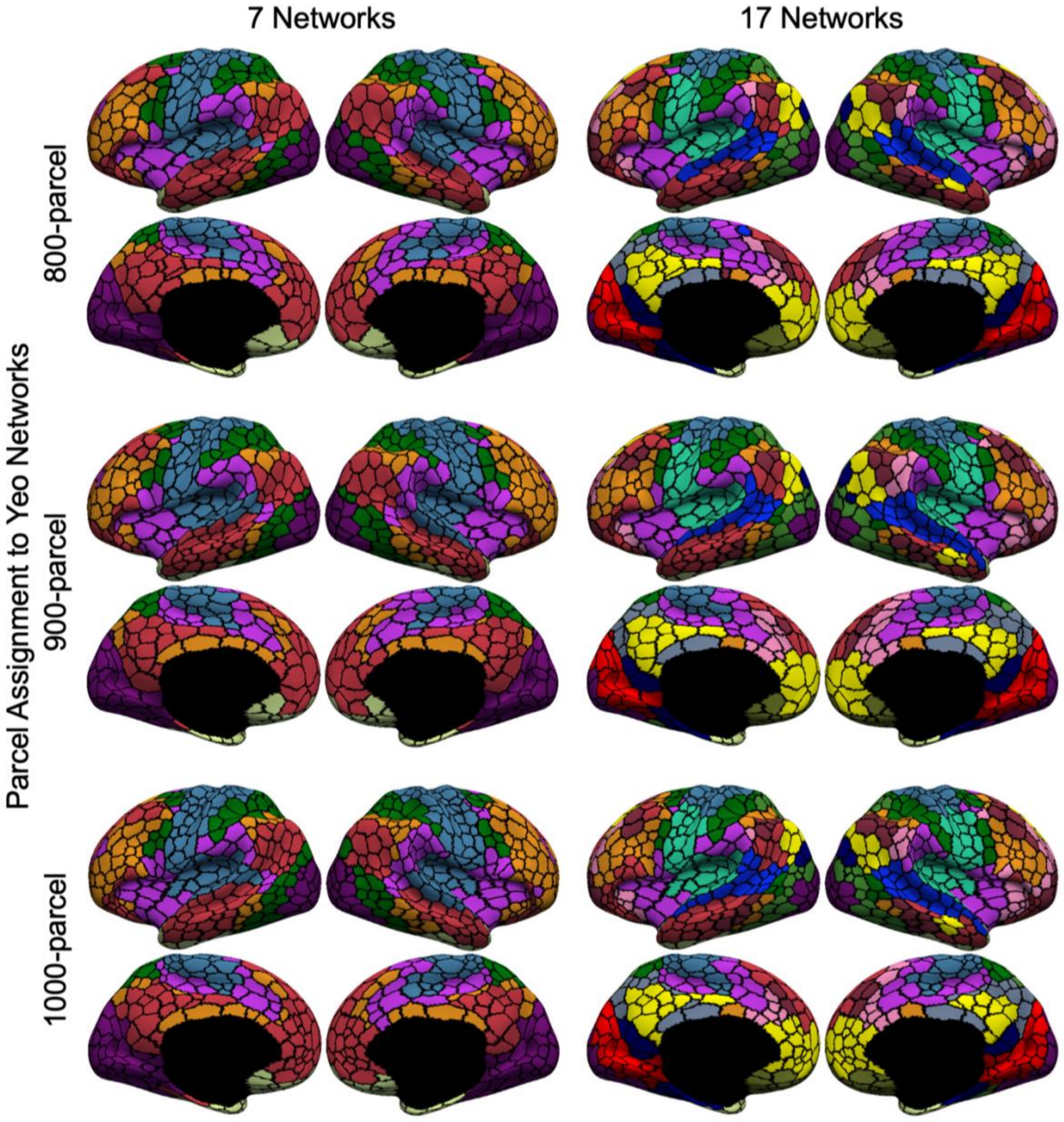


Figure S7 (cont). Assignment of parcels to 7 or 17 networks (Yeo et al., 2011) for hMRF parcellations with 800 to 1000 regions.

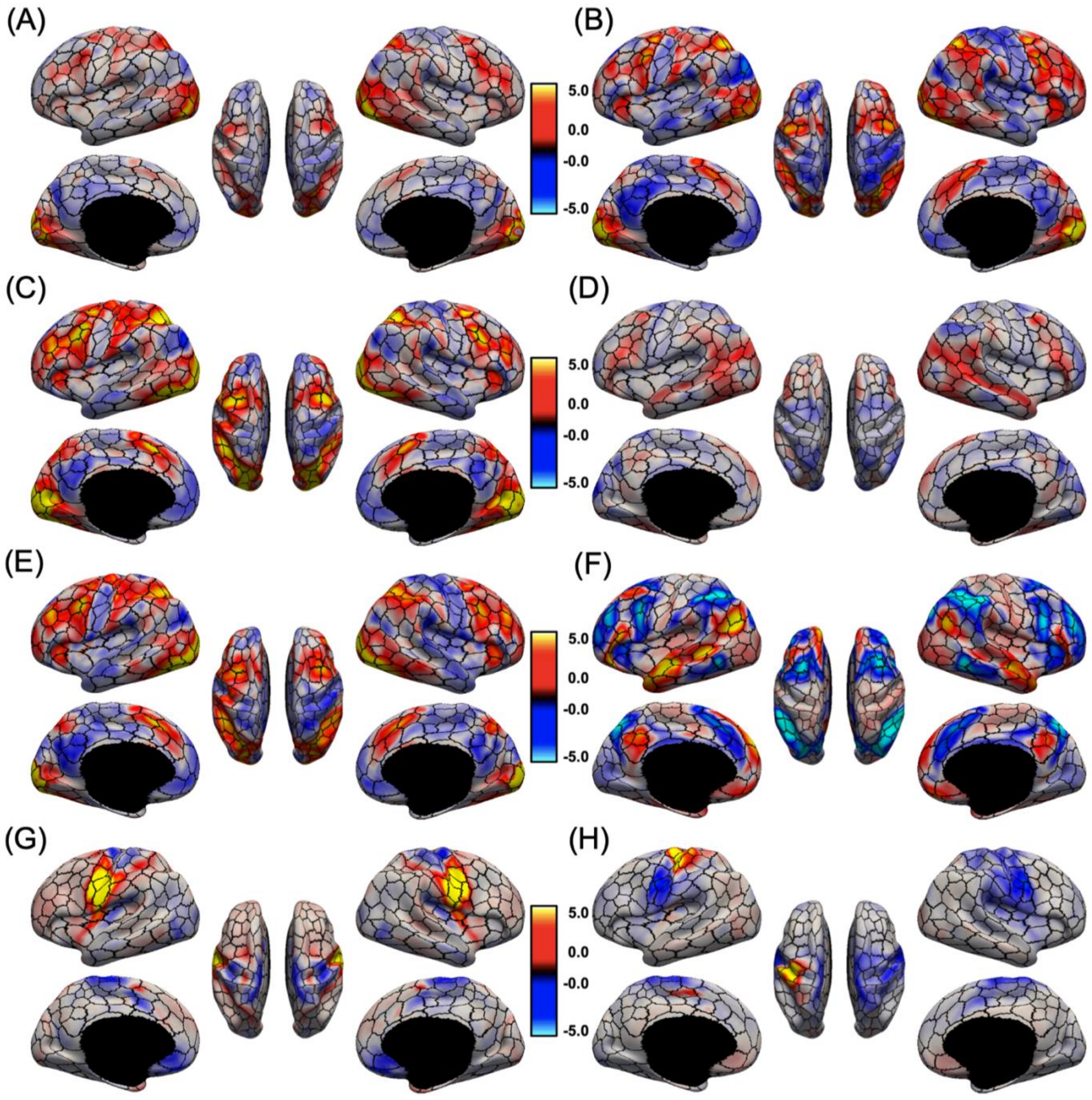


Figure S8. Group-average task activation maps of (A) emotion (faces - fixation), (B) gambling (punishment - fixation), (C) relational (matching - fixation), (D) social (theory of mind - random) and (E) working memory (2 back body - fixation), (F) language (story – math), (G) tongue motion (tongue – average motor), (H) right finger tapping (right finger – average motor) from the HCP dataset overlaid on (black) boundaries of 400-area cerebral cortex parcellation. Activations were unthresholded.

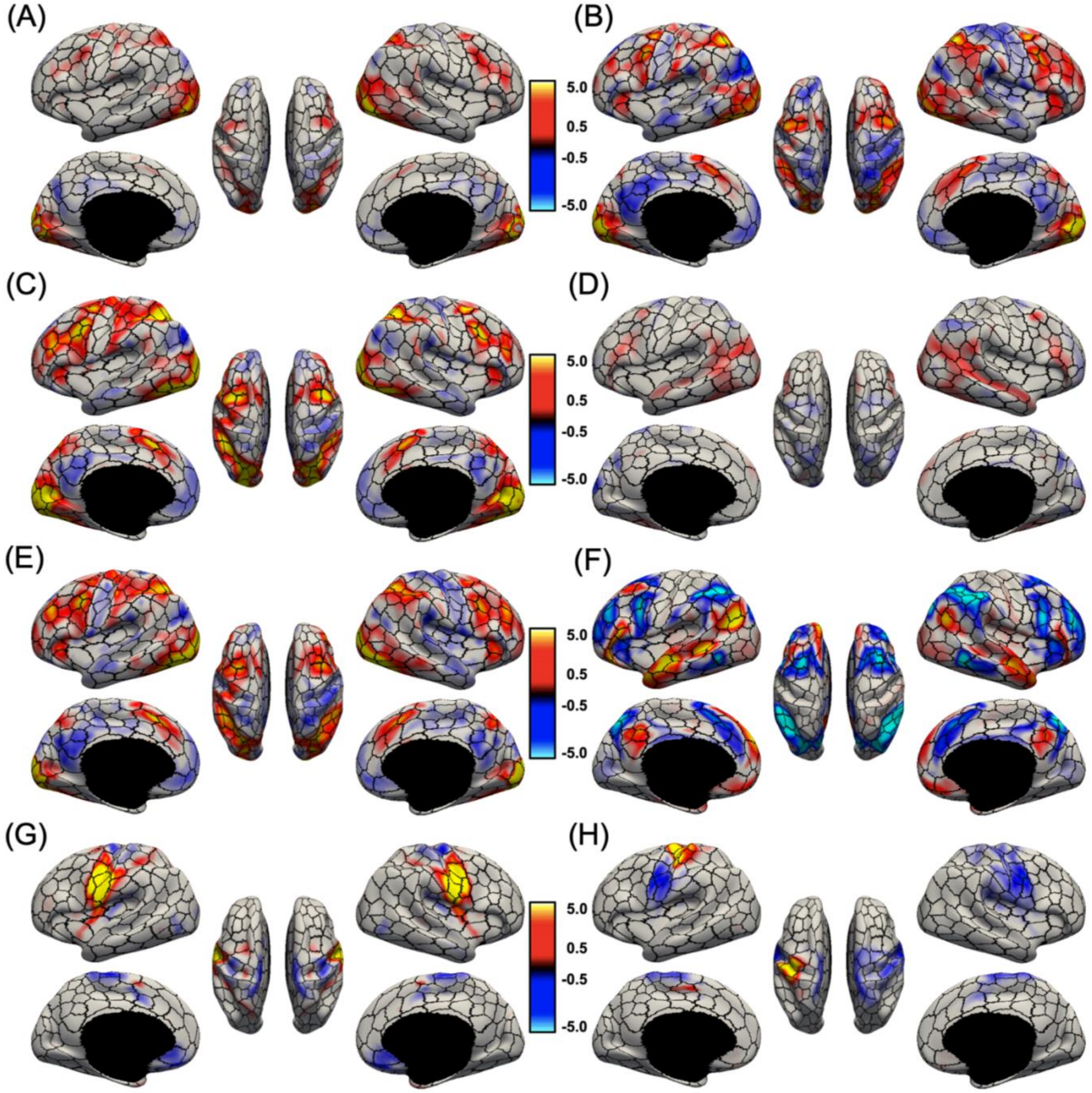


Figure S9. Group-average task activation maps of (A) emotion (faces - fixation), (B) gambling (punishment - fixation), (C) relational (matching - fixation), (D) social (theory of mind - random) and (E) working memory (2 back body - fixation), (F) language (story – math), (G) tongue motion (tongue – average motor), (H) right finger tapping (right finger – average motor) from the HCP dataset overlaid on (black) boundaries of 400-area cerebral cortex parcellation. Activations were thresholded at ±0.5.

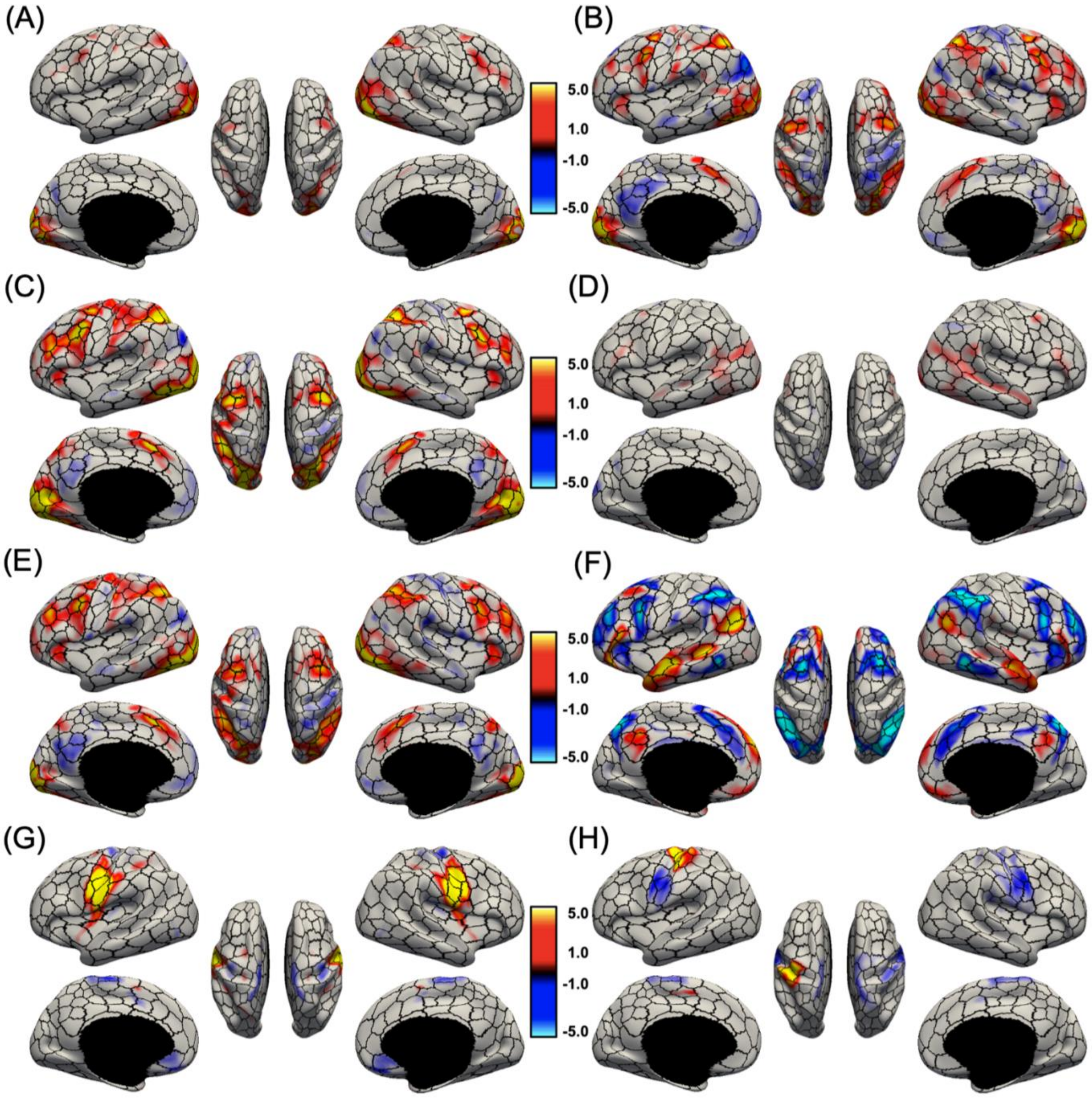


Figure S10. Group-average task activation maps of (A) emotion (faces - fixation), (B) gambling (punishment - fixation), (C) relational (matching - fixation), (D) social (theory of mind - random) and (E) working memory (2 back body - fixation), (F) language (story – math), (G) tongue motion (tongue – average motor), (H) right finger tapping (right finger – average motor) from the HCP dataset overlaid on (black) boundaries of 400-area cerebral cortex parcellation. Activations were thresholded at ±1.0.

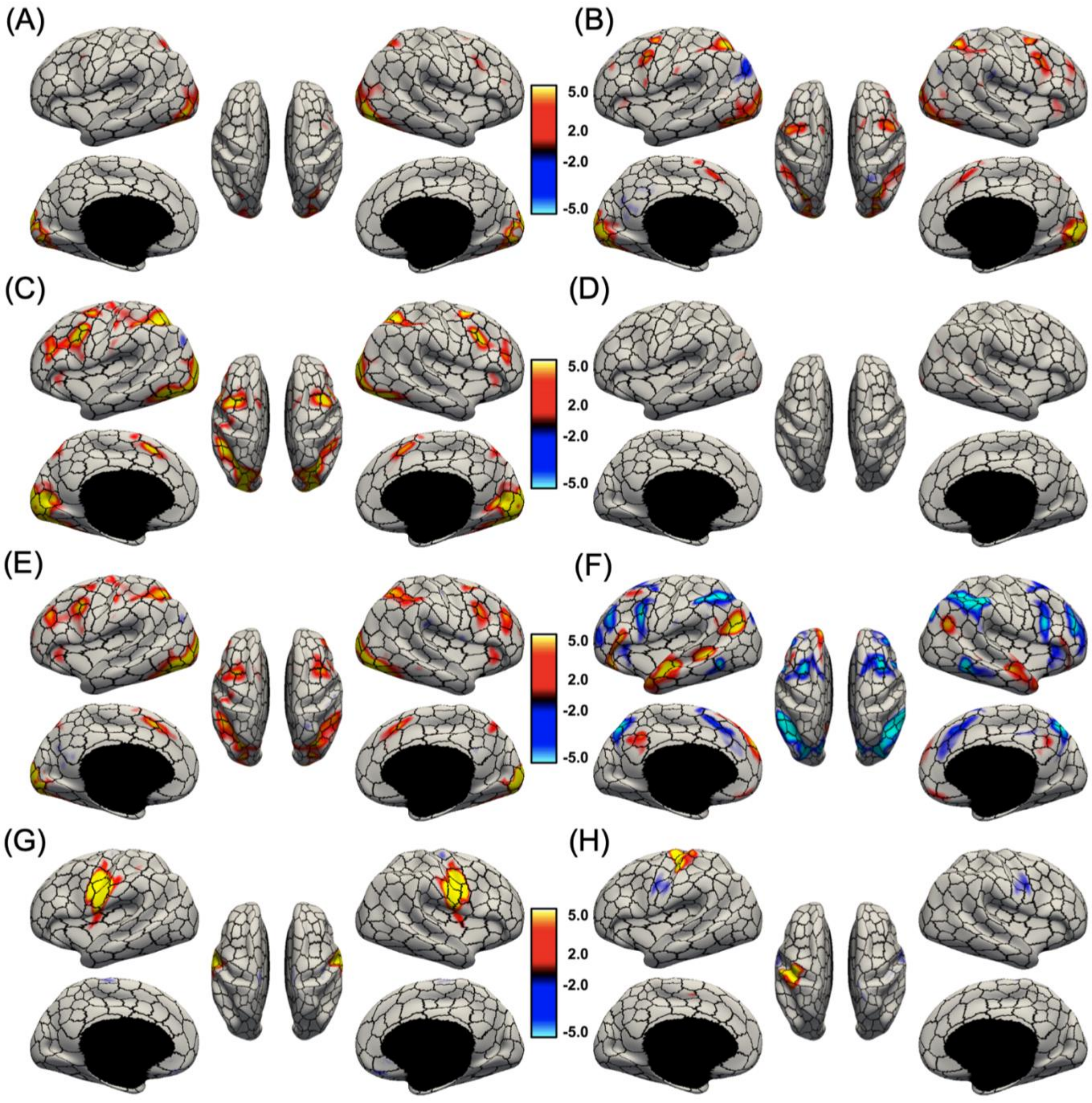


Figure S11. Group-average task activation maps of (A) emotion (faces - fixation), (B) gambling (punishment - fixation), (C) relational (matching - fixation), (D) social (theory of mind - random) and (E) working memory (2 back body - fixation), (F) language (story – math), (G) tongue motion (tongue – average motor), (H) right finger tapping (right finger – average motor) from the HCP dataset overlaid on (black) boundaries of 400-area cerebral cortex parcellation. Activations were thresholded at ±2.0.

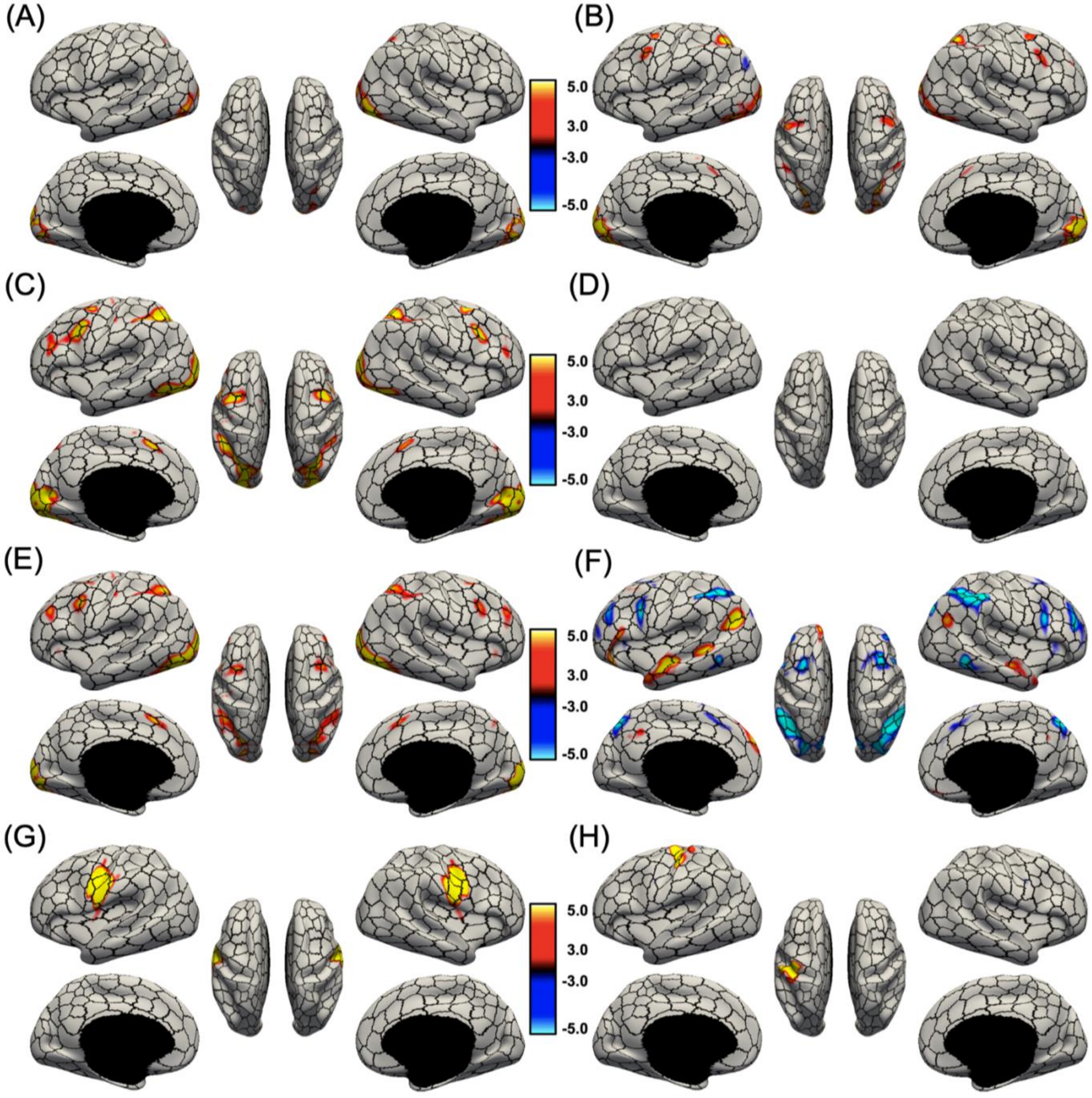


Figure S12. Group-average task activation maps of (A) emotion (faces - fixation), (B) gambling (punishment - fixation), (C) relational (matching - fixation), (D) social (theory of mind - random) and (E) working memory (2 back body - fixation), (F) language (story – math), (G) tongue motion (tongue – average motor), (H) right finger tapping (right finger – average motor) from the HCP dataset overlaid on (black) boundaries of 400-area cerebral cortex parcellation. Activations were thresholded at ±3.0.

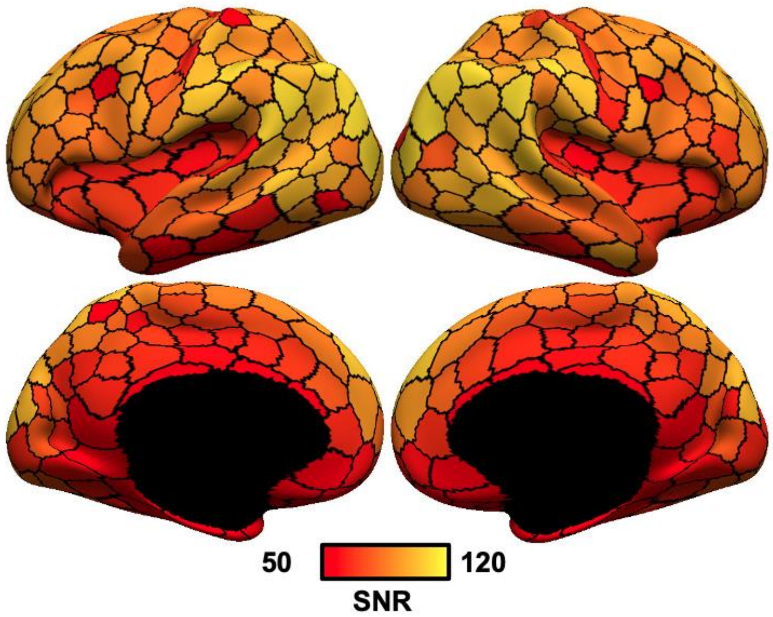


Figure S13. HCP signal-to-noise ratio (SNR) map. SNR was computed for each vertex as the mean fMRI signal divided by the standard deviation of the fMRI signal. The vertex-wise SNR map was averaged within each parcel and shown in this figure. Observe the strong anterior-to-posterior SNR gradient in the HCP dataset.

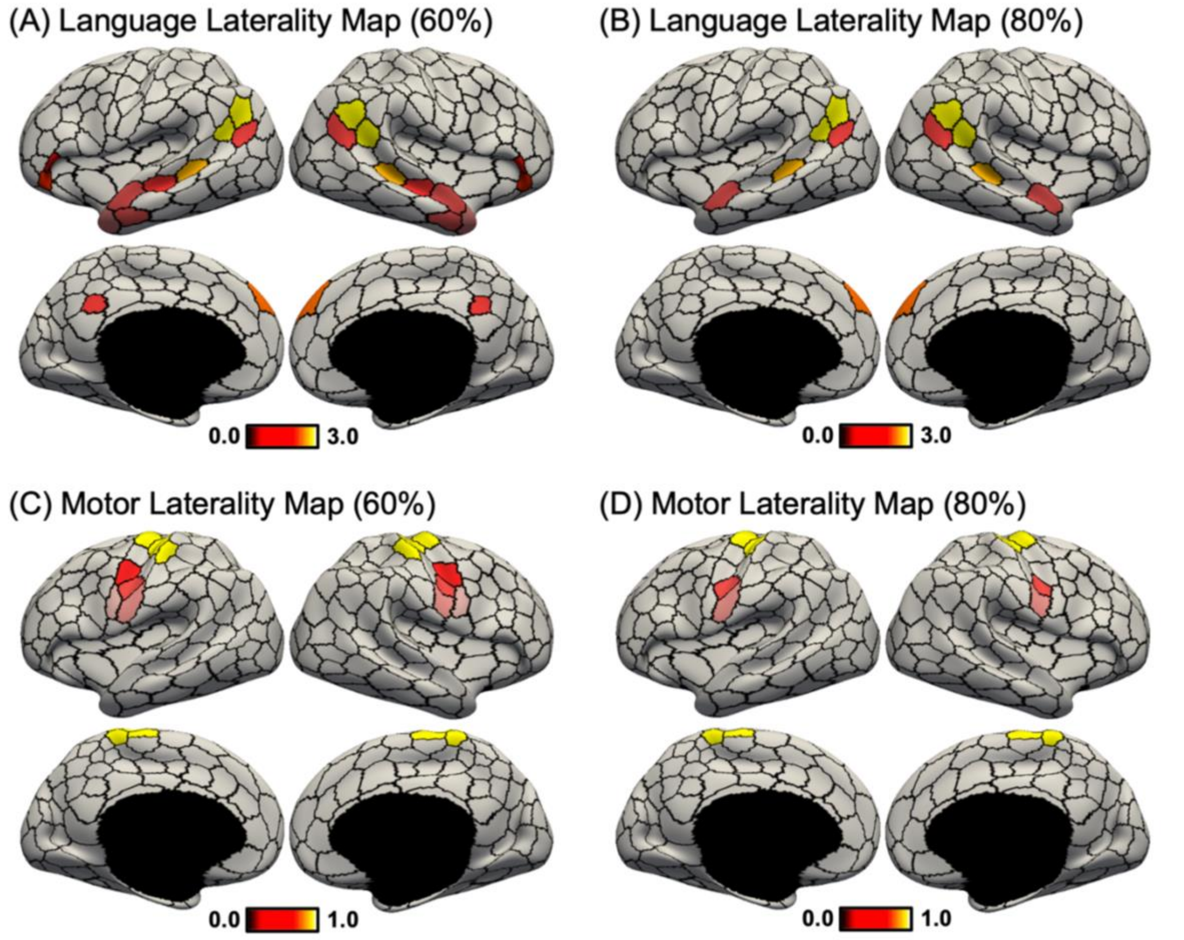


Figure S14. Task activation lateralization maps at different activation thresholds. (A) Task laterality map for the "story -math" language contrast computed for parcels whose average activations were at least 60% of the most activated parcel (number of suprathreshold parcels = 24). (B) Task laterality map for the "story -math" language contrast computed for parcels whose average activations were at least 80% of the most activated parcel (number of suprathreshold parcels = 12). (C) Task laterality map averaged across the motor contrasts, computed for parcels whose average activations were at least 60% of the most activated parcel (number of suprathreshold parcels = 18). (D) Task laterality map averaged across the motor contrasts, computed for parcels whose average activations were at least 80% of the most activated parcel (number of suprathreshold parcels = 12).

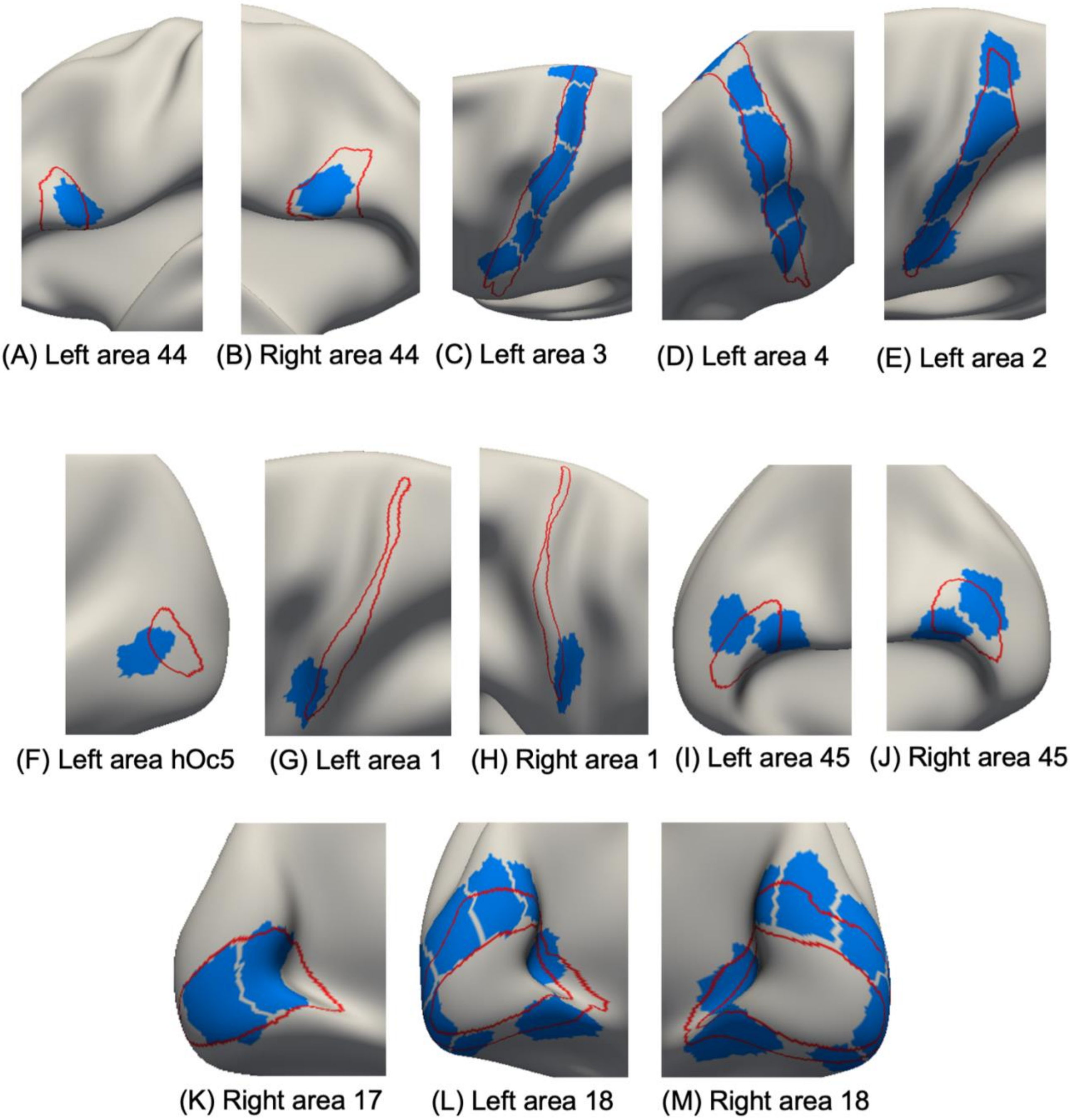


Figure S15. Parcels (blue) of the 400-region hMRF parcellation overlaid on (red) boundaries of histologically defined architectonic areas.

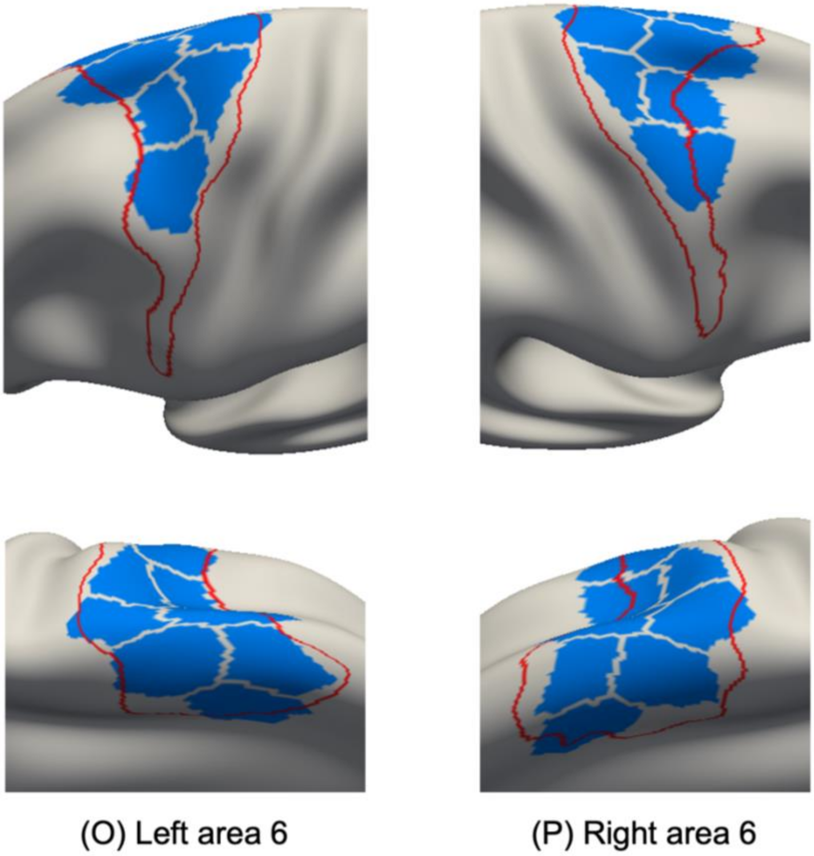


Figure S15 (cont). Parcels (blue) of the 400-region hMRF parcellation overlaid on (red) boundaries of histologically defined architectonic areas.

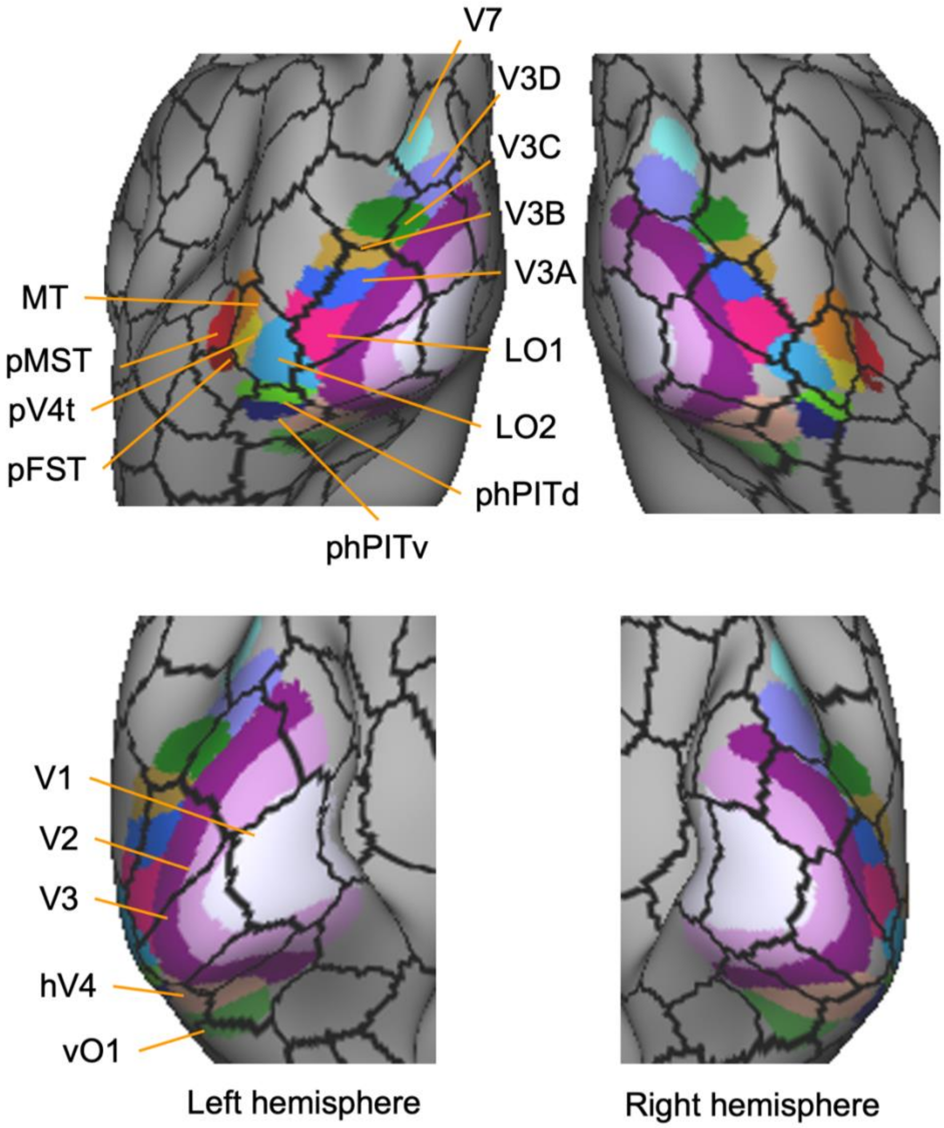


Figure S16. 18 visuotopic areas (Abdollahi et al., 2014) overlaid on (black) boundaries of the 400-region hMRF parcellation.

(A)

| Benchmark parcellation | Benchmark mean homogeneity | hMRF mean homogeneity | % difference | p-value |
|---|---|---|---|---|
| Glasser | 0.0231 | 0.0250 | 8.1% | ≈ 0 |
| Gordon | 0.0257 | 0.0289 | 12.3% | ≈ 0 |
| AICHA | 0.0233 | 0.0245 | 5.0% | ≈ 0 |
| Craddock | 0.0255 | 0.0262 | 3.0% | ≈ 0 |
| Shen | 0.0209 | 0.0221 | 5.5% | ≈ 0 |
| Schaefer | 0.0271 | 0.0271 | 0.16% | ≈ 0 |

(B)

| Benchmark parcellation | Benchmark mean homogeneity | hMRF mean homogeneity | % difference | p-value |
|---|---|---|---|---|
| Glasser | 0.1764 | 0.1892 | 7.3% | 0.0000 |
| Gordon | 0.1959 | 0.2157 | 10.1% | 0.0000 |
| AICHA | 0.1788 | 0.1875 | 4.9% | 0.0000 |
| Craddock | 0.1990 | 0.2013 | 1.2% | 0.0000 |
| Shen | 0.1613 | 0.1621 | 0.50% | 7.7254e-37 |
| Schaefer | 0.2015 | 0.2027 | 0.60% | 0.0000 |

Table S1. Resting-state homogeneity computed using HCP data in MNI152 volumetric space with (A) no spatial smoothing and (B) 6mm FWHM spatial smoothing. The hMRF parcellations exhibited better (higher) resting-state homogeneity than other non-Schaefer parcellations.

| Dataset | AAL mean homogeneity | hMRF mean homogeneity | % difference | p-value |
|---|---|---|---|---|
| ABCD | 0.2707 | 0.3549 | 31.1% | ≈ 0 |
| HCP (fsLR) | 0.0664 | 0.0799 | 20.4% | ≈ 0 |
| GSP | 0.2524 | 0.3286 | 31.1% | ≈ 0 |
| GUSTO | 0.2609 | 0.3437 | 31.7% | ≈ 0 |
| HCP (MNI152) | 0.1030 | 0.1111 | 7.9% | ≈ 0 |

Table S2. Resting-state homogeneity comparison between the AAL (http://www.gin.cnrs.fr/AAL2_files/aal2_for_SPM12.tar.gz) and hMRF parcellations. The resolution of the hMRF parcellation was matched to the AAL parcellation (82 parcels in total) for fair comparison.

| Benchmark parcellation | Benchmark mean homotopic RSFC | hMRF mean homotopic RSFC | % difference | p-value |
|---|---|---|---|---|
| Glasser | 0.1496 | 0.1550 | 3.61% | ≈ 0 |
| AICHA | 0.1540 | 0.1587 | 3.05% | ≈ 0 |

Table S3. Homotopic resting-state functional connectivity using HCP data in MNI152 volumetric space with no spatial smoothing. The hMRF parcellations exhibited better (higher) homotopic resting-state functional connectivity than the AICHA and Glasser parcellations.

| | AICHA | Craddock | Glasser | Gordon | Shen | Schaefer |
|---|---|---|---|---|---|---|
| Benchmark roundness | 0.8957 | 1.1121 | 1.0029 | 0.8544 | 1.0848 | 0.9911 |
| hMRF roundness | 1.0971 | 1.0176 | 1.1541 | 0.6838 | 1.0795 | 1.0677 |
| p value | 4.73e-11 | 6.76e-04 | 3.32e-09 | 6.15e-09 | 0.89 | 1.78e-04 |

Table S4. Comparison of parcel roundness between hMRF and benchmark parcellations with the same number of parcels. The roundness of a parcel is defined as the number of voxels within the parcel (excluding the boundary voxels) divided by the number of boundary voxels within the parcel. A larger roundness metric implies a rounder parcel. The hMRF parcellations were less round than the Gordon and Craddock parcellation, similar in roundness as the Shen parcellation and rounder than the AICHA, Glasser and Schaefer parcellations.

| | AICHA | Craddock | Glasser | Gordon | Shen | Schaefer |
|---|---|---|---|---|---|---|
| Benchmark 90/10 percentile ratio | 7.6083 | 1.7879 | 5.7689 | 8.1975 | 2.8570 | 3.2414 |
| hMRF 90/10 percentile ratio | 3.3160 | 2.8621 | 2.7612 | 3.3319 | 2.8665 | 3.1592 |
| p value | 0.00 | 0.00 | 0.00 | 0.00 | 0.96 | 0.65 |

Table S5. Comparison of parcel size distribution between hMRF and benchmark parcellations with the same number of parcels. Here, parcel size distribution was defined as the volumetric ratio between the 90th percentile parcel to the 10th percentile parcel. A smaller ratio suggests that the parcel size distribution is more uniform. The hMRF parcellations were more uniform than the AICHA, Glasser and Gordon parcellations, less uniform than the Craddock parcellation, and comparable to the Shen and Schaefer parcellations.
